# Deep learning framework ChIANet predicts protein-mediated chromatin architecture across functional contexts

**DOI:** 10.64898/2026.02.24.707640

**Authors:** Hanyu Luo, Renjie Wen, Li Tang, Lingyi Chen, Kailing Tang, Min Li

**Affiliations:** School of Computer Science and Engineering, Central South University, Changsha 410083, China; Department of Genetics, Yale School of Medicine, New Haven, CT 06510, USA

## Abstract

The spatial organization of the genome is dynamically shaped by chromatin-binding proteins, yet how protein-mediated three-dimensional (3D) architectures are specified across functional contexts remains incompletely understood. Here we present ChIANet, a multimodal deep learning framework that enables *de novo* prediction of protein-mediated chromatin contact maps and loops from protein-binding profiles alone, using the reference genome sequence as a prior. By integrating transformer-based long-range modeling with multi-task learning, ChIANet accurately reconstructs protein-mediated 3D chromatin architectures and generalizes across diverse cellular contexts. Systematic application of ChIANet to CTCF, Cohesin and RNAPII across seven human cell types reveals that chromatin architectures follow conserved organizational principles while exhibiting pronounced context-dependent reconfiguration: CTCF- and Cohesin-mediated interactions predominantly support stable structural frameworks, whereas RNAPII-mediated loops display greater variability and are closely coupled to transcriptional programs and regulatory element activity. Functional analyses further uncover distinct regulatory biases and super-enhancer associations among the three proteins. Extending this framework to cancer genomes, ChIANet captures RNAPII-mediated chromatin looping networks associated with extrachromosomal DNA (ecDNA), revealing highly connected, transcription-associated architectures within amplified ecDNA regions across multiple cancer cell types. Together, these results demonstrate that protein-mediated 3D genome organization is not determined by protein identity alone but is flexibly shaped by functional context, regulatory targets and cellular environment, establishing ChIANet as a unified and scalable approach for decoding context-dependent principles of genome folding.

## INTRODUCTION

In eukaryotic cells, chromatin is folded into a highly dynamic three-dimensional (3D) architecture whose organization is hierarchically arranged yet flexibly reconfigured across cellular and regulatory contexts, thereby governing gene expression, transcriptional coordination and cellular identity ^1–3^. This organization emerges from the collective action of DNA-binding and structural proteins that dynamically form, stabilize and remodel chromatin loops ^4,5^. Rather than constituting a uniform structural scaffold, protein-mediated chromatin loops bring distant genomic loci into spatial proximity in a context-dependent manner, enabling enhancer-promoter communication, insulating regulatory domains and supporting diverse gene regulatory programs. Such spatial regulation is fundamental to lineage specification, developmental control and genome stability ^6,7^.

Among key regulators of chromatin architecture, the CCCTC-binding factor (CTCF) functions as a sequence-specific architectural anchor that contributes to the establishment of topologically associating domain (TAD) boundaries ^8,9^. The Cohesin complex, a member of the structural maintenance of chromosomes (SMC) family, cooperates with CTCF to extrude chromatin loops and stabilize higher-order domains, while also participating in regulatory interactions that vary across genomic and cellular contexts ^10,11^. RNA polymerase II (RNAPII), in contrast, mediates transcription-associated chromatin loops by recruiting Mediator and transcription factors, forming a dynamic regulatory layer that is closely coupled to gene activity and regulatory element usage ^12,13^. These protein-mediated interactions are further shaped by chromatin states, including histone modifications such as H3K27ac and H3K27me3, which mark active and repressive regulatory regions and modulate the accessibility, stability and functional outcome of chromatin contacts ^14,15^.

Chromatin conformation capture (3C) technologies and their derivatives, including Hi-C, have enabled genome-wide mapping of chromatin contact frequencies and revealed general principles of 3D genome organization ^16–18^. Targeted assays such as ChIA-PET ^19^, HiChIP ^20^, and PLAC-seq ^21^ further integrate chromatin immunoprecipitation with proximity ligation to profile interactions associated with specific proteins or histone modifications. However, these experimental approaches remain cost-intensive and laborious, requiring large cell numbers, deep sequencing and high-quality antibodies for each target protein ^2^. Their limited scalability across proteins and cell types has constrained systematic investigation of how protein-mediated chromatin architectures vary across functional contexts, leaving a substantial gap in our understanding of context-dependent 3D genome regulation.

To overcome these limitations, computational modeling has emerged as a powerful approach for predicting chromatin interactions ^22,23^. Deep learning-based models such as Akita ^24^, Orca ^25^, and C.Origami ^26^ have achieved impressive accuracy in reconstructing Hi-C contact maps from DNA sequence or epigenomic signals ^27,28^. Other frameworks, including Peakachu ^29^ and DeepChIA-PET ^30^, predict loops from experimental Hi-C or ChIA-PET data. In parallel, another class of models formulates loop detection as an anchor-pair classification problem, in which two genomic loci are provided as input and the model predicts whether they form a loop interaction ^31–33^. Despite these advances, most existing approaches remain limited to data-dependent or protein-agnostic settings, focusing primarily on generic architectural features rather than protein-mediated regulatory mechanisms. As a result, they provide limited insight into how distinct chromatin-binding proteins organize three-dimensional genome architecture across different functional contexts or cell types. More recently, several protein-specific predictors—for example, models targeting CTCF- or YY1-mediated loops—have been developed to capture motif-driven interactions ^34,35^. However, these methods are narrowly tailored to individual proteins and specific regulatory scenarios, lacking extensibility to diverse chromatin-associated factors or broader cellular environments. Consequently, a unified computational framework capable of de novo prediction of diverse protein-mediated chromatin architectures in a context-aware manner across cell types remains lacking.

To address this gap, we developed ChIANet, a protein-specific multimodal deep learning framework that integrates genomic sequence with protein-specific ChIP-seq profiles to predict chromatin contact maps and loops mediated by distinct chromatin regulators across functional contexts. ChIANet couples a Transformer-based encoder, which captures long-range genomic dependencies from sequence and binding signals, with a multi-task decoder that jointly models contact intensity and loop probability, thereby learning unified representations of structural organization and regulatory activity. Methodologically, ChIANet combines heteroscedastic uncertainty-weighted multi-task learning with genome-wide tiling, enabling stable training, consistent resolution across genomic scales and efficient application to entire chromosomes. Unlike previous factor-specific or Hi-C-dependent models, ChIANet supports de novo reconstruction of protein-mediated three-dimensional genome architecture in unseen cell types using only ChIP-seq profiles as experimental input, without requiring Hi-C supervision or model retraining. By systematically applying ChIANet to three representative chromatin regulators—CTCF, Cohesin and RNAPII—across seven human cell types, we reveal that protein-mediated chromatin architectures follow conserved organizational rules while being flexibly reconfigured across regulatory and cellular contexts. We further demonstrate that this framework extends to cancer genomes, capturing RNAPII-mediated chromatin looping architectures associated with extrachromosomal DNA (ecDNA), which represents an extreme regulatory context characterized by copy-number amplification and dense regulatory activity.

## RESULTS

### ChIANet enables accurate and generalizable prediction of protein-mediated chromatin interactions

We developed ChIANet, a multimodal encoder-decoder framework that integrates the reference genome sequence with protein-specific ChIP-seq profiles to predict protein-mediated chromatin organization at genome scale (Fig. 1a, b; Supplementary Fig. 1). Within each 2.1-Mb window, local sequence and binding features are processed by stacked Conv1D residual blocks and an eight-layer Transformer encoder to capture long-range dependencies ^36^, followed by Conv2D decoding to jointly generate contact maps and loop matrices under a heteroscedastic uncertainty-weighted multi-task loss ^37^(Methods).

**Fig. 1:**
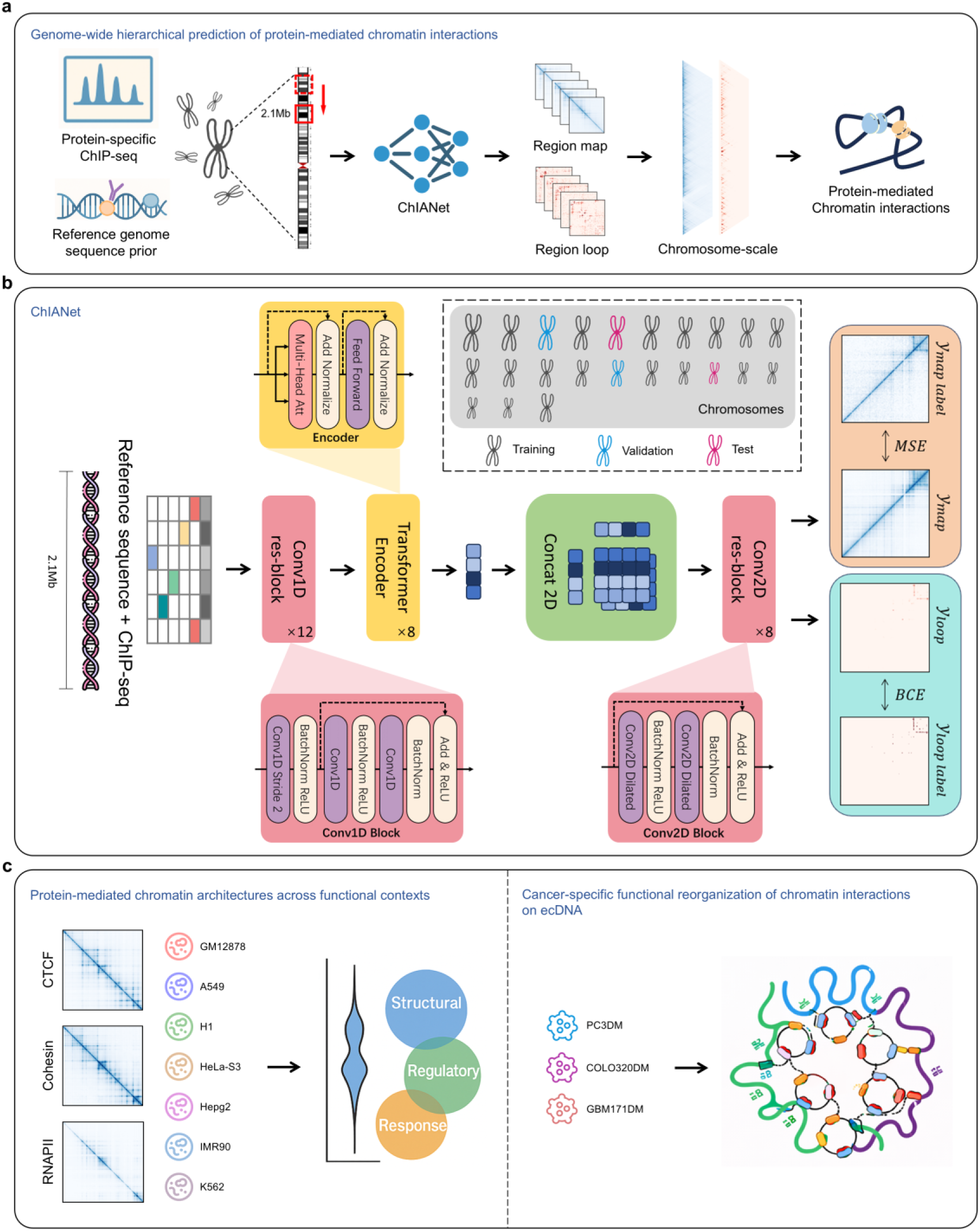
Overview of the ChIANet architecture and workflow. **(a)** Protein-specific ChIP-seq profiles, together with the reference genome sequence, are used as inputs to predict protein-mediated chromatin interactions. ChIANet generates contact maps and loop for each 2.1-Mb window, which are tiled along chromosomes and merged to obtain genome-scale predictions. **(b)** ChIANet is a multimodal, multi-task encoder-decoder framework that integrates genomic sequence and protein-specific ChIP-seq signals to predict protein-mediated chromatin organization. ChIANet jointly outputs a contact map and loop matrix under a heteroscedastic uncertainty-weighted loss. Models are trained with a 19/2/2 chromosome split (train/validation/test) to ensure unbiased evaluation. The trained model supports de novo inference in unseen cell types from ChIP-seq input alone. **(c)** Predicted chromatin interaction maps for CTCF, Cohesin and RNAPII across seven human cell types enable comparative analyses of protein-mediated chromatin architectures across functional contexts. The same framework is applied to cancer genomes to analyze RNAPII-mediated chromatin interactions associated with extrachromosomal DNA (ecDNA).

For each chromatin regulator (CTCF, Cohesin and RNAPII), we trained a dedicated model using GM12878 ChIA-PET data with a chromosome-level split (19/2/2 for training, validation and testing), ensuring strict separation of genomic contexts during evaluation (Fig. 1b). Genome-wide predictions were obtained by tiling overlapping windows and stitching regional outputs into continuous chromosomal maps (Methods). Because ChIANet conditions only on protein-specific ChIP-seq signals and the invariant genome sequence, trained models can be directly applied to unseen cell types without retraining. Using this strategy, we generated de novo protein-mediated contact maps and loop predictions across seven human cell types, establishing a unified framework for cross-context comparative analyses (Fig. 1c).

We next systematically evaluated ChIANet’s performance in reconstructing protein-mediated contact maps. As no existing method explicitly models protein-specific interactions, we compared ChIANet with three state-of-the-art Hi-C prediction models—Akita ^24^, Orca ^25^, and C.Origami ^26^(Methods)—under identical preprocessing and evaluation settings (Methods). Across all test-set genomic windows, ChIANet consistently achieved higher Pearson and Spearman correlations and lower mean squared error compared with competing methods. (Fig. 2a, Extended Data Fig. 1a). Importantly, ChIANet maintained stronger correlations at increasing genomic distances, indicating improved modeling of long-range protein-mediated contacts (Fig. 2b, Extended Data Fig. 1b). Chromosome-scale scatter analyses further confirmed the high concordance between predicted and experimental contact intensities across all proteins (Supplementary Fig. 3). Representative regions further illustrate the accurate reconstruction of both local and distal interaction patterns observed in ChIA-PET data (Fig. 2c, d, h; Extended Data Fig. 3).

**Fig. 2:**
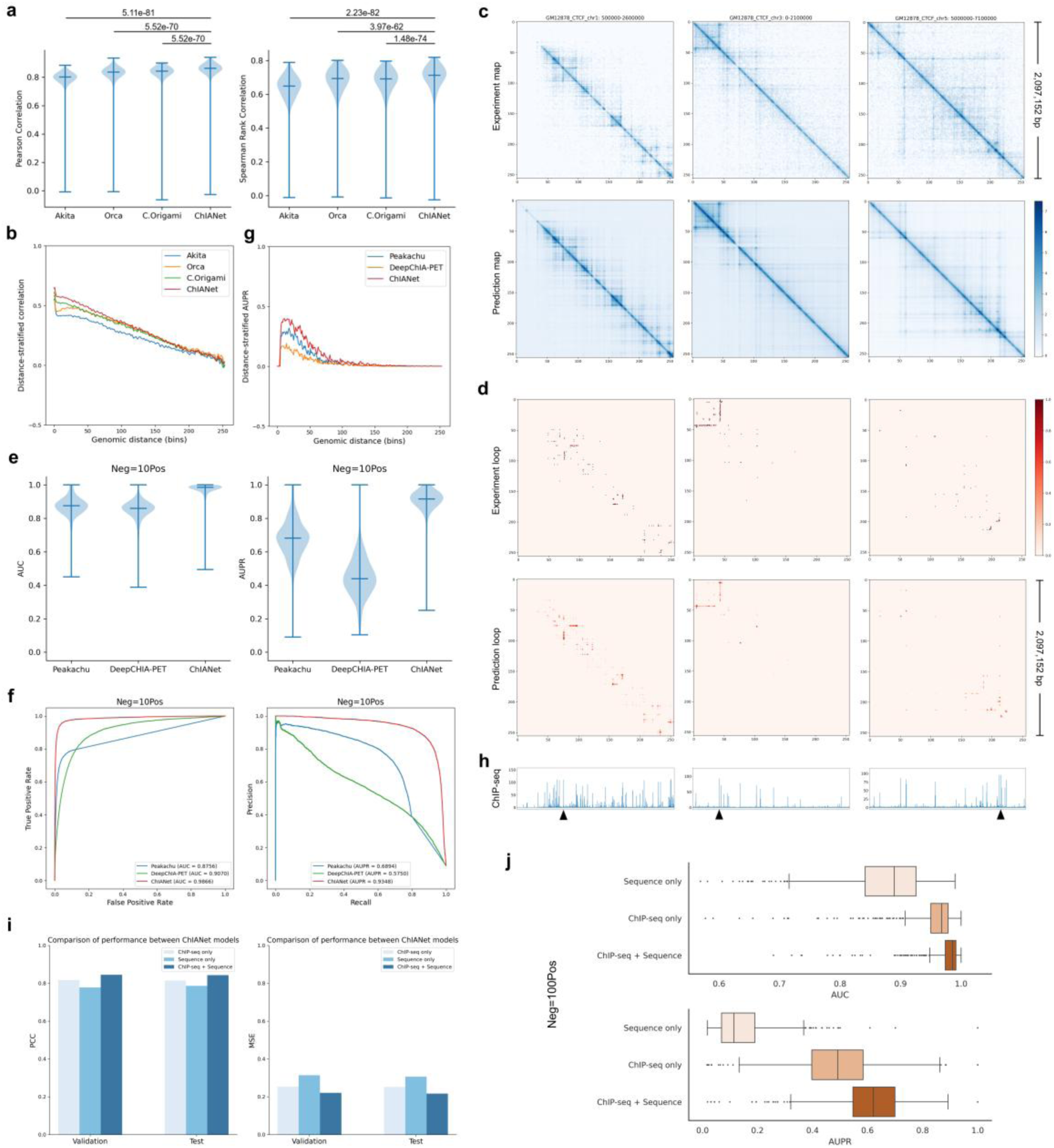
ChIANet accurately predicts protein-mediated contact maps and loop interactions. **(a)** Model performance comparison on GM12878 CTCF test sets. Pearson (left) and Spearman (right) correlations between predicted and experimental contact maps across all 2.1-Mb windows, benchmarked against Akita, Orca and C.Origami. Violin plots show the distribution across genomic windows; horizontal lines indicate minimum, mean and maximum values. Statistical significance was assessed using paired two-sided Wilcoxon signed-rank tests. **(b)** Distance-stratified Pearson correlations between predicted and experimental contact maps for different models. **c-d,** Representative examples of experimental and model-predicted results for GM12878 CTCF. (c) Contact maps and (d) loops shown for training (chr1), validation (chr3), and test (chr5) genomic regions **(e)** Genome-wide comparison of AUC and AUPR for loop prediction (positive-to-negative ratio = 1:10) among Peakachu, DeepChIA-PET and ChIANet. **(f)** Chromosome-scale receiver operating characteristic (ROC; left) and precision-recall (PR; right) curves showing overall AUC and AUPR on the GM12878 CTCF test set. **(g)** Distance-stratified area under the precision-recall curve (AUPR) for loop prediction. **(h)** Protein-specific ChIP-seq input profiles corresponding to the same regions. **(i)** Comparison of model performance across different input modalities—ChIP-seq only, sequence only, and sequence + ChIP-seq (full model)—on the contact map prediction task. Bars show Pearson correlation coefficients (PCC, left) and mean squared error (MSE, right) on the validation and test sets. **(j)** Comparison of window-level AUC and AUPR for the loop prediction task under an imbalanced classification ratio (1:100, Neg:Pos).

We further assessed loop prediction performance by comparing ChIANet with Peakachu ^29^ and DeepChIA-PET ^30^, the two genome-wide frameworks capable of direct loop inference (Methods). ChIANet consistently achieved the highest AUC and AUPR at both window and chromosome scales (Fig. 2e, f; Extended Data Fig. 2a, b), and maintained robust performance under increasingly imbalanced classification settings (Extended Data Fig. 1d). Notably, whereas both baseline methods require Hi-C contact maps as input, ChIANet relies solely on sequence and protein-specific ChIP-seq profiles, demonstrating that accurate loop prediction can be achieved without Hi-C supervision. The model also achieved the highest distance-stratified AUPR, reflecting its capacity to capture both proximal and distal looping patterns (Fig. 2g; Extended Data Fig. 1c, Extended Data Fig. 4).

Together, these results establish ChIANet as an accurate and generalizable framework for predicting protein-mediated chromatin interactions, providing a robust foundation for downstream analyses of protein-specific regulatory architectures across functional contexts.

### Multimodal integration improves protein-specific interaction prediction

While CTCF binds to well-defined DNA motifs, other chromatin regulators such as Cohesin and RNAPII display weaker sequence specificity and encode their chromatin interactions more strongly through context-dependent chromatin features ^8^. To dissect the relative contributions of sequence and ChIP-seq features in protein-mediated interaction prediction, we conducted ablation experiments by training two ChIANet variants with identical architectures but different inputs: a sequence-only model and a ChIP-seq-only model. Across all three proteins in GM12878, the full sequence + ChIP-seq model (original ChIANet) achieved the lowest convergence loss for both contact map and loop prediction tasks (Extended Data Fig. 5a). Notably, the sequence-only model for CTCF converged more stably, whereas those for Cohesin and RNAPII showed pronounced fluctuations during training, reflecting the limited ability of sequence information alone to capture interaction patterns of proteins that are less motif-driven and more context-dependent.

For the contact map prediction task, the full model consistently achieved the highest Pearson correlation coefficients (PCC) and the lowest mean squared errors (MSE) across all proteins (Fig. 2i; Extended Data Fig. 5b). In distance-stratified correlation analyses, ChIANet also showed the strongest performance (Extended Data Fig. 5c). In contrast, the sequence-only model rapidly lost correlation beyond 50 bins, whereas the ChIP-seq-only model maintained moderate performance, highlighting the necessity of ChIP-seq information for modeling protein-dependent long-range interactions. For the loop prediction task, a similar trend was observed (Extended Data Fig. 5d). Even under increasingly imbalanced classification ratios, the full model maintained robust AUPR values across all proteins (Fig. 2j; Supplementary Fig. 4). At the chromosome scale (1:100 negative-to-positive ratio), ChIANet reached an AUPR of 0.683 for CTCF—representing improvements of +0.536 and +0.196 over the sequence-only and ChIP-seq-only models, respectively—and 0.364 for RNAPII (+0.226 and +0.068), and 0.670 for Cohesin (+0.523 and +0.140) (Extended Data Fig. 5e).

To visualize how multimodal integration enhances prediction fidelity, we examined representative genomic regions. As shown in Supplementary Fig. 5, the full ChIANet model accurately reconstructed clear protein-specific topological architectures and high-confidence loops consistent with ChIA-PET data. In contrast, the ChIP-seq-only model produced blurred topological domains with weakened distal contacts, while the sequence-only model failed to recover any discernible structure or loop signal. Together, these analyses demonstrate that integrating nucleotide sequence with protein-specific ChIP-seq profiles enables ChIANet to capture distinct informational regimes underlying protein-mediated chromatin architectures, spanning motif-driven and context-dependent regulatory interactions.

### *De novo* prediction of protein-mediated chromatin organization across cellular contexts

Having established the modality advantage of ChIANet, we next evaluated its ability to generalize across cell types. We trained a CTCF-specific model exclusively on GM12878 cells and performed *de novo* predictions of CTCF-mediated chromatin architecture in H1 cells without any retraining. Remarkably, ChIANet faithfully reconstructed both local and long-range contact patterns that closely mirrored the experimental ChIA-PET maps in H1 (Fig. 3a; Extended Data Figs. 3, 4). The predicted contact maps and loop structures captured pronounced chromatin rearrangements between GM12878 and H1, consistent with differential CTCF occupancy along the genome.

**Fig. 3:**
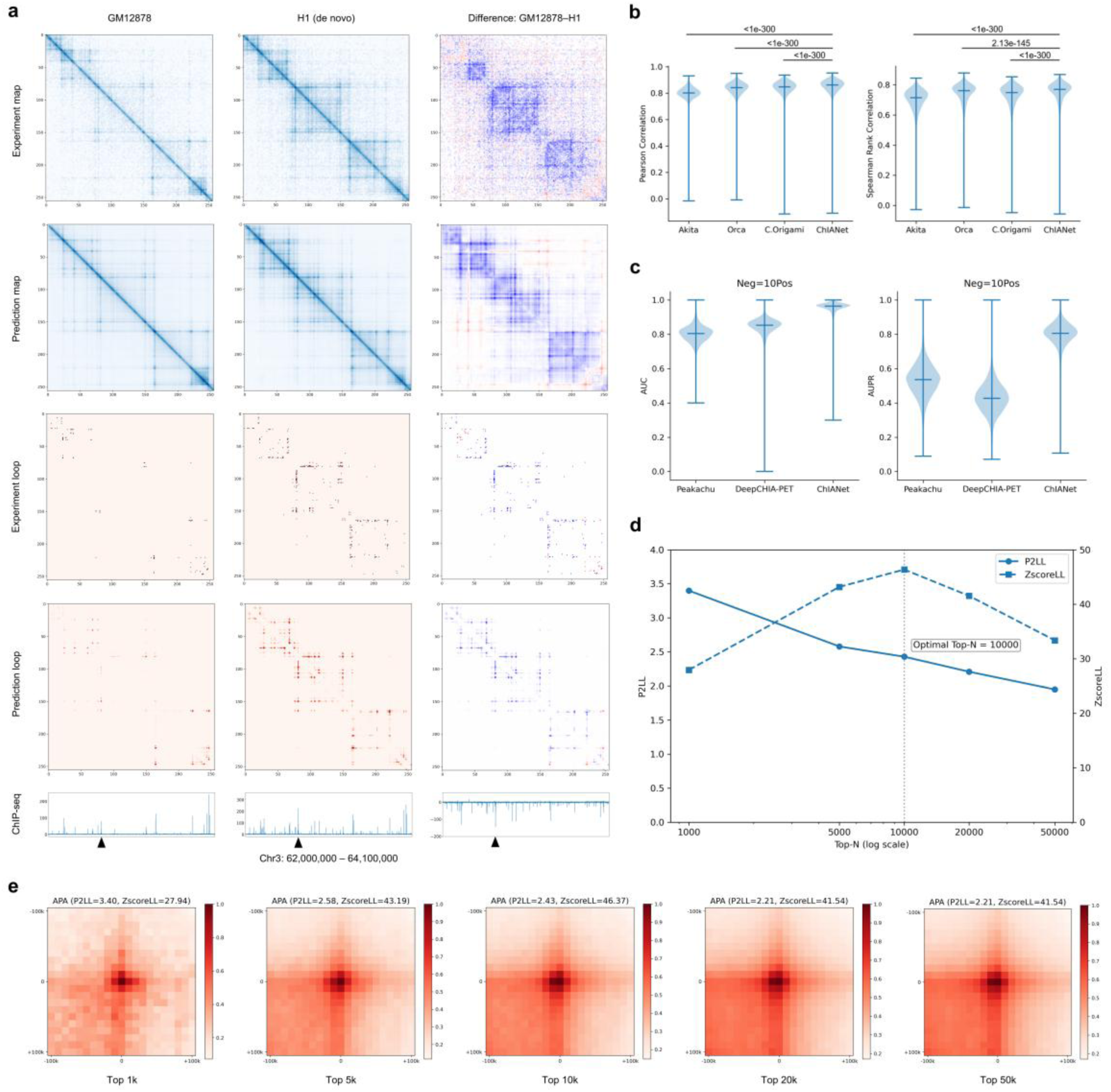
*De novo* prediction of cell type-specific, protein-mediated chromatin structure. **(a)** Experimental and predicted CTCF-mediated contact maps and loop matrices for GM12878 (left) and H1 (center), together with their difference maps (GM12878 - H1, right). Blue/red in the difference panels indicate contacts or loops that are stronger in GM12878 or H1, respectively. The bottom track shows the CTCF ChIP-seq signal used for conditioning. **(b)** Genome-wide comparison of contact map prediction performance on H1 CTCF. Violin plots show Pearson correlation (left) and Spearman rank correlation (right) between predicted and experimental 2.1-Mb windows (n = 5,977), with ChIANet achieving the highest median concordance. Statistical significance was assessed using paired two-sided Wilcoxon signed-rank tests. **(c)** Genome-wide comparison of loop prediction on H1 CTCF at a 1:10 positive-to-negative ratio. Violin plots show window-level AUC (left) and AUPR (right), indicating that ChIANet maintains competitive loop-level accuracy in *de novo* prediction. **(d)** APA quality metrics for *de novo* predicted H1 CTCF loops as a function of the number of top-ranked loops retained. P2LL (solid line) and ZscoreLL (dashed line) were computed on loop sets of increasing size (Top 1k to Top 50k). Both metrics peak around Top 10k, indicating that this cutoff yields the best trade-off between loop strength and noise. **(e)** APA heatmaps of *de novo* predicted H1 loops at different cutoffs (Top 1k, 5k, 10k, 20k and 50k), shown in a ±100-kb window around the loop anchor pair.

For the contact map prediction task, ChIANet achieved comparable or higher correlations than sequence-based and multimodal Hi-C prediction models across all genomic windows, with median Pearson and Spearman correlations exceeding 0.8 (Fig. 3b; Extended Data Fig. 6). For the loop prediction task, ChIANet also exhibited strong transferability, yielding the highest AUC and AUPR across a wide range of positive-to-negative ratios (1:1 to 1:100) compared with Peakachu and DeepChIA-PET (Fig. 3c; Extended Data Fig. 7; Supplementary Fig. 6). Notably, while both baseline models require Hi-C contact maps as input features, ChIANet relies solely on DNA sequence and protein-specific ChIP-seq tracks, underscoring its capacity to generalize to unseen cellular contexts without Hi-C guidance.

To further quantify global *de novo* loop prediction accuracy, we performed Aggregate Peak Analysis (APA) on the top-scoring predicted loops ^38^ (Fig. 3d, e; Methods). As the number of predicted loops increased from 1k to 50k, the enrichment metrics—P2LL and ZscoreLL—initially rose and peaked around 10,000 loops, indicating an optimal balance between precision and coverage (Fig. 3d). APA heatmaps showed strong focal enrichment centered at predicted loop anchors across all thresholds, consistent with interaction patterns observed in experimental Hi-C maps (Fig. 3e).

Overall, these results demonstrate that ChIANet can generalize across cell types to accurately reconstruct both contact maps and loop-level features of protein-mediated chromatin interactions. By integrating DNA sequence and protein-specific ChIP-seq information, ChIANet captures transferable organizational rules of protein-mediated chromatin architecture, enabling high-fidelity *de novo* prediction across distinct cellular contexts.

### Protein-specific yet coordinated chromatin architectures are conserved across cell types

To examine how distinct chromatin regulators jointly organize three-dimensional genome architecture across cellular contexts, we applied ChIANet to predict protein-mediated contact maps and loops for CTCF, Cohesin and RNAPII across seven representative human cell types (GM12878, H1, A549, HeLa-S3, IMR90, HepG2 and K562).

Using H1 as an illustrative example, the predicted contact maps revealed distinct yet coordinated organizational patterns among the three proteins (Fig. 4a). CTCF- and Cohesin-mediated maps exhibited highly concordant domain-level structures, whereas RNAPII-mediated interactions displayed more spatially diffuse patterns preferentially enriched at transcriptionally active regions. Genome-wide pairwise correlations confirmed this relationship, with consistently higher concordance between CTCF and Cohesin compared with protein pairs involving RNAPII (Fig. 4b; Extended Data Fig. 8a). Importantly, global UMAP embedding of all 2.1-Mb windows demonstrated that while CTCF and Cohesin occupied overlapping manifolds, RNAPII maps formed a partially separated yet adjacent structural space, indicating that transcription-associated interactions are organized in close coordination with, rather than independently from, the architectural backbone (Fig. 4c; Extended Data Fig. 8c). These relationships were preserved across individual chromosomes (Fig. 4d; Extended Data Fig. 8b) and across genomic distance scales, with stronger long-range concordance for CTCF-Cohesin pairs (Fig. 4e; Extended Data Fig. 9a). Together, these results indicate that architectural proteins (CTCF/Cohesin) cooperatively maintain domain-level structure, whereas RNAPII contributes a transcription-associated layer.

**Fig. 4:**
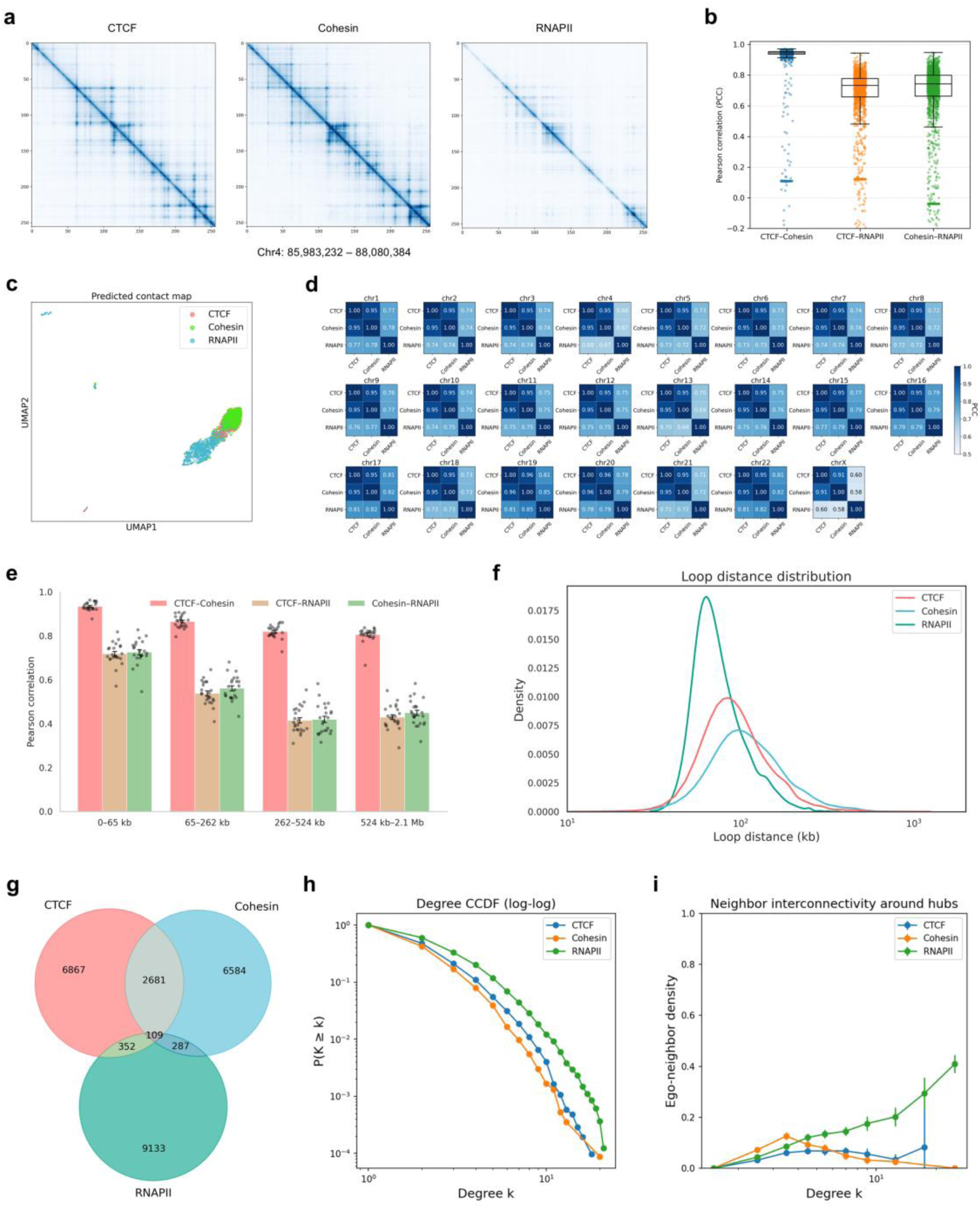
Distinct 3D chromatin architectures mediated by different regulatory proteins in H1. **(a)** Representative predicted contact maps for CTCF-, Cohesin-, and RNAPII-mediated chromatin interactions within the same genomic region (Chr4: 85.98-88.08 Mb). CTCF and Cohesin maps exhibit sharply defined domain boundaries, whereas RNAPII shows diffuse, transcription-associated interaction patterns. **(b)** Genome-wide pairwise Pearson correlation of predicted contact maps across all 2.1-Mb windows, illustrating strong similarity between CTCF and Cohesin, and weaker correlation between RNAPII and the other two factors. **(c)** UMAP embedding of all predicted contact maps across the genome, showing substantial overlap between CTCF and Cohesin clusters, while RNAPII forms a distinct group, reflecting a divergent interaction landscape. **(d)** Chromosome-wise Pearson correlations between predicted maps of different proteins, with the highest values consistently observed between CTCF and Cohesin, indicating conserved structural coupling across chromosomes. **(e)** Distance-stratified correlations between predicted contact maps of different proteins. Correlations between CTCF-RNAPII or Cohesin-RNAPII drop markedly at long genomic distances (>524 kb), whereas CTCF-Cohesin correlations remain stable, indicating their shared role in maintaining higher-order domain organization. **(f)** Distribution of loop lengths for the top 10,000 predicted loops of each protein. RNAPII-mediated loops are generally shorter, consistent with promoter-enhancer proximity, whereas CTCF and Cohesin loops span broader genomic ranges. **(g)** Venn diagram showing the overlap of top predicted loops among the three proteins, highlighting substantial sharing between CTCF and Cohesin. **(h)** Complementary cumulative degree distribution (CCDF) of loop networks for each protein, revealing scale-free behavior with RNAPII networks showing higher connectivity at low degrees and CTCF/Cohesin networks forming stronger high-degree hubs. **(i)** Neighbor interconnectivity as a function of loop hub degree. CTCF and Cohesin display greater local clustering around high-degree nodes than RNAPII.

Loop-level analyses further highlighted both shared and protein-specific organizational principles. RNAPII-mediated loops were consistently shorter in genomic span, whereas CTCF- and Cohesin-mediated loops extended over broader distances (Fig. 4f; Extended Data Fig. 9b). Substantial overlap was observed between CTCF and Cohesin loop sets, whereas RNAPII loops showed limited intersection, consistent with their preferential engagement in localized regulatory interactions (Fig. 4g; Extended Data Fig. 9c). Despite these differences, network topology analyses revealed convergent global properties across all three proteins, including scale-free degree distributions indicative of hub-centered organization (Fig. 4h; Extended Data Fig. 10a). Across cell types, CTCF and Cohesin networks were enriched for high-degree hubs that contributed disproportionately to long-range connectivity, whereas RNAPII networks were characterized by smaller k-core structures and locally clustered interaction communities (Fig. 4i; Extended Data Fig. 10b-d), reflecting a more dynamic and context-responsive mode of chromatin organization.

Collectively, these analyses reveal a conserved yet flexible organizational framework in which architectural proteins and transcription-associated regulators operate in a coordinated manner across diverse cellular contexts. Rather than forming isolated structural layers, CTCF-, Cohesin- and RNAPII-mediated interactions collectively shape chromatin architecture through context-dependent reweighting of shared organizational principles, providing a mechanistic basis for both structural stability and regulatory plasticity across cell types.

### Cross-cell-type conservation and variability of protein-mediated chromatin architecture

Building on the coordinated yet protein-specific organizational patterns observed within individual cell types, we next examined how protein-mediated chromatin architectures are preserved or reconfigured across distinct cellular contexts. Genome-wide correlations between predicted contact maps revealed that CTCF- and Cohesin-mediated interactions display substantially higher cross-cell-type concordance than RNAPII-mediated interactions (Fig. 5a), indicating that architectural features associated with these proteins are more consistently maintained across cellular states, whereas transcription-associated contacts exhibit pronounced context dependence. To quantify this behavior at higher resolution, we computed a conservation score for each genomic window across the seven cell types (Methods). Chromosome-resolved analyses confirmed a clear stratification, with CTCF and Cohesin exhibiting uniformly high conservation and RNAPII showing markedly greater dispersion (Supplementary Fig. 9a). Visualization of these conservation landscapes along individual chromosomes further revealed extended domains of stable CTCF- and Cohesin-mediated organization interspersed with regions in which RNAPII-mediated interactions varied substantially between cell types (Supplementary Fig. 9b). Representative loci were selected to illustrate conserved versus context-variable architectural patterns (Fig. 5b).

**Fig. 5:**
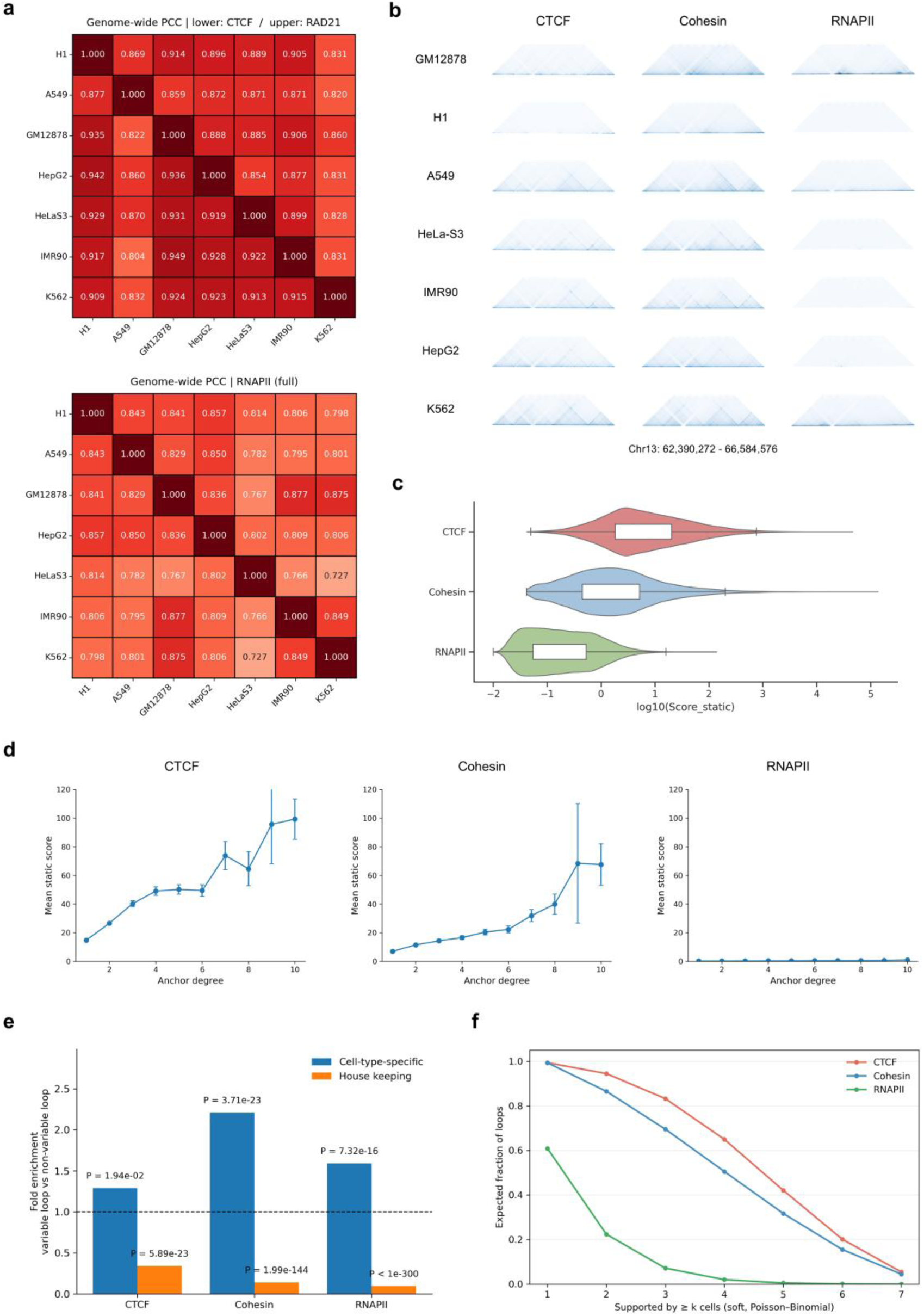
Cross-cell-type conservation and variability of protein-mediated chromatin organization. **(a)** Genome-wide Pearson correlation coefficients (PCCs) between predicted contact maps across seven human cell types for each protein. The lower triangle of the upper heatmap shows CTCF-mediated correlations, the upper triangle shows Cohesin, and the bottom panel shows RNAPII. **(b)** Representative regions on chromosome 13 (62.39-66.58 Mb) illustrating cross-cell-type differences in contact maps for CTCF, Cohesin, and RNAPII. **(c)** Distribution of loop static scores quantifying the cross-cell-type conservation of predicted loops for each protein. **(d)** Relationship between node degree and mean loop stability across protein-specific loop networks. **(e)** Enrichment analysis comparing variable and stable loops near cell type-specific versus housekeeping genes (P values, Fisher’s exact test). **(f)** Fraction of predicted loops supported across increasing numbers of cell types, estimated by a Poisson-Binomial model.

We next extended this analysis to the loop level. For each protein, a pan-cell set of predicted loops was assembled, and loop conservation across cell types was quantified using a static score. In total, 34,751 CTCF, 48,847 Cohesin and 46,093 RNAPII loops were identified. Based on their static scores, loops were stratified into stable and variable categories (Supplementary Fig. 10). The resulting distributions revealed a hierarchical pattern of conservation, with CTCF loops exhibiting the greatest stability, followed by Cohesin and RNAPII (Fig. 5c), consistent with their distinct organizational roles. At the network level, loop stability was strongly associated with anchor connectivity. Loops anchored at high-degree nodes were significantly more stable across cell types, particularly for CTCF and Cohesin (Fig. 5d), suggesting that highly connected hubs serve as conserved organizational scaffolds. In contrast, lower-degree RNAPII loops exhibited greater variability, reflecting their preferential engagement in cell-type-specific regulatory interactions. To link loop conservation with transcriptional programs, genes were classified as housekeeping or cell-type-specific based on RNA-seq expression profiles. Variable loops were significantly enriched in the vicinity of cell-type-specific genes, whereas stable loops preferentially associated with housekeeping genes (Fig. 5e), indicating that context-dependent rewiring of chromatin loops underlies transcriptional diversification across cell types.

Finally, loop conservation was assessed using a probabilistic support metric that quantifies the number of cell types in which each loop is detected. CTCF and Cohesin loops were supported by substantially more cell types than RNAPII loops (Fig. 5f), with more than half of CTCF and Cohesin loops retained in at least four cell types, whereas the majority of RNAPII loops were restricted to one or two. Together, these analyses demonstrate that protein-mediated chromatin architecture is governed by a balance between conserved structural scaffolding and context-dependent regulatory reconfiguration, with different proteins contributing distinctially to architectural stability and transcriptional plasticity across cellular contexts.

### Protein-mediated chromatin loops exhibit context-dependent functional biases and regulatory potential

We next examined how protein-mediated chromatin loops are functionally deployed across genomic contexts and how their regulatory potential varies with protein identity and interaction topology. At the genome-wide scale, CTCF- and Cohesin-mediated loops were preferentially associated with chromatin domain boundaries, whereas RNAPII-mediated loops were enriched at regulatory elements, including promoters and enhancers (Fig. 6a; Extended Data Fig. 11a). Importantly, the degree of functional enrichment scaled with anchor connectivity: highly connected CTCF and Cohesin anchors showed progressively stronger association with domain boundaries, whereas increasing RNAPII connectivity was accompanied by a pronounced shift toward promoter-linked interactions, indicating context-dependent reweighting of loop function as network complexity increases. Classification of pan-cell-type loops further revealed distinct functional interaction classes associated with different proteins. CTCF-mediated loops were enriched for structural configurations, including a substantial fraction of silencer-promoter interactions, whereas RNAPII-mediated loops were dominated by promoter-promoter and enhancer-promoter interactions (Fig. 6b). These functional compositions were consistently observed across all additional cell types analyzed (Extended Data Fig. 11b), indicating that protein-specific interaction biases represent conserved organizational tendencies rather than cell-type-specific artifacts.

**Fig. 6:**
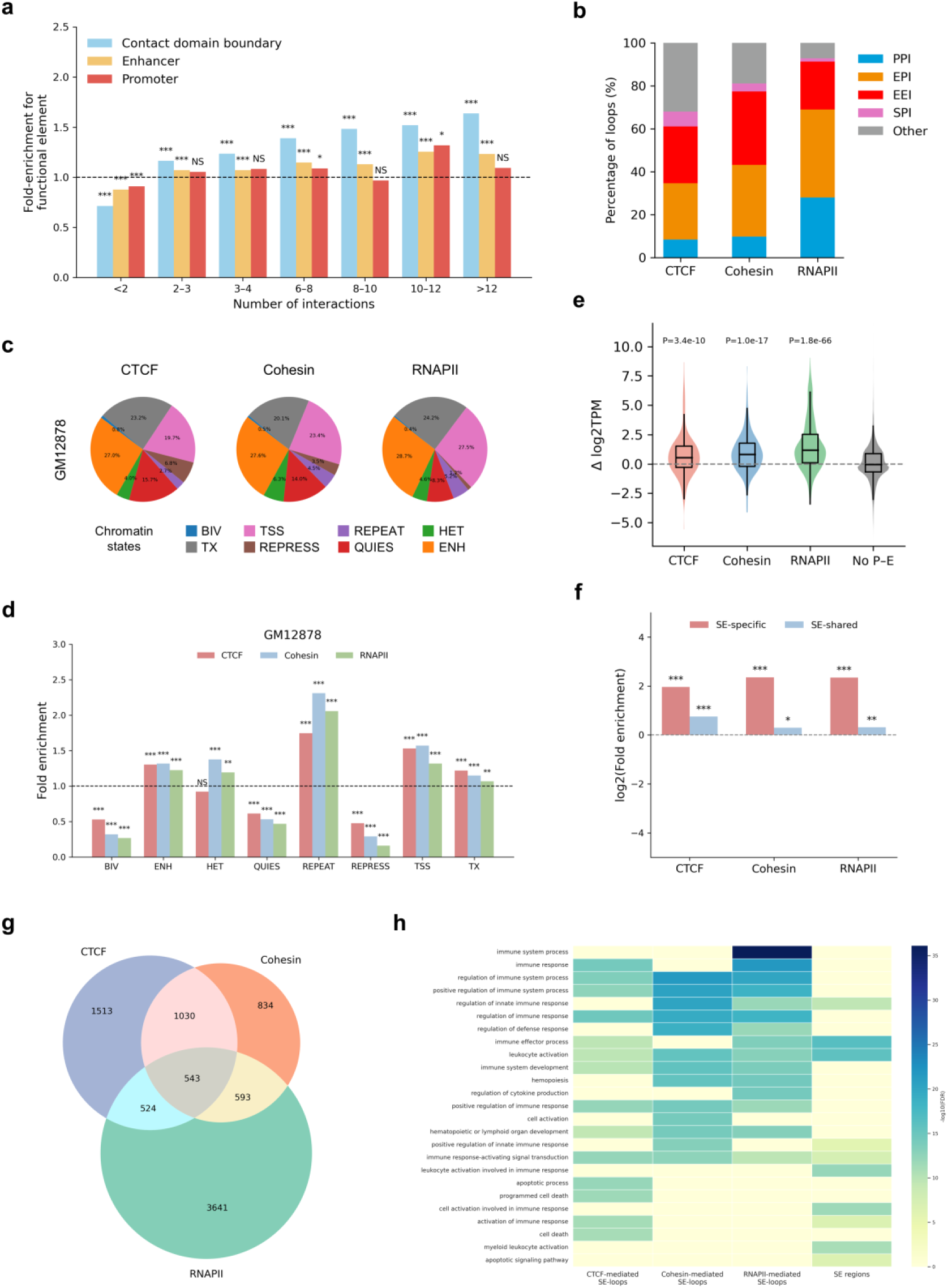
Functional bias and regulatory potential of protein-mediated chromatin interactions. **(a)** Functional enrichment of anchors from pan-cell-type CTCF-mediated loops for contact domain boundaries, enhancers, and promoters, stratified by the number of loop interactions per anchor. Statistical significance was assessed using two-sided Fisher’s exact tests (Number of interactions = 34,751, \**P<*0.05, \*\**P<*0.01, \*\*\**P<*0.001; *NS*, not significant). **(b)** Classification of loops mediated by CTCF, Cohesin, and RNAPII across all cell types into promoter-promoter interactions (PPI), enhancer-promoter interactions (EPI), enhancer-enhancer interactions (EEI), silencer-promoter interactions (SPI), and other types. **(c)** Chromatin-state composition of cell-type-specific loop anchors in GM12878. **(d)** Fold enrichment of chromatin states for cell-type-specific loop anchors mediated by CTCF, Cohesin, and RNAPII in GM12878 (n = 2,149, 2,626, and 2,033, respectively). Enrichment significance was evaluated using two-sided Fisher’s exact tests. **(e)** Differential gene expression associated with genes linked to cell-type-specific loops in GM12878. P-values were calculated by two-sided Wilcoxon rank-sum tests. **(f)** Enrichment of GM12878-specific loops relative to non-specific loops at cell-type-specific super-enhancers (SE-specific) and shared super-enhancers (SE-shared), with significance assessed by Fisher’s exact test. **(g)** Overlap of genes associated with CTCF-, Cohesin-, and RNAPII-mediated loops in GM12878, showing both shared and unique targets. **(h)** Canonical pathway enrichment analysis of genes associated with SE-linked loops mediated by each protein in GM12878, compared with SE regions themselves.

To further contextualize these differences, we integrated chromatin-state annotations for cell-type-specific loops. CTCF-specific anchors were most frequently localized within quiescent and repressed chromatin states, consistent with an insulative or structural role, whereas RNAPII-specific anchors were predominantly associated with active transcription start sites and enhancer states (Fig. 6c; Extended Data Fig. 12a). Quantitative enrichment analyses confirmed this dichotomy, with CTCF- and Cohesin-mediated loops showing strong enrichment at repressive chromatin features and RNAPII-mediated loops preferentially linked to transcriptionally active regions (Fig. 6d; Extended Data Fig. 12b).

We next connected loop architecture with transcriptional output by examining expression levels of genes associated with cell-type-specific loops. RNAPII- and Cohesin-mediated loops were linked to significantly higher gene expression compared with non-regulatory contacts, whereas CTCF-mediated loops showed only modest effects (Fig. 6e; Extended Data Fig. 11c). Among promoter-enhancer loops, loop strength positively correlated with differential gene expression across cell types, with stronger associations observed for RNAPII and Cohesin than for CTCF (Extended Data Fig. 13a). These results indicate that while CTCF-mediated loops primarily constrain chromatin topology, Cohesin and RNAPII-mediated loops more directly modulate transcriptional responsiveness in a context-dependent manner.

We further explored the relationship between protein-mediated loops and super-enhancers (SEs) ^39^. Cell-type-specific loops mediated by Cohesin and RNAPII were significantly enriched in the vicinity of cell-type-specific SEs, whereas CTCF-mediated loops showed weaker and less consistent enrichment across cell types (Fig. 6f; Extended Data Fig. 13b). This pattern suggests that architectural scaffolding provided by CTCF is selectively coupled to transcriptional amplification through Cohesin- and RNAPII-mediated interactions in regulatory contexts associated with high enhancer activity. At the gene level, RNAPII-mediated loops targeted the largest and most distinct set of genes across cell types, whereas CTCF and Cohesin shared a substantial fraction of their targets, consistent with their coordinated roles in organizing chromatin structure (Fig. 6g; Extended Data Fig. 14a). Functional enrichment analyses of SE-associated loop genes further revealed complementary regulatory programs among the three proteins (Fig. 6h). In GM12878, CTCF-associated SE loops were enriched for chromatin organization and architectural maintenance, Cohesin-associated loops for replication and cohesion-related pathways, and RNAPII-associated loops for immune activation and transcriptional execution. Similar trends were observed across additional cell types (Extended Data Fig. 14b).

Collectively, these analyses demonstrate that protein-mediated chromatin loops are not intrinsically defined by protein identity alone, but are functionally deployed in a context-dependent manner. CTCF, Cohesin and RNAPII contribute distinct yet coordinated looping programs that collectively balance structural constraint and regulatory flexibility, providing a mechanistic link between three-dimensional genome organization and transcriptional control across cellular contexts.

### Context-dependent reorganization of RNAPII-mediated chromatin architecture on ecDNA in cancer

Extrachromosomal DNA (ecDNA) represents an extreme regulatory context in cancer genomes, characterized by focal copy-number amplification and dense clustering of regulatory elements ^40–42^. To examine how protein-mediated chromatin architecture is reorganized under this context, we analyzed RNAPII-mediated chromatin interactions in three cancer cell types with available ecDNA contact maps generated by ChIA-Drop ^43^. Circular visualization of representative ecDNA regions revealed pronounced copy-number amplification accompanied by dense occupancy of RNAPII, MED1 and H3K27ac, indicative of highly active transcriptional hubs (Fig. 7a). Consistently, ChIP-seq signals of RNAPII, MED1 and H3K27ac were strongly enriched around ecDNA-associated contact anchors in PC3DM, with similar enrichment patterns observed across biological replicates in COLO320DM and GBM171DM (Fig. 7b; Supplementary Fig. 15a). Genome browser views further confirmed concordant spatial localization of ChIA-Drop interaction signals and transcription-associated epigenomic marks within ecDNA regions across all three cancer cell types (Supplementary Fig. 15b).

**Fig. 7:**
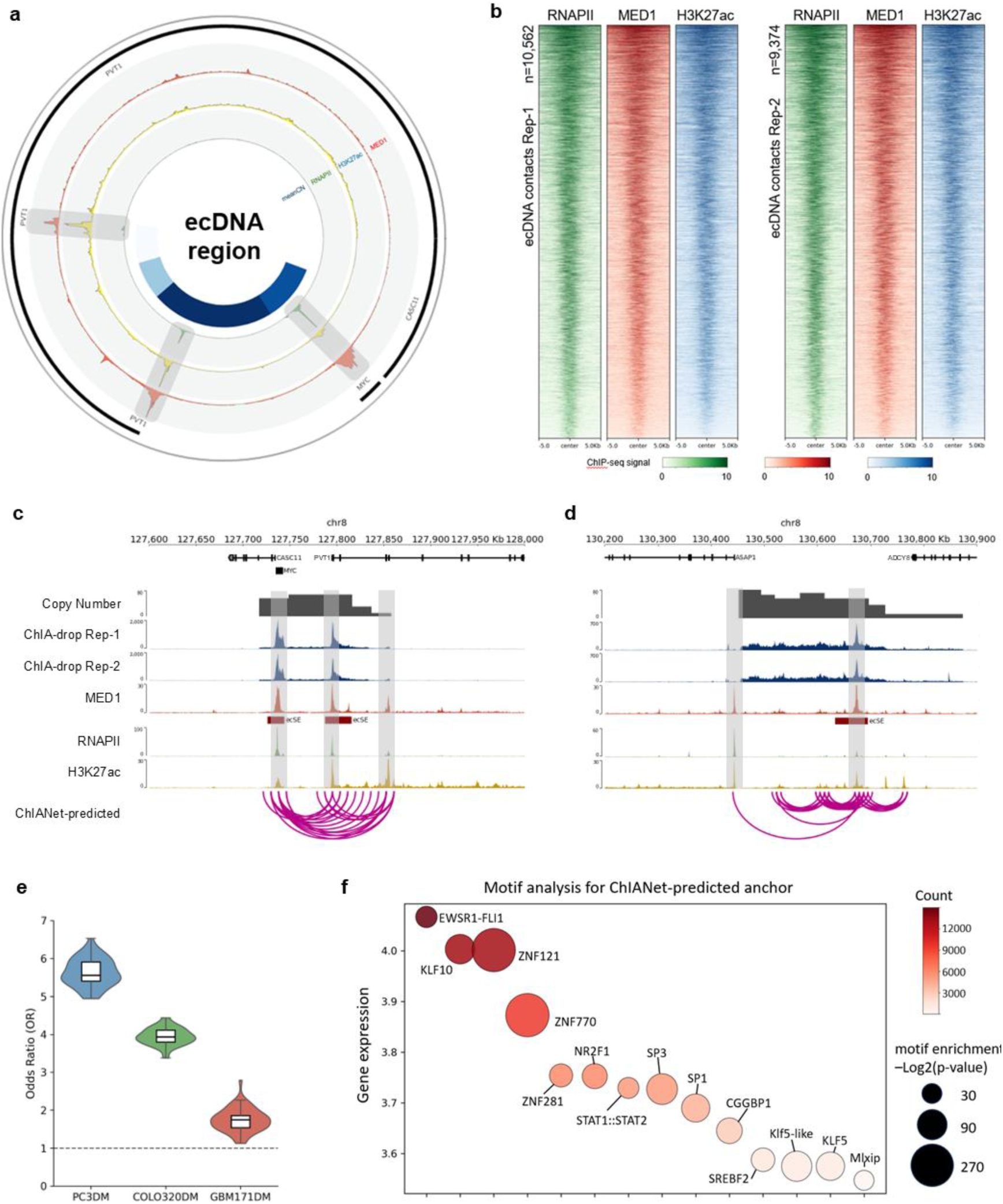
RNAPII-mediated chromatin looping at ecDNA regions in cancer cells. **(a)** Circular overview of a representative ecDNA region. From inner to outer tracks, the circos plot shows average copy-number profiles, followed by ChIP-seq signal intensities of RNAPII, H3K27ac and MED1. Gene annotations are shown in black. **(b)** Heatmaps showing ChIP-seq signal densities of RNAPII, MED1 and H3K27ac centered on ecDNA-associated contact anchors in PC3DM cells. Results from two biological replicates are shown (Rep-1, n = 10,562; Rep-2, n = 9,374). Signals are displayed within ± 5 kb of anchor centers. Color scales indicate normalized ChIP-seq signal intensities. **(c, d)** Genome browser views of two high-copy-number regions on chromosome 8. Tracks show copy-number profiles, ChIA-drop interaction signals from two biological replicates, MED1, RNAPII and H3K27ac ChIP-seq signals, and ChIANet-predicted RNAPII-mediated chromatin loops. EcDNA-associated regions are highlighted in grey. Genomic coordinates are indicated above each panel. **(e)** Enrichment analysis of ChIANet-predicted RNAPII loop anchors at ecDNA hubs compared with randomly sampled RNAPII peaks across PC3DM, COLO320DM and GBM171DM cells. Violin plots show odds ratios calculated using Fisher’s exact tests. The dashed line indicates an odds ratio of 1. **(f)** Motif analysis of ChIANet-predicted RNAPII loop anchors in PC3DM cells. Each point represents a transcription factor motif. Point size denotes motif enrichment significance (-log2 P value). Color intensity indicates RNA-seq-derived gene expression levels (raw expression counts) of the corresponding transcription factors. The y-axis shows gene expression levels.

Motivated by the close coupling between ecDNA contacts and transcription-associated epigenomic features, we applied ChIANet to predict RNAPII-mediated chromatin loops in the three cancer cell types. Predicted loops formed dense interaction networks within high-copy-number ecDNA regions, consistent with previously reported ecDNA-associated interaction structures detected (Fig. 7c, d). Quantitative enrichment analysis showed that ChIANet-predicted RNAPII loop anchors were significantly overrepresented at ecDNA contact regions compared with distance-matched random RNAPII peak sets across all three cancer cell types (Fig. 7e), indicating preferential deployment of RNAPII-mediated loops within ecDNA topology. Stratification of ChIA-Drop-defined n-way Genome Extrusion Modules (nGEMs) by interaction strength further revealed a positive association between nGEM connectivity and MED1 and H3K27ac signal intensities in PC3DM and COLO320DM, whereas weaker trends were observed in GBM171DM (Supplementary Fig. 12).

Despite sharing a common RNAPII-centered architectural framework, ecDNA-associated looping networks exhibited pronounced cancer-type-specific regulatory signatures. Motif analysis of ChIANet-predicted RNAPII loop anchors revealed distinct transcription factor landscapes coupled to expression across cancer cell types (Fig. 7f; Supplementary Fig. 15c). PC3DM showed enrichment of STAT1::STAT2, SP1/SP3 and KLF-family motifs, together with metabolic regulators such as SREBF2 and MLXIP. In COLO320DM, enriched motifs included TCF4 and additional GC-rich or architectural factors (for example MAZ, HMGA1 and SP1/SP3), consistent with a dominant Wnt/ β -catenin-associated regulatory program. In GBM171DM, enriched motifs included immediate-early and regulatory transcription factors such as EGR1 and ZEB1/ZEB2, together with SP1/SP3 and KLF-related motifs. Consistent with these motif-level distinctions, Gene Ontology enrichment analysis of genes associated with predicted RNAPII loop anchors identified both shared and cancer-type-specific functional programs (Extended Data Fig. 16). PC3DM was enriched for translation-related processes and chromatin organization, COLO320DM for chromatin and DNA organization and β-catenin-TCF complex assembly, whereas GBM171DM showed enrichment of post-transcriptional regulation, RNA catabolic processes and immune-related signaling.

Together, these results demonstrate that ecDNA constitutes an extreme functional context in which RNAPII-mediated chromatin architecture is extensively reorganized, forming dense and highly connected looping networks that couple copy-number amplification with transcriptional regulation in a cancer-type-specific manner.

## DISCUSSION

Understanding how specific chromatin-binding proteins shape the three-dimensional genome remains a central challenge in regulatory genomics ^44,45^. Existing Hi-C-based approaches cannot disentangle protein-specific contributions to genome folding, whereas sequence-only models lack the regulatory context required to capture condition-dependent chromatin organization. ^33^. To address these limitations, we developed ChIANet, a multimodal deep learning framework that integrates DNA sequence and protein-specific ChIP-seq data to enable *de novo* prediction of protein-mediated chromatin interactions. By jointly modeling contact maps and chromatin loops within a unified encoder-decoder architecture, ChIANet accurately reconstructs genome-wide chromatin organization mediated by CTCF, Cohesin and RNAPII, capturing both local and long-range architectural features and generalizing robustly across cell types without retraining.

Systematic application of ChIANet across seven human cell lines reveals a conserved hierarchical organization of the 3D genome in which distinct chromatin regulators play coordinated but nonredundant roles. Architectural proteins CTCF and Cohesin establish a stable structural backbone that preserves domain-scale organization across cellular contexts, whereas RNAPII mediates a more dynamic, transcription-associated interaction layer. Loop-level analyses further support this division of labor: CTCF- and Cohesin-mediated loops are longer, more conserved and highly connected, while RNAPII-mediated loops are shorter, more variable and preferentially associated with promoters and enhancers. Integration with chromatin-state annotations, gene expression and super-enhancer landscapes demonstrates that Cohesin and RNAPII loops are tightly coupled to transcriptional output, whereas CTCF primarily contributes to structural insulation. Together, these results support a multilayered model of genome organization in which architectural stability and regulatory flexibility are jointly encoded by different protein-mediated interaction networks.

Importantly, extending ChIANet to cancer genomes carrying extrachromosomal DNA (ecDNA) reveals that this hierarchical organization is not fixed, but can be profoundly reorganized under extreme functional contexts. EcDNA represents a regulatory environment characterized by circular topology, high copy number and dense clustering of regulatory elements, in which canonical chromosomal constraints on genome folding are relaxed. In this context, RNAPII-mediated interactions no longer constitute a secondary regulatory layer, but instead dominate chromatin architecture, forming dense, highly interconnected looping networks that couple transcriptional coactivator recruitment with amplified gene expression. Quantitative enrichment, nGEM stratification and epigenomic coupling analyses indicate that RNAPII assumes an architectural role on ecDNA that is qualitatively distinct from its function on linear chromosomes. Moreover, despite sharing a common RNAPII-centered framework, ecDNA-associated interaction networks display pronounced cancer-type-specific transcription factor and functional signatures, underscoring how functional context and regulatory demand jointly reshape protein-mediated 3D genome organization.

Notably, the conceptual design of ChIANet is not restricted to the specific chromatin regulators examined here. By conditioning chromatin interaction prediction on protein-specific binding profiles together with underlying genomic sequence, ChIANet provides a generalizable framework that is, in principle, extensible to other chromatin-associated factors, including transcription factors, cofactors and chromatin modifiers, provided that appropriate binding and genomic context information are available. This flexibility positions ChIANet as a broadly applicable platform for interrogating how diverse regulatory proteins contribute to three-dimensional genome organization across distinct cellular states and biological contexts.

Together, these findings advance a context-dependent view of chromatin architecture in which protein identity alone does not uniquely determine structural function. Instead, the architectural role of a given protein emerges from the interplay between its biochemical activity, regulatory targets and the genomic environment in which it operates. By capturing both conserved organizational principles and context-specific architectural rewiring, ChIANet provides a scalable framework for decoding how protein-mediated chromatin interactions adapt across cell types, regulatory states and disease-associated genome configurations. Beyond descriptive modeling, the protein-conditioned nature of ChIANet naturally suggests future opportunities for in silico interrogation of genome folding. In principle, systematic perturbation of protein binding landscapes, regulatory states or genome configurations could enable predictive exploration of how changes in regulatory inputs reshape three-dimensional genome organization. While such applications remain to be explored, these perspectives highlight the potential of ChIANet not only as a predictive model, but also as a hypothesis-generating framework for dissecting the functional logic that shapes the three-dimensional genome ^46,47^.

## METHODS

### ChIA-PET data processing

We used four ChIA-PET datasets in this study: CTCF (GM12878), CTCF (H1), Cohesin (GM12878) and RNAPII (GM12878) (Supplementary Table 1). All datasets are publicly available from ENCODE ^48^ (http://www.encodeproject.org/) and 4DN ^49^ (https://data.4dnucleome.org/). For Cohesin-mediated chromatin interactions, we used the ChIA-PET dataset generated with antibodies against RAD21 and SMC1A, and processed them jointly as cohesin ChIA-PET to capture cohesin-associated loops. Raw reads were processed using the ChIA-PIPE ^50^ pipeline (https://github.com/TheJacksonLaboratory/ChIA-PIPE) to generate contact maps and loop calls aligned to the GRCh38 reference genome. For contact maps, interactions were called using raw counts at 10-kb resolution, and the matrices were log2 transformed with a pseudocount of 1 to stabilize variance.

### Loop quality control and filtering

Loop calls generated by ChIA-PIPE were further filtered to ensure high-confidence interactions. We first removed inter-chromosomal loops to retain only intra-chromosomal interactions. Loops with a supporting PET count (Score) < 5 were discarded to exclude weakly supported contacts. We also excluded short-range loops (< 50 kb) to remove potential self-ligation or local background artifacts. The remaining high-confidence intra-chromosomal loops were used in all downstream analyses. The number of loops before and after filtering for each dataset is summarized in Supplementary Table 2, and an example of quality control before and after filtering is shown in Supplementary Fig. 2.

### ChIP-seq data processing

We collected 21 human ChIP-seq datasets from ENCODE ^48^ (http://www.encodeproject.org/) corresponding to three proteins (CTCF, RAD21, RNAPII) across seven cell types. For each protein-cell-type combination, we downloaded all available isogenic replicate BAM files aligned to hg38 and merged replicates to obtain a unified signal track. To ensure signal comparability across cell types, we used deepTools ^51^ (bamCoverage) to generate genome-wide coverage files in bigWig format with binSize = 1 bp and RPGC (reads per genomic content) normalization. The resulting tracks were further log2 transformed with a pseudocount of 1 and used as input feature signals for ChIANet.

### DNA sequence

The human reference genome (GRCh38/hg38) was downloaded from the UCSC Genome Browser ^52^. We used the primary assembly and kept all nucleotide symbols present in the FASTA file. In addition to the canonical bases (A, C, G and T), positions annotated as unknown (“N”) were preserved and treated as a separate category to avoid information loss in low-mappability or unresolved regions. For model input, the genome was converted to a five-channel one-hot representation corresponding to A, C, G, T and N, and the same reference sequence was used for all cell types and proteins to ensure a shared coordinate system.

### Training data construction

ChIANet was trained on GM12878 for three proteins—CTCF, Cohesin and RNAPII—using one model per protein. For each model, inputs consisted of the DNA sequence and the corresponding protein-specific ChIP-seq signal from a 2,097,152-bp (2.1-Mb) genomic window. The prediction targets were the protein-mediated contact map and loop matrix for the same window.ChIA-PET contact maps were initially generated at 10-kb resolution and then downsampled to 8,192-bp (256*256) resolution by bilinear interpolation to match the model output grid. Loop targets were represented as a 256*256 binary matrix initialized to zeros, with bins overlapping ChIA-PET loop anchors set to 1. For both map and loop outputs, only the upper-triangular entries were used as training labels to avoid redundancy.

To create training samples, we tiled the genome with a sliding window of 2.1 Mb and a stride of 50 kb. Windows overlapping centromeric or telomeric regions were excluded. Chromosomes were randomly assigned to training, validation and test sets at the chromosome level: chromosomes 3 and 15 were held out for validation, chromosomes 5 and 18 were held out for testing, and all remaining autosomes were used for training. This procedure yielded 4,672 training, 571 validation and 511 test samples.

### Model architecture

ChIANet was implemented in PyTorch ^53^ and consists of three main components: a 1D convolutional encoder, a Transformer module, and a multi-task convolutional decoder. DNA sequence and protein-specific ChIP-seq signals were concatenated along the channel dimension and jointly fed into the encoder.

The encoder begins with a 1D convolution layer (kernel size = 11, stride = 2) to capture local motif-like features. To reduce the input length from 2.1 Mb to 256 bins, the encoder applies 12 convolutional modules, each composed of a residual block followed by a scaling block. The residual block contains two 1D convolutions (kernel = 5, padding = 2), each followed by batch normalization and ReLU activation, with skip connections to facilitate gradient flow and long-range feature propagation. The scaling block performs downsampling via a stride-2 convolution (kernel = 5) followed by batch normalization and ReLU. Hidden dimensions were progressively increased (32, 32, 32, 32, 64, 64, 128, 128, 128, 128, 256, 256), resulting in a final encoder output of 256 bins * 256 channels.

The Transformer module contains eight self-attention layers with eight attention heads and 256-dimensional hidden units. Each layer includes multi-head attention, feed-forward layers, layer normalization and dropout (rate = 0.1). Sinusoidal positional encoding was applied to preserve genomic order. This module captures long-range dependencies between bins, enabling the model to learn spatial interaction patterns from sequence and ChIP-seq features ^36^.

The decoder consists of five dilated residual convolutional blocks, with dilation rates set to 2, 4, 8, 16 and 32 to ensure each output pixel has a receptive field covering the entire 2.1 Mb input. Each block includes two 2D convolutions (kernel = 3), batch normalization and ReLU activation with residual connections. Two final 1*1 convolutions produce the outputs: one for the contact map and one for the loop matrix. Both outputs are flattened to the upper-triangular part (length = 32,640). The contact-map branch is trained using mean squared error (MSE) loss, and the loop branch using binary cross-entropy (BCE) loss. A heteroscedastic uncertainty weighting strategy balances the two objectives, and gradients are propagated end-to-end for joint optimization ^37^.

### Model training and prediction

We trained three protein-specific ChIANet models—CTCF, Cohesin and RNAPII—using GM12878 data. Each model jointly optimized two tasks: contact map prediction and loop prediction. To balance the contributions of the two losses, we adopted a heteroscedastic uncertainty weighting strategy ^37^:

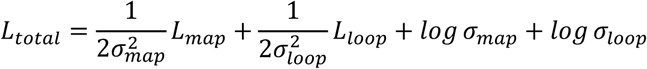

where *L*_*map*_ and *L*_*loop*_ denote the mean squared error (MSE) and binary cross-entropy (BCE) losses, respectively. The uncertainty parameters *σ*_*map*_ and *σ*_*loop*_ were jointly learned with model weights to adaptively reweight the two tasks during training.

Training was performed using the Adam ^54^ optimizer (initial learning rate 2e-4, batch size = 16) for 100 epochs. We employed a linear warm-up from 1e-5 followed by a cosine annealing schedule with a minimum learning rate of 1e-6. An early stopping strategy was used to prevent overfitting: training was terminated if the total loss did not decrease for 10 consecutive epochs. To enhance generalization, two forms of data augmentation were applied: (1) random shifts of the 2.1-Mb genomic window within ±0.36 Mb, and (2) reverse-complement flipping of the DNA sequence and corresponding contact/loop matrices with a probability of 0.5. Each model was trained on a single NVIDIA A6000 GPU (∼11 hours per model).

For in-cell predictions (GM12878), model inputs included the 2.1-Mb reference sequence and the protein-specific ChIP-seq profile. For cross-cell *de novo* predictions, we replaced the GM12878 ChIP-seq track with the corresponding track from the target cell type while keeping the reference genome sequence unchanged.

### Chromosome-scale prediction assembly

To generate chromosome-scale predictions, we tiled the genome using a sliding window of 2.1 Mb with a stride of 262,144 bp (one-eighth of the window size). For each window, ChIANet produced predictions for both the contact map and loop matrix. Adjacent regional outputs were then merged to reconstruct the full chromosome-scale maps. To correct for regions of overlap between neighboring windows, we calculated the number of overlapping predictions per pixel and normalized each pixel value by its corresponding overlap count. The resulting genome-wide contact maps and loop matrices provide continuous, bias-corrected representations of protein-mediated chromatin interactions suitable for downstream quantitative and visualization analyses (Supplementary Fig. 7, 8).

### Distance-stratified correlation and AUPR analysis

Distance-stratified metrics were computed to evaluate model performance across genomic distances. For each predicted 2.1-Mb window, we measured the average performance as a function of distance *i* (in bins), corresponding to diagonals offset by *i* from the main diagonal of the contact matrix. The distance-stratified correlation was defined as the Pearson correlation coefficient between the predicted and experimental values along the *i*-th offset diagonal. Similarly, the distance-stratified AUPR was calculated by comparing the predicted and experimental loop matrices along the same offset positions. For both metrics, values were averaged across all genomic windows to obtain genome-wide estimates of model performance as a function of genomic distance.

### Comparison with existing methods

Due to no existing framework specifically addresses genome-wide prediction of protein-mediated chromatin interactions, we benchmarked ChIANet against previously published models that represent the most relevant variants for each prediction task, using task-specific evaluation metrics to ensure fair comparison.

For the contact map prediction task, we compared ChIANet with three state-of-the-art Hi-C prediction models— Akita ^24^, Orca ^25^, and C.Origami ^26^ — which are most closely aligned in architecture and objective. Other methods such as DeepC ^27^ and Epiphany ^28^, which rely on multi-omics pretraining, and ChromaFold ^55^, which uses single-cell ATAC-seq as input, were excluded as they are not directly comparable to our protein-specific setting.

To adapt Akita for this study, we modified the model’s input and output window size from 1 Mb to 2.1 Mb, and adjusted the output resolution to 256*256 to match ChIANet’s prediction scale. For Orca, we selected its 2.1-Mb (8,192-bp resolution) configuration. For C.Origami, we removed ATAC-seq features and replaced CTCF ChIP-seq tracks with protein-specific ChIP-seq profiles corresponding to CTCF, Cohesin, or RNAPII. All models were retrained from scratch using the same GM12878 ChIA-PET datasets for the three proteins to ensure a consistent training and evaluation setup.

For the loop prediction task, we focused on models capable of *de novo*, genome-wide loop prediction rather than anchor-pair classification, as most existing approaches require pre-defined loop candidates. We therefore compared ChIANet with Peakachu ^29^ and DeepChIA-PET ^30^, two frameworks capable of predicting loops directly from chromatin contact data.

For Peakachu, we used 10-kb-resolution Hi-C contact maps as input and the quality-controlled, protein-mediated loops as training labels. For DeepChIA-PET, we adapted the original input and output dimensions from 250 bins (10 kb) to 210 bins to match ChIANet’s 2.1-Mb window. The model was trained using paired Hi-C and protein-specific ChIP-seq signals as input, with the same filtered ChIA-PET loops as supervision.

All models were trained and evaluated under identical genomic partitions and preprocessing protocols described above to enable fair, protein-specific performance comparison.

### Aggregate Peak Analysis (APA) of Predicted Loops

To evaluate the aggregate enrichment of predicted loops, we performed an Aggregate Peak Analysis (APA) using loops generated by ChIANet. First, all loop predictions from individual 2.1-Mb windows were merged to obtain chromosome-scale prediction matrices. From these genome-wide predictions, the top-scoring loops were selected at different TopN (N = 1k, 5k, 10k, 20k, 50k). Each set of predicted loop anchors was then mapped onto the corresponding experimental Hi-C contact matrices of the same cell type.

APA was conducted using the Juicer toolkit, which computes two standard quantitative metrics:

**P2LL (Peak-to-Lower-Left ratio):** the ratio of the central pixel intensity (representing the predicted loop) to the average intensity of the lower-left background region, reflecting local enrichment strength.

**ZscoreLL:** a standardized enrichment score calculated by normalizing the central signal to the variance of the lower-left background, capturing the statistical significance of enrichment.

APA heatmaps were visualized as 2D intensity plots centered on loop anchors, with darker central pixels indicating stronger enrichment of Hi-C contacts at predicted loop positions. Separate APA maps were generated for each Top N threshold to assess loop quality and the trade-off between precision and coverage.

### UMAP visualization of contact map manifolds

For each cell type, we embedded the genome-wide collection of predicted contact maps into a two-dimensional space for visualization. Specifically, for every 2.1-Mb window, the upper triangle of the ChIANet-predicted contact matrix (diagonal excluded) was vectorized to form a single feature vector per window. The resulting matrix of window-by-features was first reduced to 50 principal components using PCA ^56^ implemented in scikit-learn (Python), and then further reduced to two dimensions using UMAP ^57^ implemented in the umap-learn Python package. PCA outputs were passed directly to UMAP, and a fixed random seed was set for reproducibility. The two-dimensional UMAP embeddings were used to visualize the distribution and separation of protein-specific contact map windows within each cell type.

### Contact-map conservation score

For each genomic window *w* and protein *p*, we computed a cross-cell conservation score by averaging the pairwise Pearson correlations of the predicted contact maps across the *K* = 7 cell types:

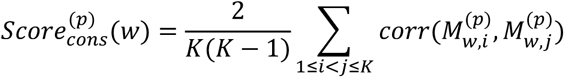

Here *corr*(·,·) is the same window-level PCC used elsewhere (computed on the upper triangle with the standard distance range), and 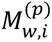 denotes the contact map for window *w* in cell type *i*. This yields a single scalar summarizing how similarly a window is organized across cell types.

### Pan-cell-type loop definition

To characterize protein-mediated chromatin loops across multiple cell types, we performed genome-wide ChIANet predictions for seven representative human cell lines (GM12878, H1, A549, HeLa-S3, IMR90, HepG2, and K562). For each protein (CTCF, Cohesin, and RNAPII), the top 10,000 predicted loops per cell type were combined to construct a pan-cell union set. Duplicate loops were merged based on genomic coordinates, and the corresponding prediction probability scores were retained for downstream analyses.

This process yielded 34,751 CTCF, 48,847 Cohesin, and 46,093 RNAPII loops, representing the comprehensive cross-cell landscape of protein-mediated chromatin interactions.

### Static score and classification of stable and variable loops

To quantify the cross-cell stability of each loop, we defined a static score based on the distribution of predicted loop probabilities across the seven cell types. For each loop, the prediction scores were first normalized to form a probability distribution *f*_*j*_ over cell types. The divergence between this empirical distribution and a uniform reference distribution *q*_*j*_ was measured using Kullback-Leibler (KL) divergence:

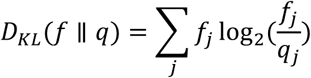

The static score was then defined as the inverse of this divergence, weighted by the mean predicted probability:

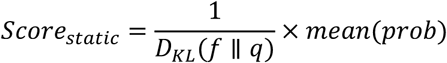

A higher static score indicates that a loop is consistently detected across multiple cell types (i.e., more conserved). Within each protein’s pan-cell union set, the top 20% of loops by static score were classified as stable loops, while the bottom 20% were designated as variable loops.

### Definition of housekeeping and cell-type-specific genes and loop enrichment analysis

RNA-seq data for the seven human cell types (GM12878, H1, A549, HeLa-S3, IMR90, HepG2, and K562) were obtained from the ENCODE (http://www.encodeproject.org/) consortium (Supplementary Table 3). For each gene, expression levels across the seven cell types were used to compute the same KL divergence-based variability metric as defined in Static score and classification of stable and variable loops. Genes with low expression (minimum TPM ≤ 1) were excluded. Genes within the top 10% of KL divergence (highly variable expression) were classified as cell-type-specific genes, whereas those within the bottom 10% (low variability) were designated as housekeeping genes.

To assess whether variable or stable loops were preferentially associated with specific gene classes, we examined the overlap between loop anchors and gene bodies or promoter regions. For each loop set (variable vs. stable), we tabulated the number of loops overlapping cell-type-specific or housekeeping genes and compared these to non-overlapping loops. Two-sided Fisher’s exact tests were used to evaluate the enrichment significance of each association.

### Definition of genomic regulatory elements

To establish a unified reference set of regulatory annotations, we compiled four major classes of genomic elements from ENCODE and GENCODE ^58^ resources (Supplementary Table 3).

#### Contact domain boundary

Contact domain boundaries were aggregated from high-resolution Hi-C data of GM12878, A549, HepG2, IMR90, and K562 (H1 and HeLa-S3 were unavailable). Boundaries from individual cell types were merged, and overlapping segments were unified to define a pan-cellular set of domain boundaries.

#### Enhancer

Enhancers were derived from H3K27ac narrowPeak files across seven ENCODE cell types (GM12878, H1, A549, HeLa-S3, IMR90, HepG2, and K562). Peaks from all cell types were combined, and overlapping regions were merged to generate a pan-cellular enhancer catalogue.

#### Silencer

Silencers were similarly defined from H3K27me3 narrowPeak files across the six cell types (A549 were unavailable), followed by merging of overlapping regions to construct a comprehensive silencer reference.

#### Promoter

Promoters were defined as ±500 bp windows flanking the transcription start site (TSS) of each annotated gene in GENCODE v29.

### Chromatin state enrichment analysis of cell-type-specific loops

Chromatin state annotations for seven cell types were obtained from the Roadmap Epigenomics Mapping Consortium ^59^ (15-state model; Supplementary Table 4). To simplify downstream analysis, the 15 chromatin states were consolidated into eight major categories: transcription start site (TSS: 1_TssA, 2_TssAFlnk), bivalent (BIV: 10_TssBiv, 11_BivFlnk), transcription (TX: 3_TxFlnk, 4_Tx, 5_TxWk), repressive (REPRESS: 13_ReprPC, 14_ReprPCWk), repeat-associated (REPEAT: 8_ZNF/Rpts), enhancer (ENH: 12_EnhBiv, 6_EnhG, 7_Enh), heterochromatic (HET: 9_Het), and quiescent (QUIES: 15_Quies).

For each cell type, cell-type-specific loops were defined by computing loop-specific z-scores based on the proportion of loops present across other cell types. Loop probabilities were first logit-transformed to approximate a normal distribution:

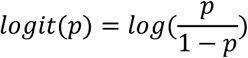

A pairwise comparison strategy was used to calculate a specificity z-score for each loop:

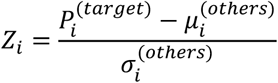

where 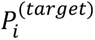 is the loop probability in the target cell type, and 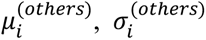 represent the mean and standard deviation across the remaining cell types. Loops ranked within the top 5% of *Z*_*i*_ values were designated as cell-type-specific loops for that cell type.

To assess the enrichment of chromatin states in cell-type-specific loop anchors, we constructed 2×2 contingency tables for each chromatin category. (1) anchors overlapping the chromatin state in specific loops, (2) anchors not overlapping the state in specific loops, (3) anchors overlapping the state in non-specific loops, and (4) anchors not overlapping the state in non-specific loops. Statistical significance of enrichment or depletion was assessed using two-sided Fisher’s exact tests.

### Identification and functional analysis of protein-mediated super-enhancer loops

Super-enhancers (SEs) are large clusters of enhancers densely occupied by transcriptional co-activators, and they play central roles in establishing cell identity-specific gene expression programs ^60^. To investigate the relationship between SEs and protein-mediated chromatin loops, we obtained SE annotations for seven human cell types from the SEdb database (v2.0) ^61^.

Cell-type-specific SEs were defined as those unique to a single cell type (i.e., not overlapping SE regions in any other cell type), and their distributions are shown in Supplementary Fig. 11a, c. Shared SEs were defined as SEs present in at least three cell types, as determined from the SE-sharing curve (Supplementary Fig. 11b).

Protein-mediated SE loops were identified by intersecting loop anchors with cell-type-specific SEs. For functional characterization, GREAT (Genomic Regions Enrichment of Annotations Tool, v4.0.4) ^62^ was applied to both SE regions and protein-mediated SE loops using default association parameters. Gene Ontology (GO) enrichment analyses were performed under the “Basal plus extension” association rule, and enriched biological processes were reported after multiple testing correction.

### EcDNA-associated chromatin contact definition

EcDNA-associated chromatin contacts were obtained from processed RNAPII ChIA-Drop interaction datasets for PC3DM, COLO320DM and GBM171DM cancer cells. Chromatin interactions were represented as multiplex interaction complexes with genomic interaction anchors. We directly used the provided interaction anchors for downstream analyses. Chromatin contacts were defined as ecDNA-associated if at least one interaction anchor overlapped annotated ecDNA regions. Contacts were classified as cis interactions (among ecDNA regions) or trans interactions (between ecDNA and chromosomal regions). For ecDNA-chromosomal interactions, only trans contacts with one anchor overlapping an ecDNA region and the other anchor mapping to chromosomal loci were retained. Overlapping anchors across libraries were merged to define interaction nodes, which were used for all subsequent analyses, including epigenomic signal aggregation, enrichment analysis and comparison with ChIANet-predicted RNAPII loop anchors.

### nGEM stratification and epigenomic association analysis

Higher-order chromatin interaction units were analyzed using n-way Genome Extrusion Modules (nGEMs) defined in the processed RNAPII ChIA-Drop datasets. Each nGEM represents a multiplex chromatin interaction complex comprising multiple genomic fragments identified within a single interaction unit. nGEMs were stratified by interaction complexity based on the number of participating fragments and grouped into deciles from low to high complexity. For each nGEM decile, epigenomic signal intensities of MED1 and H3K27ac were aggregated at the corresponding interaction anchors. Associations between nGEM complexity and epigenomic signals were evaluated using Spearman correlation across deciles and biological replicates.

### Identification of ecDNA-associated super-enhancers

EcDNA-associated super-enhancers (ecSEs) were identified by integrating MED1 ChIP-seq-defined super-enhancer annotations with annotated ecDNA regions. Super-enhancer intervals were obtained from MED1 ChIP-seq data using the ROSE algorithm with default parameters. Super-enhancers whose genomic coordinates overlapped annotated ecDNA regions were classified as ecDNA-associated super-enhancers (ecSEs). Genomic overlap was determined by interval intersection, and all super-enhancers satisfying this criterion were retained as ecSEs for downstream analyses.

### Enrichment of predicted loop anchors at ecDNA contacts

To assess whether ChIANet-predicted RNAPII loop anchors were preferentially enriched at ecDNA-associated chromatin contacts, we compared their overlap with ecDNA contact anchors against a distance-matched random background. Genome-wide RNAPII ChIP-seq peaks were used as the background pool. For each set of predicted loop anchors, random RNAPII peaks were sampled to match the genomic distance distribution to ecDNA contact anchors, thereby controlling for distance-dependent biases.

Enrichment was evaluated using Fisher’s exact test by contrasting the number of predicted loop anchors overlapping ecDNA contact anchors with the corresponding overlap observed for the distance-matched random RNAPII peaks. This procedure was repeated 50 times with independent random samplings. Odds ratios (ORs) were calculated for each iteration, and the distribution of ORs was used to summarize enrichment across randomizations.

### Motif enrichment analysis of ecDNA-associated RNAPII loop anchors

Motif enrichment analysis was performed on ChIANet-predicted RNAPII loop anchors that overlapped annotated ecDNA regions. Genomic sequences corresponding to these anchors were analyzed using the Analysis of Motif Enrichment (AME) tool from the MEME suite, with all parameters set to default values. Motif enrichment was assessed against the JASPAR CORE 2026 transcription factor binding motif database. To integrate motif enrichment with transcriptional activity, RNA-seq expression data were used to annotate transcription factors corresponding to enriched motifs. Raw RNA-seq read counts were used as the measure of gene expression, and transcription factors with expression counts greater than 300 were retained for downstream analysis. Motifs exhibiting statistically significant enrichment were identified based on a threshold of -log₂(p-value) > 20. Analyses focused on transcription factor binding motifs meeting both the enrichment and expression criteria.

### Gene Ontology enrichment analysis of ChIANet-predicted loop anchors

Gene Ontology (GO) enrichment analysis was performed to characterize functional programs associated with ChIANet-predicted chromatin loop anchors in PC3DM, COLO320DM and GBM171DM cells. Predicted loop anchors for each cell type were analyzed separately. Loop-associated genes were assigned using the Genomic Regions Enrichment of Annotations Tool (GREAT, v4.0.4) by associating loop anchor regions with nearby genes. GREAT analyses were conducted under the basal plus extension association rule using default parameters. Enrichment was assessed for GO biological process terms, and statistically significant categories were identified after multiple testing correction. All GO enrichment analyses were performed independently for each cancer cell type using the corresponding sets of predicted loop anchors.

## DATA AVAILABILITY

Most of the ChIA-PET, ChIP-seq, RNA-seq and Hi-C datasets used in this study were obtained from public resources, including the ENCODE portal and the 4D Nucleome (4DN) data portal, with accession codes provided in the corresponding Methods sections or Supplementary Information. ecDNA-associated chromatin interaction datasets for the three cancer cell types (PC3DM, COLO320DM and GBM171DM) were obtained from the NCBI Gene Expression Omnibus under accession number GSE275060.

## CODE AVAILABILITY

The code for ChIANet is available at https://github.com/lhy0322/ChIANet.

## SUPPLEMENTARY DATA

Supplementary Tables 1-4 and Figs. 1-12.

## ACKNOWLEDGEMENTS

The authors thank colleagues for helpful discussions. Detailed acknowledgements and funding information will be provided upon journal submission.

## EXTENDED DATA

**Extended Data Fig. 1:**
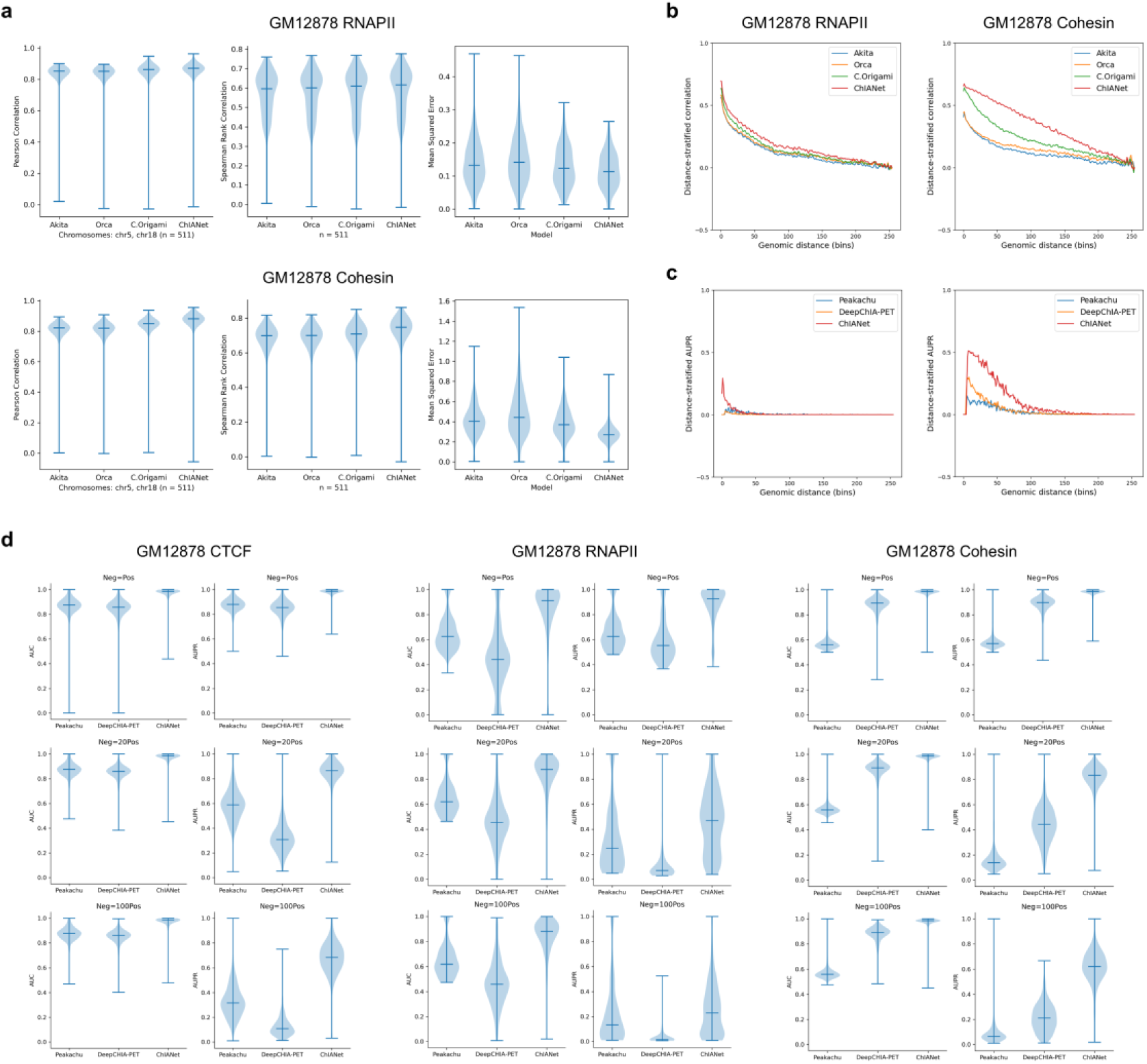
ChIANet performance on RNAPII- and Cohesin-mediated chromatin interactions. **(a)** Benchmarking ChIANet against Akita, Orca and C.Origami on GM12878 RNAPII (top) and GM12878 Cohesin (bottom). For each protein, Pearson correlation (left) and Spearman rank correlation (middle) were computed between predicted and experimental contact maps across all 2.1-Mb windows (n = 511). Right, window-level mean squared error (MSE) between predicted and experimental contact maps for the same models, showing that ChIANet achieves comparable or lower reconstruction error. **(b)** Distance-stratified Pearson correlation between predicted and experimental contact maps for GM12878 RNAPII (left) and GM12878 Cohesin (right), showing that ChIANet maintains competitive or improved correlation at longer genomic distances. **(c)** Distance-stratified AUPR for loop prediction on GM12878 RNAPII (left) and GM12878 Cohesin (right), compared with Peakachu and DeepChIA-PET. **(d)** Genome-wide loop prediction under different positive-to-negative ratios (1:1, 1:20 and 1:100) for GM12878 CTCF, GM12878 RNAPII and GM12878 cohesin. Violin plots show AUC (left in each panel) and AUPR (right in each panel) for Peakachu, DeepChIA-PET and ChIANet, indicating that ChIANet remains robust under increasingly imbalanced classification settings.

**Extended Data Fig. 2:**
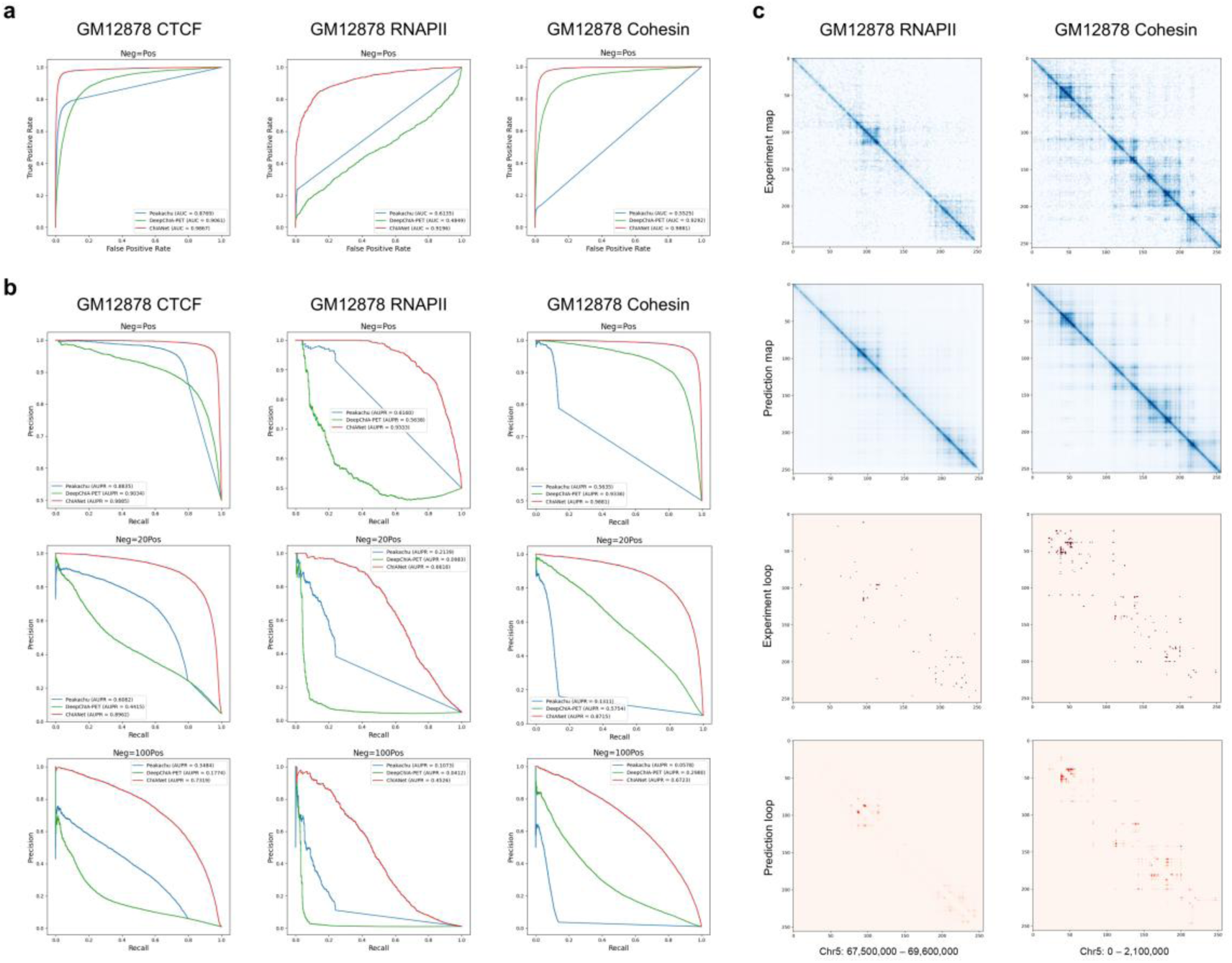
ChIANet performance on RNAPII- and Cohesin-mediated chromatin interactions. **(a)** Receiver operating characteristic (ROC) curves comparing loop prediction performance of Peakachu, DeepChIA-PET and ChIANet for GM12878 CTCF (left), RNAPII (middle) and Cohesin (right) at a 1:1 positive-to-negative ratio. ChIANet achieves the highest AUC across all proteins. **(b)** Precision-recall (PR) curves under increasingly imbalanced conditions (positive-to-negative ratios of 1:1, 1:20 and 1:100) for the same proteins. ChIANet consistently maintains superior AUPR compared with baseline models, demonstrating robustness to class imbalance. **(c)** Representative examples of experimental and predicted contact maps (top) and loop matrices (bottom) for RNAPII (left) and Cohesin (right) in GM12878. Experimental contact maps and loop calls are shown in the upper panels, while ChIANet predictions are shown below, highlighting accurate reconstruction of protein-mediated chromatin interactions at the 2.1-Mb scale.

**Extended Data Fig. 3:**
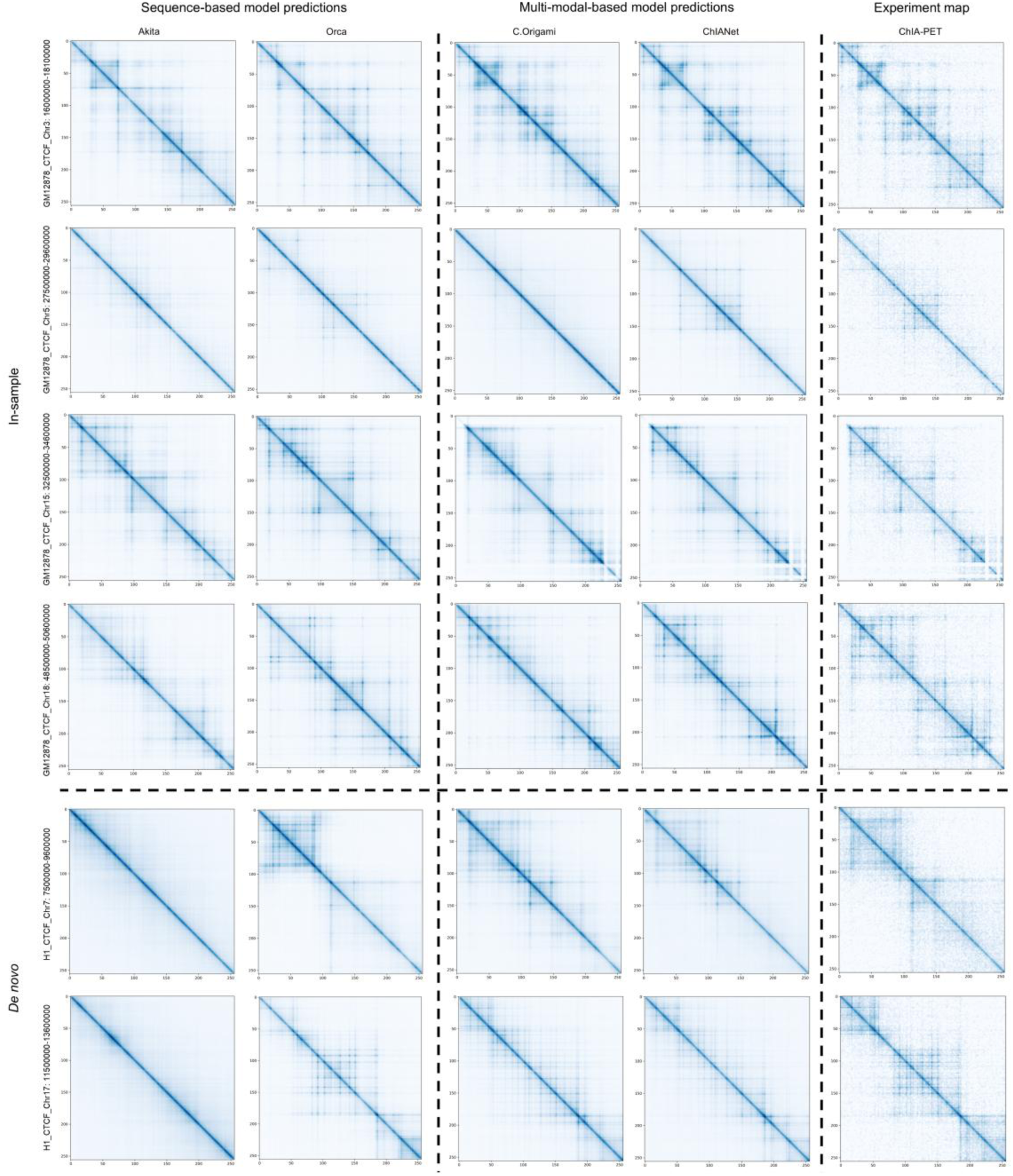
Representative examples of protein-specific contact map predictions across models. Comparison of predicted and experimental contact maps for randomly selected 2.1-Mb genomic windows across CTCF-mediated chromatin interactions in GM12878 and H1 cell types. Models are grouped by input modality: sequence-based models (Akita, Orca), multi-modal models (C.Origami, ChIANet), and the corresponding experimental ChIA-PET contact maps. For each protein and region, predictions from Akita, Orca, C.Origami, and ChIANet are shown alongside experimental ChIA-PET maps. ChIANet demonstrates stronger agreement with experimental contact patterns, accurately recovering both short- and long-range structures, while maintaining cell-type specificity in *de novo* predictions (bottom panels).

**Extended Data Fig. 4:**
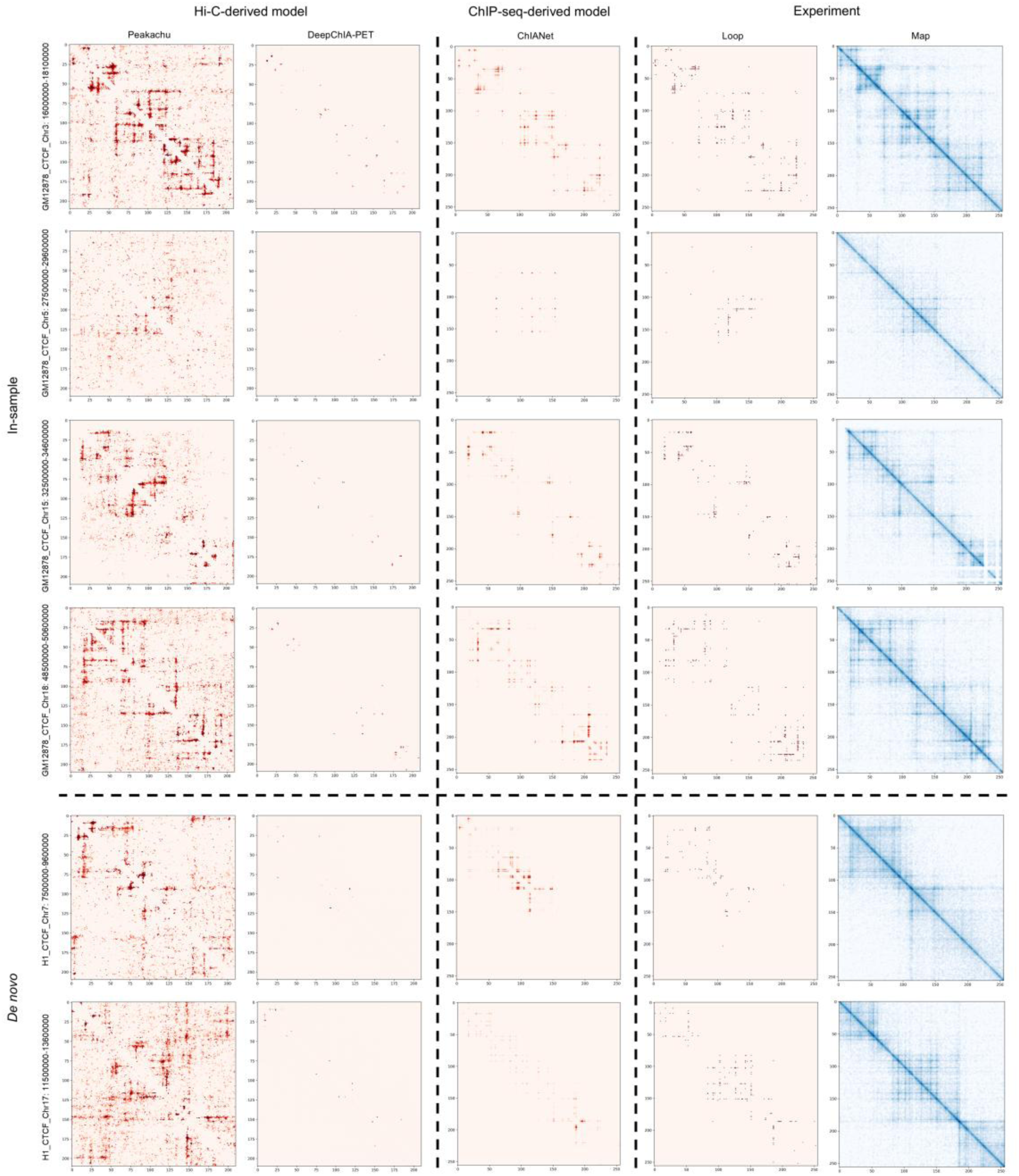
Representative examples of protein-specific loop predictions across models. Comparison of loop prediction outputs from Peakachu, DeepChIA-PET, and ChIANet for randomly selected 2.1-Mb genomic regions across CTCF-mediated chromatin interactions in GM12878 and H1 cell types. Models are grouped by input modality: Hi-C-derived models (Peakachu, DeepChIA-PET), ChIP-seq-derived model (ChIANet), and the corresponding experimental ChIA-PET loops and contact maps. For each region, predicted loop matrices (red) are shown alongside experimental contact maps (blue). ChIANet predictions exhibit higher spatial precision and cleaner topological boundaries compared to Hi-C-based models, capturing both prominent and distal loops visible in experimental data. *De novo* predictions in H1 demonstrate the model’s generalization ability across cell types using only sequence and protein-specific ChIP-seq as input.

**Extended Data Fig. 5:**
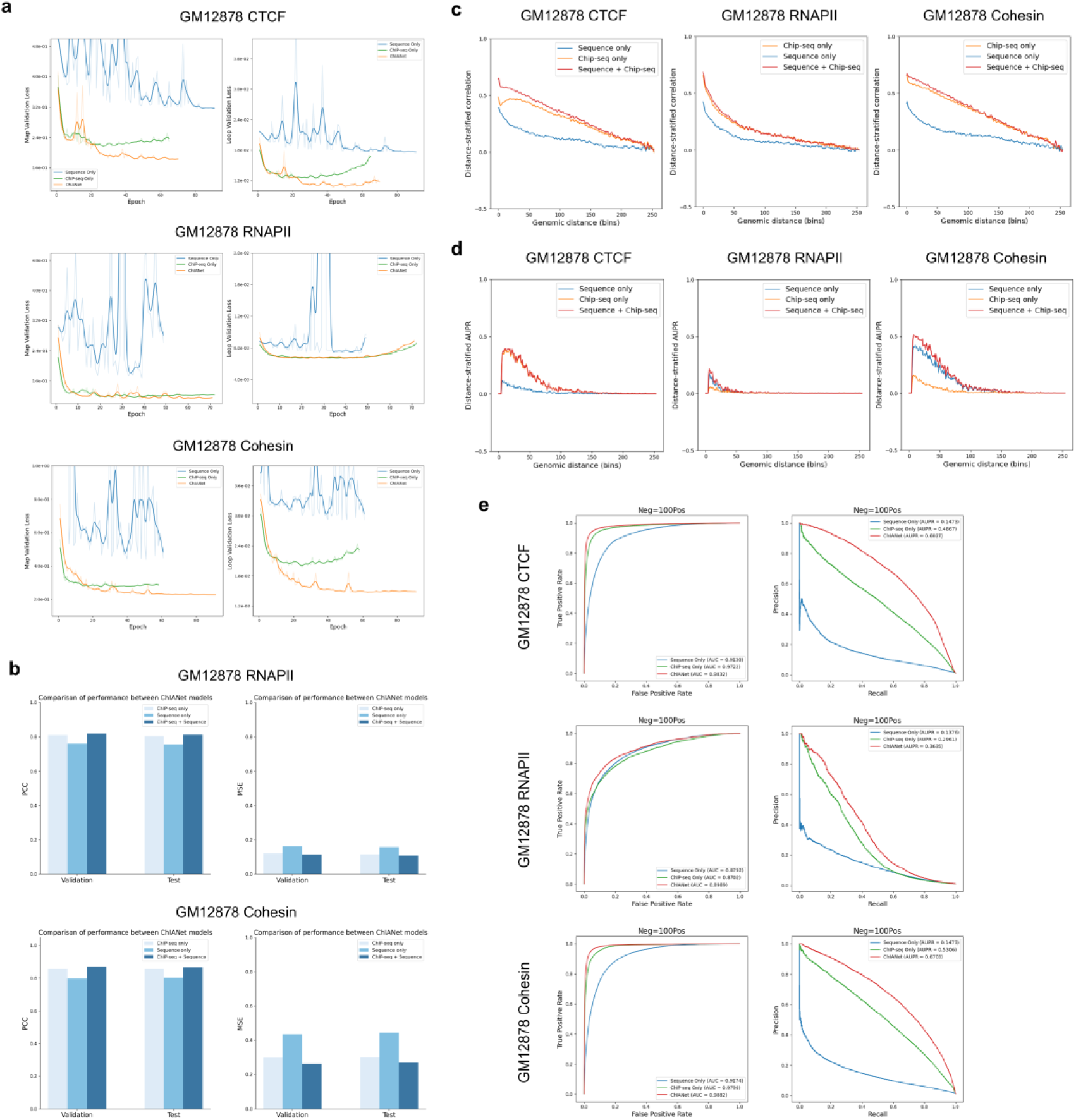
Ablation analysis of ChIANet across proteins and evaluation of multimodal contributions. **(a)** Training dynamics of the sequence only, ChIP-seq only, and sequence + ChIP-seq (full) ChIANet models on GM12878 datasets for CTCF, RNAPII, and Cohesin. Curves show the contact map loss (left) and loop loss (right) across training epochs, indicating improved convergence and lower task-specific losses when integrating both input modalities. **(b)** Comparison of contact map prediction performance across models for RNAPII and Cohesin on the validation and test sets. Bars represent Pearson correlation coefficients (PCC, left) and mean squared error (MSE, right), showing consistent performance gains when combining ChIP-seq and sequence features. **(c)** Distance-stratified Pearson correlations between predicted and experimental contact maps for CTCF, RNAPII and Cohesin, demonstrating that multimodal integration enhances long-range contact modeling compared to single-modality inputs. **(d)** Distance-stratified area under the precision-recall curve (AUPR) for loop prediction, showing that the combined model achieves higher precision-recall performance across genomic distances. **(e)** Chromosome-scale comparisons of AUC and AUPR for loop prediction under a 1:100 negative-to-positive sampling ratio for CTCF, RNAPII, and Cohesin. The full model consistently achieves the highest overall accuracy, highlighting that both DNA sequence and ChIP-seq signals are essential for accurately modeling protein-mediated chromatin interactions.

**Extended Data Fig. 6:**
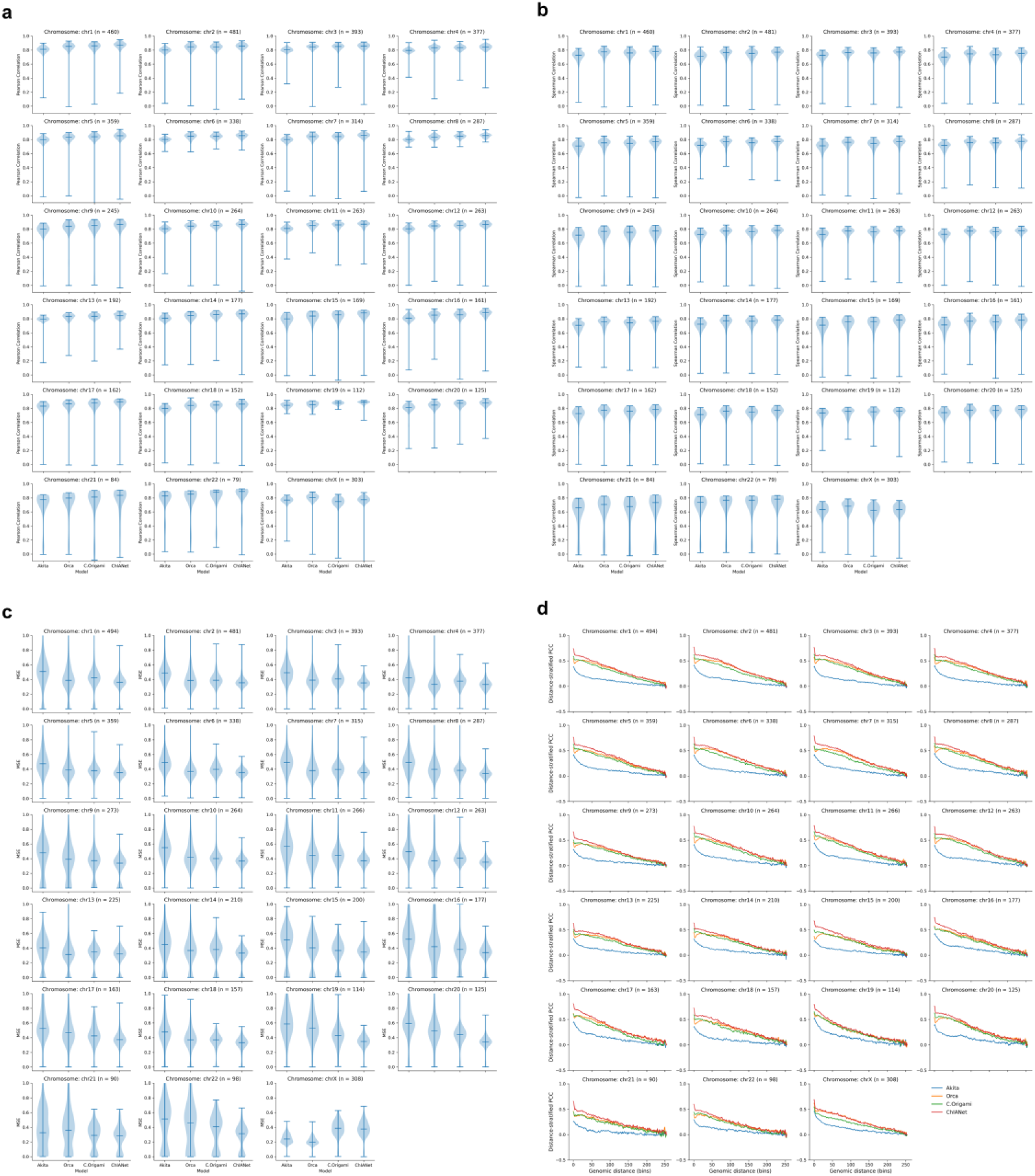
Genome-wide chromosome-level evaluation of *de novo* contact map prediction in H1 CTCF. **a-c,** Per-chromosome violin plots showing the distribution of ChIANet and baseline model performance (Akita, Orca, and C.Origami) across all 2.1-Mb windows in the *de novo* prediction of H1 CTCF-mediated contact maps. Each violin represents the variability of window-level scores within each chromosome, with the number of evaluated windows (n) indicated above each panel. ChIANet consistently achieves higher correlation and lower error across most chromosomes. **(a)** Pearson correlation (PCC). **(b)** Spearman rank correlation. **(c)** Mean squared error (MSE). **(d)** Distance-stratified Pearson correlation per chromosome, showing the decay of predictive accuracy with increasing genomic distance. ChIANet maintains stronger long-range correlation than sequence-based (Akita, Orca) and multimodal (C.Origami) models, demonstrating robust genome-wide modeling of distal CTCF-mediated chromatin interactions.

**Extended Data Fig. 7:**
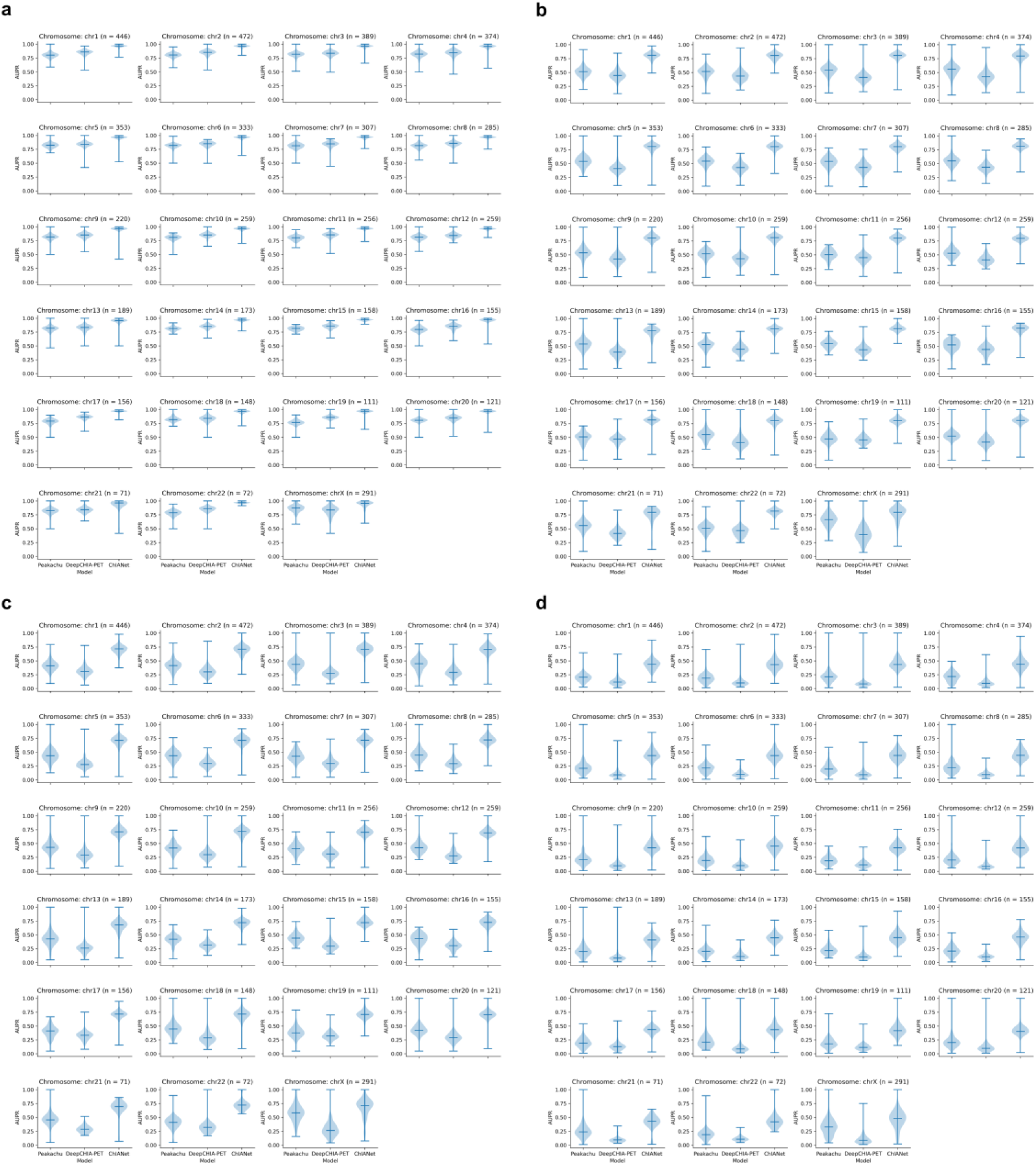
Genome-wide chromosome-level evaluation of *de novo* loop prediction in H1 CTCF. **a-d**, Per-chromosome violin plots showing window-level AUPR for ChIANet and baseline loop detectors (Peakachu, DeepChIA-PET) across all 2.1-Mb windows in the *de novo* prediction setting. Each violin summarizes the distribution within one chromosome; the number of evaluated windows (n) is indicated above each panel. Class-imbalance is varied by the positive:negative ratio used to compute AUPR: (a) 1:1, (b) 1:10, (c) 1:20, (d) 1:100. This figure provides genome-wide, chromosome-resolved comparisons of AUPR under increasing imbalance for the three models.

**Extended Data Fig. 8:**
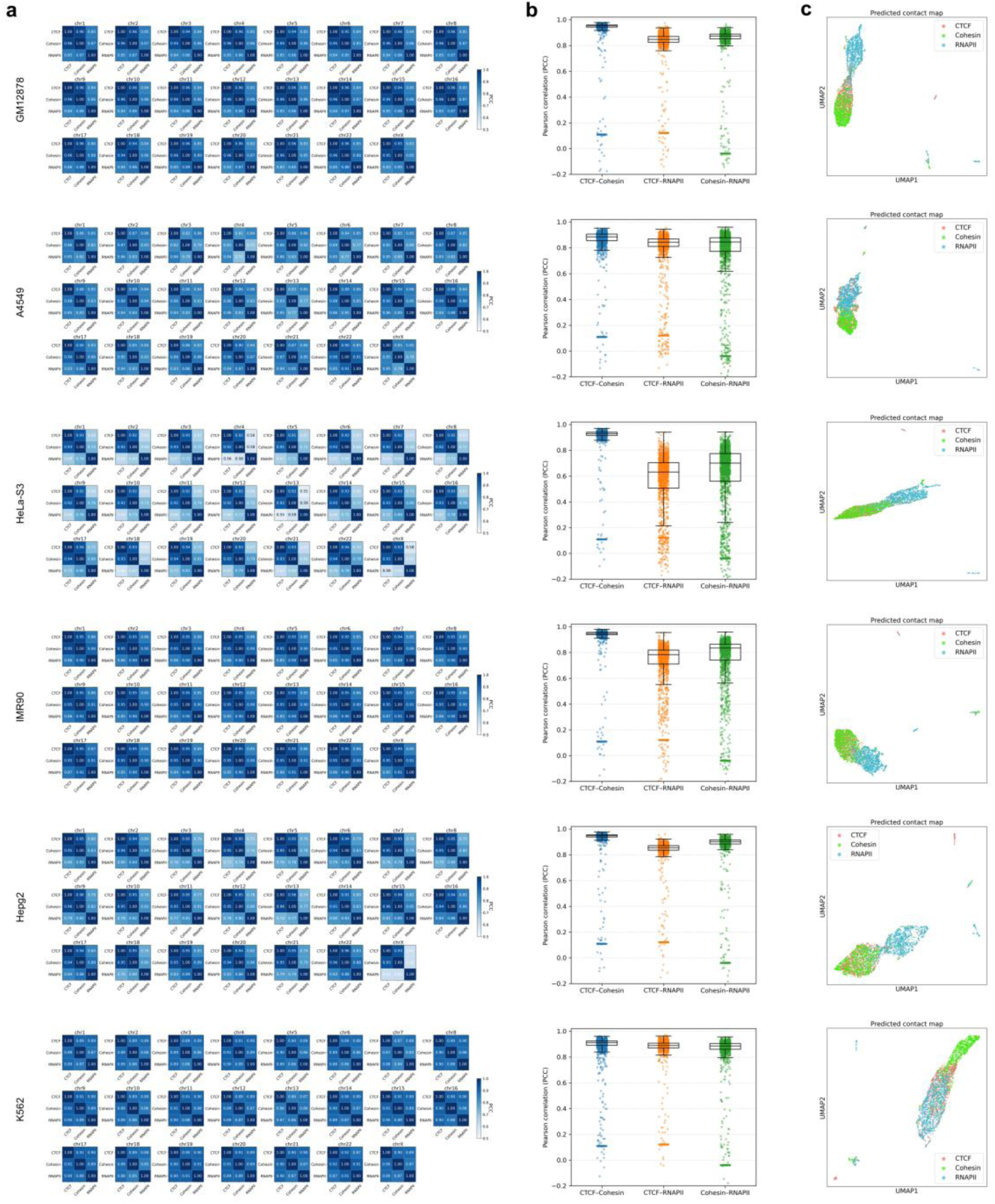
Cross-protein comparison of predicted chromatin architectures across multiple cell types. **(a)** Chromosome-wise Pearson correlation matrices of predicted contact maps between CTCF-, Cohesin-, and RNAPII-mediated interactions across six representative cell types (GM12878, A549, HeLa-S3, IMR90, HepG2, and K562). Consistent with the GM12878 results, CTCF and Cohesin show the strongest pairwise correlations in all cell types, indicating a conserved structural coupling between these two architectural proteins, whereas RNAPII-mediated maps exhibit lower similarity with either. **(b)** Genome-wide pairwise Pearson correlation distributions of predicted contact maps for each protein pair. The consistently higher CTCF-Cohesin correlation compared to CTCF-RNAPII and Cohesin-RNAPII highlights that transcription-associated (RNAPII) interactions form a distinct topological regime from the architectural scaffolds of CTCF/Cohesin. **(c)** UMAP embedding of all predicted contact maps across the genome in each cell type. CTCF and Cohesin clusters show substantial overlap, whereas RNAPII maps form a clearly separated manifold, reinforcing the distinct 3D organization principles of structural versus transcriptional protein-mediated chromatin interactions.

**Extended Data Fig. 9:**
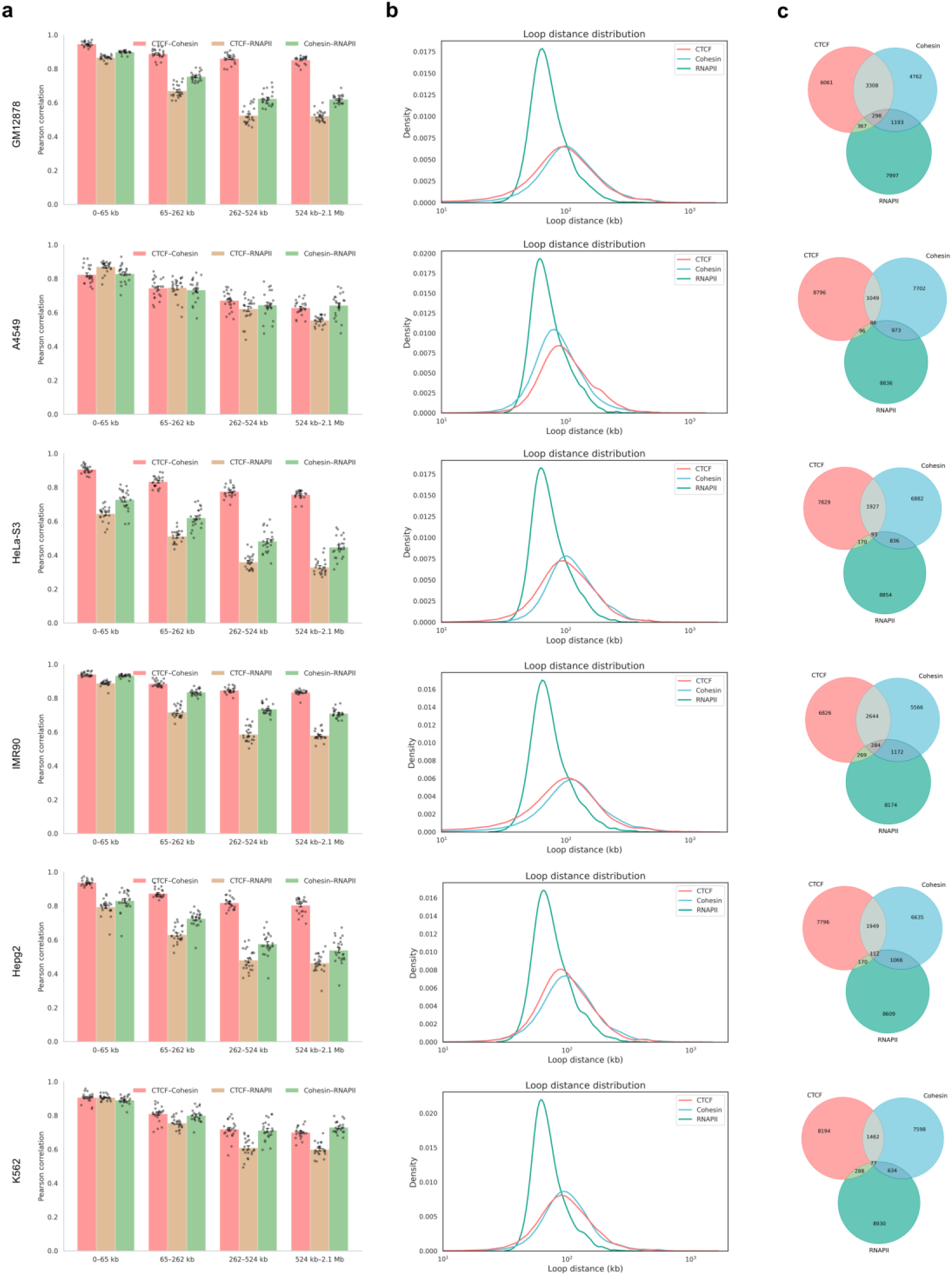
Cross-cell-type comparison of protein-specific chromatin organization patterns. **(a)** Distance-stratified Pearson correlations between predicted contact maps for each pair of proteins (CTCF-Cohesin, CTCF-RNAPII, and Cohesin-RNAPII) across six representative cell types (GM12878, A549, HeLa-S3, IMR90, HepG2, and K562). Consistent across all contexts, CTCF and Cohesin maintain the highest correlation across distance scales, whereas correlations involving RNAPII decline sharply at long genomic distances (>524 kb), indicating its distinct transcription-associated interaction regime. **(b)** Loop distance distributions of the top 10,000 predicted loops for each protein across the six cell types. RNAPII-mediated loops are consistently shorter and more localized, whereas CTCF and Cohesin loops span broader genomic ranges, reflecting their architectural scaffolding roles. **(c)** Venn diagrams showing overlap among the top predicted loops for CTCF, Cohesin, and RNAPII in each cell type. Across all cell types, CTCF and Cohesin share substantial subsets of loops, while RNAPII loops largely occupy distinct regulatory neighborhoods, underscoring protein-specific topological organization principles conserved across the genome.

**Extended Data Fig. 10:**
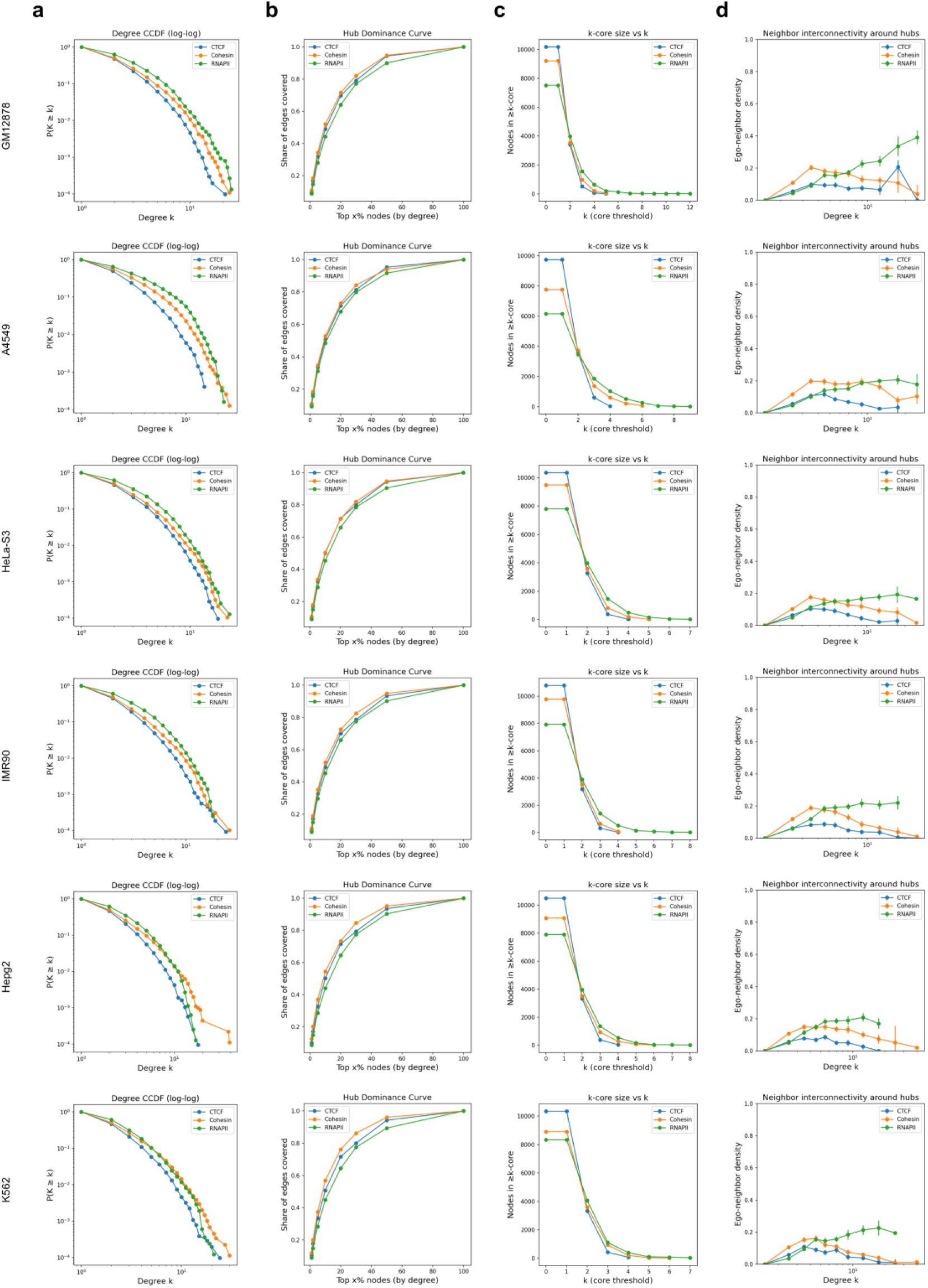
Cross-cell-type network topology of protein-mediated chromatin loops. **(a)** Complementary cumulative degree distributions (CCDF, log-log scale) of loop interaction networks for CTCF-, Cohesin-, and RNAPII-mediated interactions across six representative cell types (GM12878, A549, HeLa-S3, IMR90, HepG2, and K562). All networks display scale-free topologies, with RNAPII showing slightly higher low-degree connectivity, consistent with its dense transcription-associated loops. **(b)** Hub dominance curves showing the cumulative fraction of total loop edges explained by the top-ranked nodes (by degree). Across all cell types, CTCF and Cohesin hubs dominate a larger proportion of total edges than RNAPII, reflecting their roles as structural anchors in chromatin organization. **(c)** k-core decomposition analysis of loop networks. The size of the k-core decreases more gradually for CTCF and Cohesin compared with RNAPII, indicating that architectural loops form more stable hierarchical cores, whereas RNAPII loops are more peripheral and transient. **(d)** Neighbor interconnectivity (ego-neighbor density) around loop hubs across degrees. CTCF and Cohesin display higher local clustering around high-degree nodes, suggesting cooperative domain insulation and compartmentalization, while RNAPII exhibits weaker neighbor interconnectivity, consistent with its dispersed regulatory contacts associated with transcriptional activity.

**Extended Data Fig. 11:**
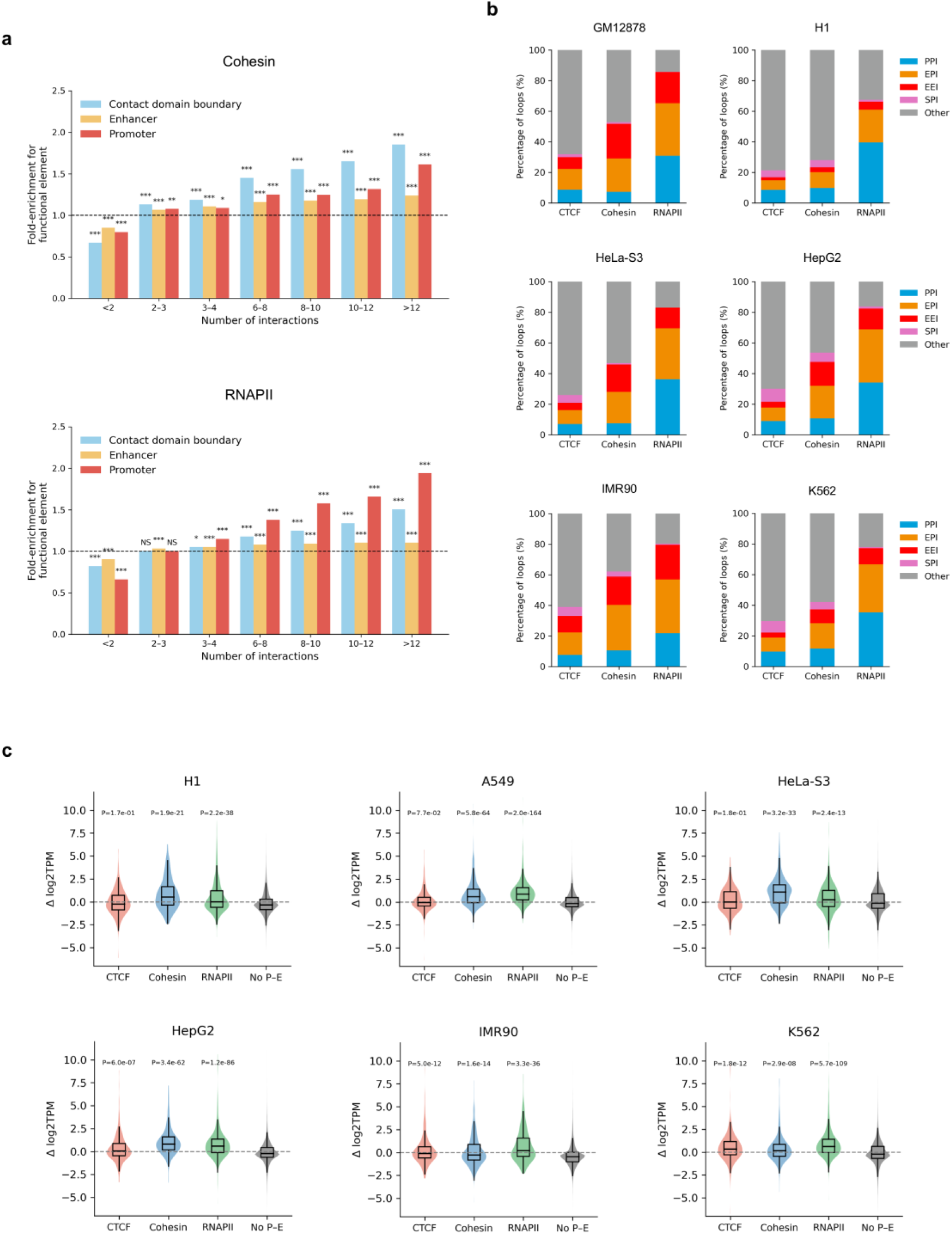
Functional enrichment and regulatory classification of protein-mediated loops across multiple cell types. **(a)** Functional enrichment of anchors from pan-cell-type Cohesin- and RNAPII-mediated loops for contact domain boundaries, enhancers, and promoters, stratified by the number of loop interactions per anchor. Statistical significance was assessed using two-sided Fisher’s exact tests (Number of Cohesin interactions = 48,847, Number of RNAPII interactions = 46,093, \**P<*0.05, \*\**P<*0.01, \*\*\**P<*0.001; *NS*, not significant). **(b)** Distribution of loop categories mediated by CTCF, Cohesin, and RNAPII across six representative cell types (GM12878, H1, HeLa-S3, HepG2, IMR90, and K562). Loops were classified as promoter-promoter (PPI), enhancer-promoter (EPI), enhancer-enhancer (EEI), silencer-promoter (SPI), or other types based on genomic element annotations at loop anchors. **(c)** Differential gene expression associated with genes linked to cell-type-specific loops mediated by CTCF, Cohesin, or RNAPII across the six cell types. Each violin represents the distribution of expression changes for genes engaged in protein-mediated loops compared with genes not participating in promoter-enhancer loops (“No P-E”). Statistical significance was determined using two-sided Wilcoxon rank-sum tests.

**Extended Data Fig. 12:**
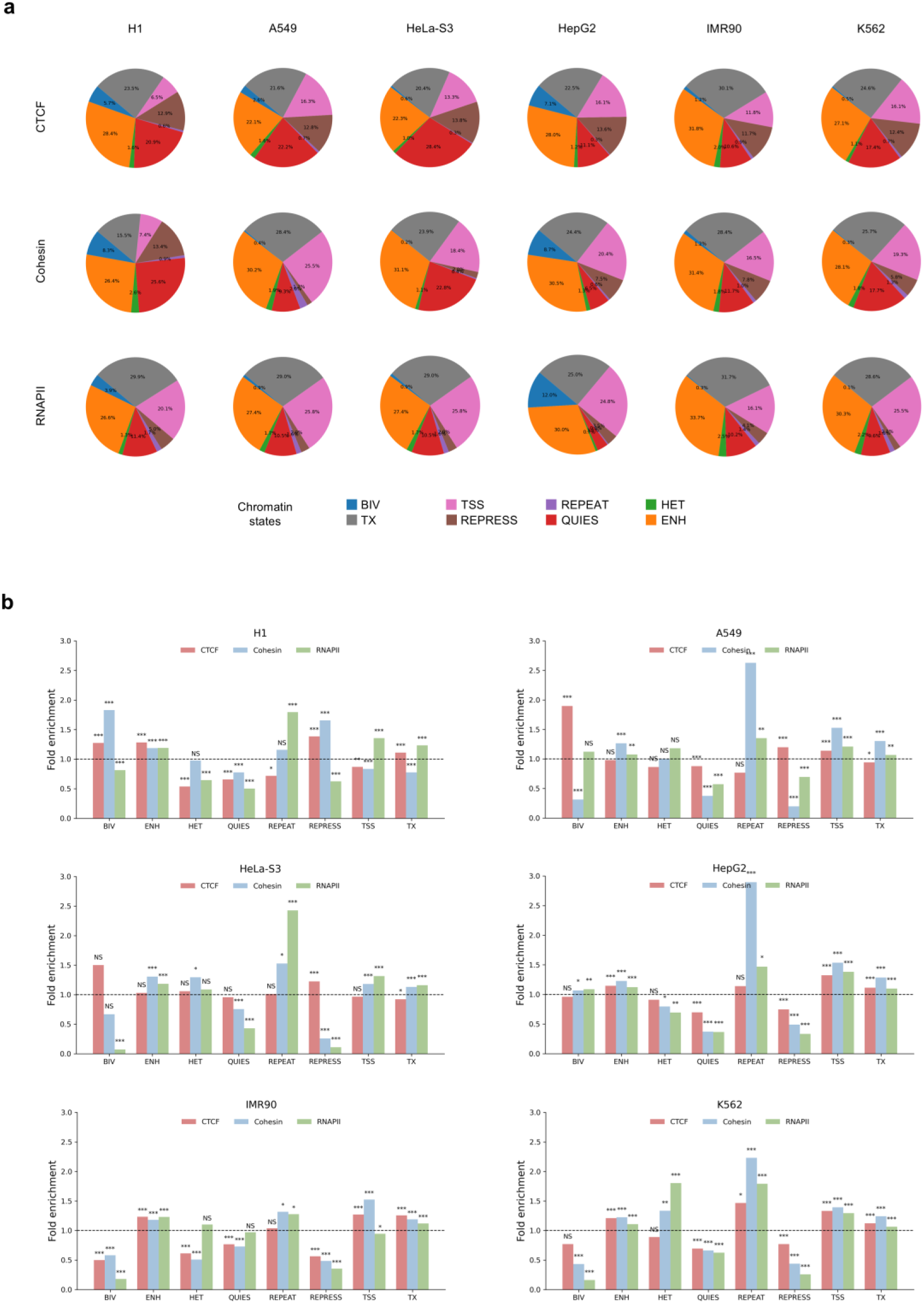
Chromatin-state composition and enrichment of protein-specific loop anchors across additional cell types. **(a)** Chromatin-state composition of cell-type-specific loop anchors mediated by CTCF, Cohesin, and RNAPII across six representative human cell types (H1, A549, HeLa-S3, HepG2, IMR90, and K562). Each pie chart shows the proportion of chromatin states. **(b)** Fold enrichment of chromatin states at cell-type-specific loop anchors for CTCF-, Cohesin-, and RNAPII-mediated loops in the same six cell types. Statistical significance was evaluated using two-sided Fisher’s exact tests (\**P<*0.05, \*\**P<*0.01, \*\*\**P<*0.001; *NS*, not significant).

**Extended Data Fig. 13:**
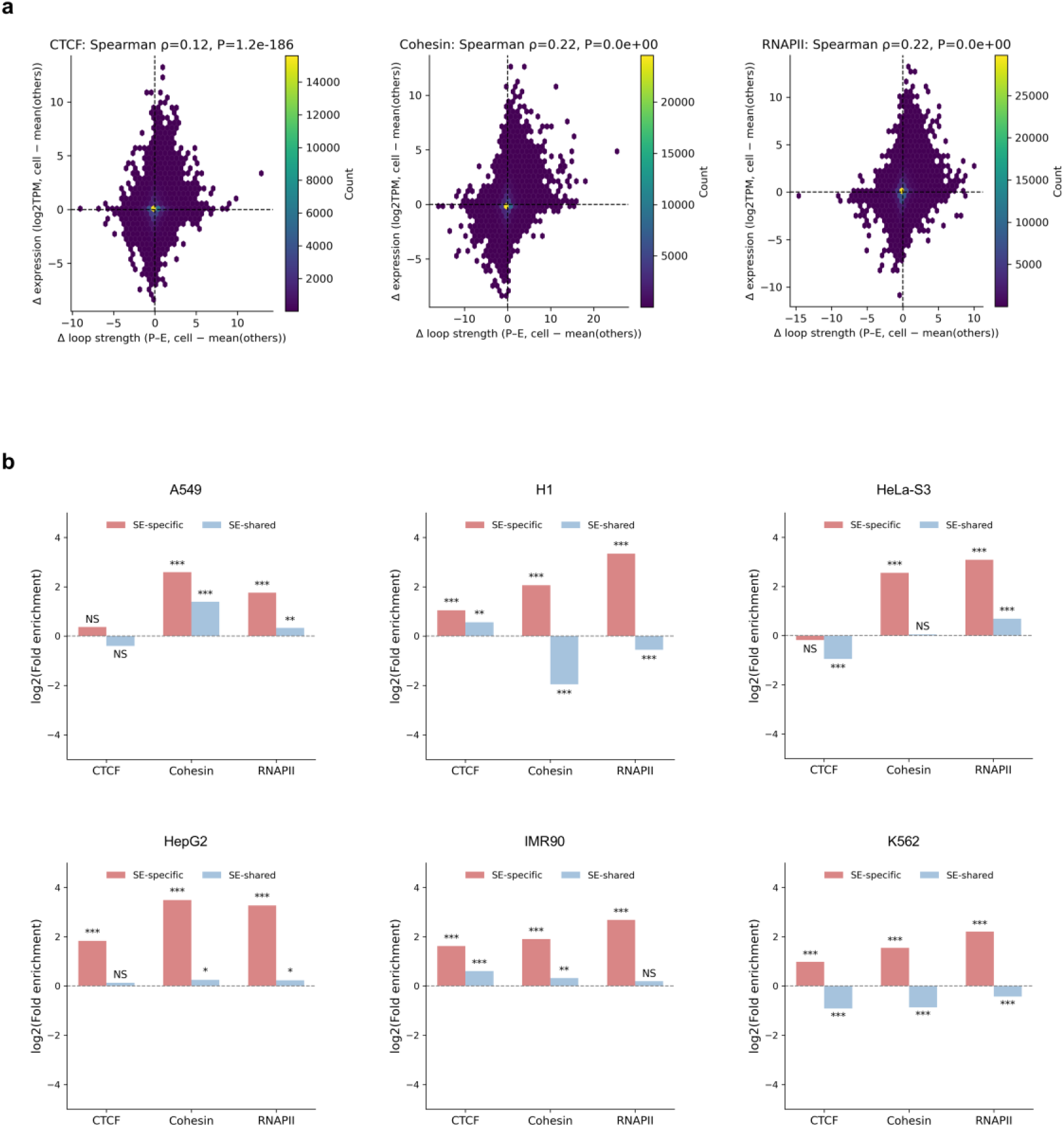
Functional coupling between loop strength, gene expression, and super-enhancer association across multiple cell types. **(a)** Relationship between predicted loop strength and the corresponding gene expression change for CTCF-, Cohesin-, and RNAPII-mediated loops. Each point represents a promoter-enhancer pair, with color indicating density. **(b)** Fold enrichment of cell-type-specific versus non-specific loops in cell-type-specific or shared super-enhancers (SEs) across six representative cell types (A549, H1, HeLa-S3, HepG2, IMR90, and K562). SE-specific loops were significantly enriched around cell-type-specific SEs, while shared SEs were more frequently linked by common loops. Statistical significance was assessed using two-sided Fisher’s exact tests (\**P<*0.05, \*\**P<*0.01, \*\*\**P<*0.001; *NS*, not significant).

**Extended Data Fig. 14:**
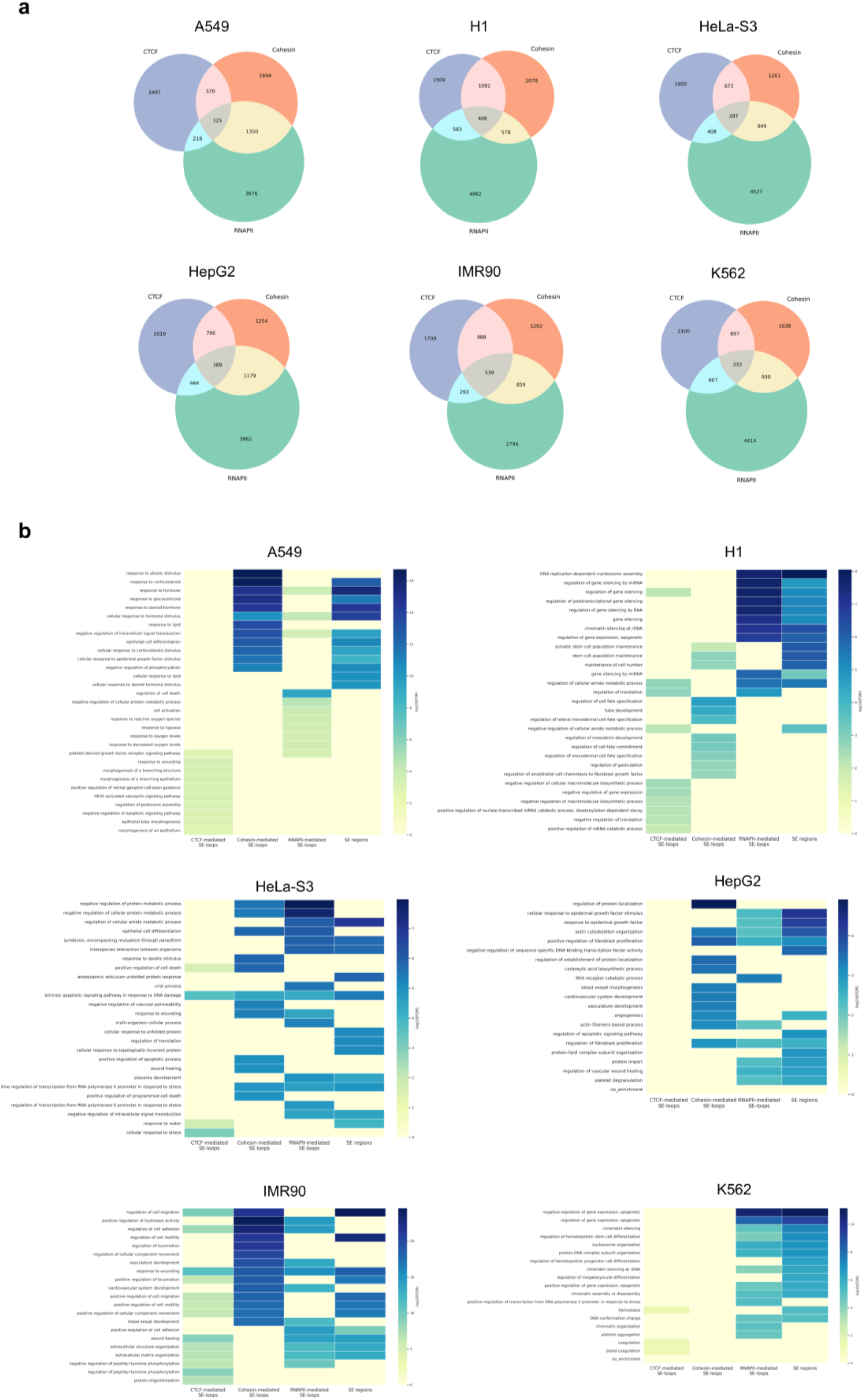
Overlap and canonical pathway enrichment of genes associated with protein-mediated SE-linked loops across multiple cell types. **(a) Overlap of genes associated** with CTCF-, Cohesin-, and RNAPII-mediated loops across six representative human cell types (A549, H1, HeLa-S3, HepG2, IMR90, and K562). Venn diagrams depict the intersection among genes linked to loops mediated by each protein, highlighting both shared and protein-specific regulatory targets. **(b)** Canonical pathway enrichment analysis of genes associated with super-enhancer (SE)-linked loops mediated by CTCF, Cohesin, and RNAPII in the same six cell types. The top enriched pathways are shown for each protein-specific SE-loop set as well as SE regions alone, revealing both common and protein-distinct biological processes.

**Extended Data Fig. 15:**
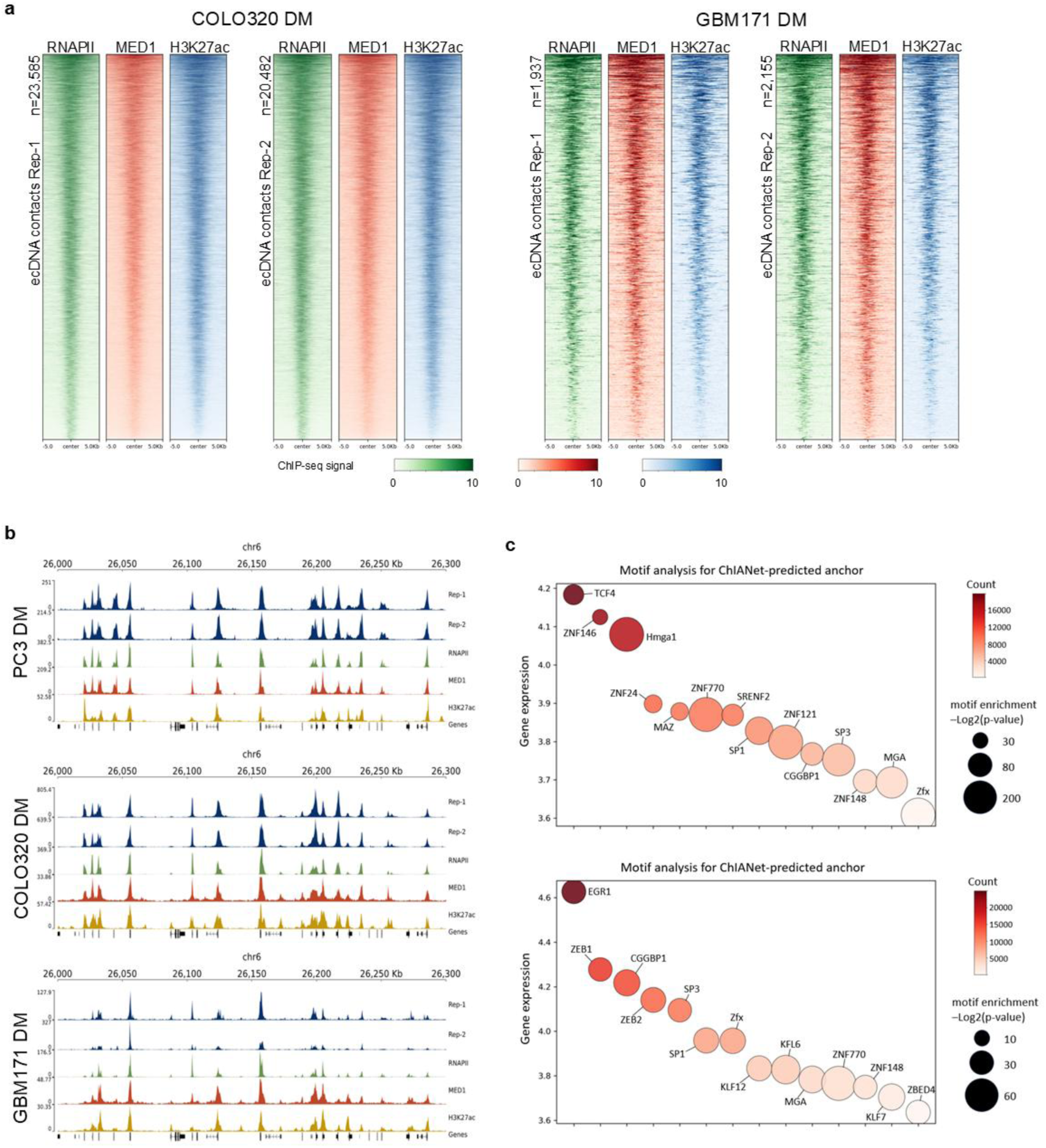
Epigenomic and motif features of ecDNA-associated RNAPII interactions across cancer cell types. **(a)** Heatmaps showing ChIP-seq signal densities of RNAPII, MED1 and H3K27ac centered on ecDNA-associated contact anchors in COLO320DM and GBM171DM cells. Results from two biological replicates are shown for each cell type (COLO320DM Rep-1, n = 23,585; Rep-2, n = 20,482; GBM171DM Rep-1, n = 1,937; Rep-2, n = 2,155). Signals are displayed within ±5 kb of anchor centers. Color scales indicate normalized ChIP-seq signal intensities. **(b)** Genome browser views of representative ecDNA-associated regions in PC3DM, COLO320DM and GBM171DM cells. Tracks show ChIA-drop interaction signals from two biological replicates, RNAPII, MED1 and H3K27ac ChIP-seq signals, and gene annotations. **(c)** Motif analysis of ChIANet-predicted RNAPII loop anchors in COLO320DM (top) and GBM171DM (bottom) cells. Each point represents a transcription factor motif. Point size denotes motif enrichment significance (-log2 P value), and color intensity indicates RNA-seq-derived gene expression levels (raw expression counts) of the corresponding transcription factors. The y-axis shows gene expression levels.

**Extended Data Fig. 16:**
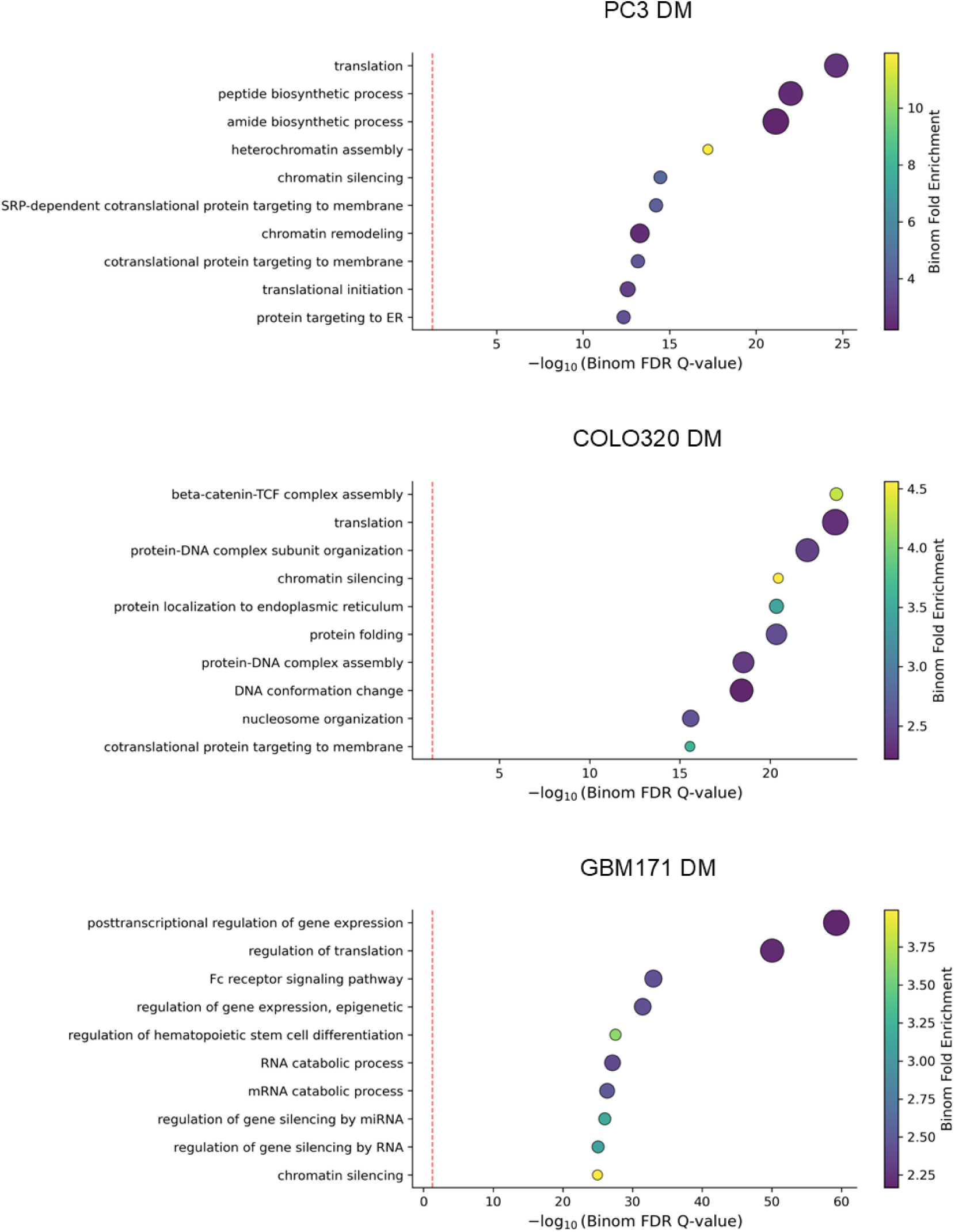
Gene Ontology enrichment analysis of ChIANet-predicted loop anchors across cancer cell types. Gene Ontology (GO) biological process enrichment analysis of ChIANet-predicted chromatin loop anchors in PC3DM (top), COLO320DM (middle) and GBM171DM (bottom) cells. Each point represents an enriched GO term. The x-axis shows -log10-transformed binomial FDR Q values. Point color indicates binomial fold enrichment, and point size reflects the number of genes associated with each GO term. The red dashed line denotes the significance threshold used for enrichment analysis.

