## Supplementary Information for "Deep learning framework ChIANet predicts protein-mediated chromatin architecture across functional contexts"

| Protein | Cell type | ChIP-seq | ChIA-PET | Note |
|---|---|---|---|---|
| CTCF | GM12878 | ENCFF453EJM<br>ENCFF355CYX | ENCSR184YZV | Train/Val/Test |
| CTCF | H1 | ENCFF767YLK | ENCSR095SXC | *De novo* |
| CTCF | A549 | ENCFF560QWU<br>ENCFF813GIN | | *De novo* |
| CTCF | HeLa-S3 | ENCFF898VGF<br>ENCFF586MXO | | *De novo* |
| CTCF | HepG2 | ENCFF012FMD<br>ENCFF487UUI | | *De novo* |
| CTCF | IMR90 | ENCFF584CBK<br>ENCFF160QTH | | *De novo* |
| CTCF | K562 | ENCFF172KOJ<br>ENCFF265ZSP | | *De novo* |
| RNAPII | GM12878 | ENCFF501USI<br>ENCFF886CYK | ENCSR905HWW | Train/Val/Test |
| RNAPII | H1 | ENCFF102CMM<br>ENCFF069GAM | | *De novo* |
| RNAPII | A549 | ENCFF816DKP<br>ENCFF641ZJE | | *De novo* |
| RNAPII | HeLa-S3 | ENCFF390YVA<br>ENCFF311UCL | | *De novo* |
| RNAPII | HepG2 | ENCFF835GBL<br>ENCFF845YGC | | *De novo* |
| RNAPII | IMR90 | ENCFF740GRA<br>ENCFF143IJT | | *De novo* |
| RNAPII | K562 | ENCFF201SIE<br>ENCFF267TTN | | *De novo* |
| RAD21 (Cohesin) | GM12878 | ENCFF873QGF<br>ENCFF638BXR | 4DNESHSMKNGT | Train/Val/Test |
| RAD21 (Cohesin) | H1 | ENCFF100EES<br>ENCFF200HXK | | *De novo* |
| RAD21 (Cohesin) | A549 | ENCFF341CJU<br>ENCFF636BLH | | *De novo* |
| RAD21 (Cohesin) | HeLa-S3 | ENCFF632OBR<br>ENCFF705SCW | | *De novo* |
| RAD21 (Cohesin) | HepG2 | ENCFF319BXY<br>ENCFF221VFQ | | *De novo* |
| RAD21 (Cohesin) | IMR90 | ENCFF918IXN<br>ENCFF824JOL | | *De novo* |

| | | | | |
|---|---|---|---|---|
| RAD21 (Cohesin) | K562 | ENCFF985ZJW<br>ENCFF150ILI | | *De novo* |

**Supplementary Table 1. ChIP-seq and ChIA-PET data used for training, validation and testing.**

| **Protein** | **Cell type** | **Raw** | **Quality Controled** |
|---|---|---|---|
| CTCF | GM12878 | 1181427 | 152338 |
| CTCF | H1 | 2111961 | 212573 |
| RNAPII | GM12878 | 450480 | 66833 |
| Cohesin | GM12878 | 2019473 | 276093 |

**Supplementary Table 2. Summary of ChIA–PET loop counts before and after quality control.**

| **Cell type** | **H3k27ac** | **H3k27me3** | **RNA-seq** | **Hi-C** |
|---|---|---|---|---|
| GM12878 | ENCSR000AKC | ENCSR000AKD | ENCSR843RJV | ENCSR968KAY |
| H1 | ENCSR880SUY | ENCSR216OGD | ENCSR043RSE | - |
| A549 | ENCSR778NQS | - | ENCSR937WIG | ENCSR662QKG |
| HeLa-S3 | ENCSR000AOC | ENCSR000APB | ENCSR000CPR | - |
| HepG2 | ENCSR000AMO | ENCSR000AOL | ENCSR985KAT | ENCSR194SRI |
| IMR90 | ENCSR002YRE | ENCSR431UUY | ENCSR000CTQ | ENCSR852KQC |
| K562 | ENCSR000AKP | ENCSR000AKQ | ENCSR000AEM | ENCSR000AEM |

**Supplementary Table 3. Overview of H3k27ac, H3k27me3, RNA-seq and Hi-C datasets used in this study.**

| Cell type | REMC ID |
| --- | --- |
| GM12878 | E003 |
| H1 | E017 |
| A549 | E114 |
| HeLa-S3 | E116 |
| HepG2 | E117 |
| IMR90 | E118 |
| K562 | E123 |

**Supplementary Table 4. Epigenome Roadmap Project cell line identifiers.**

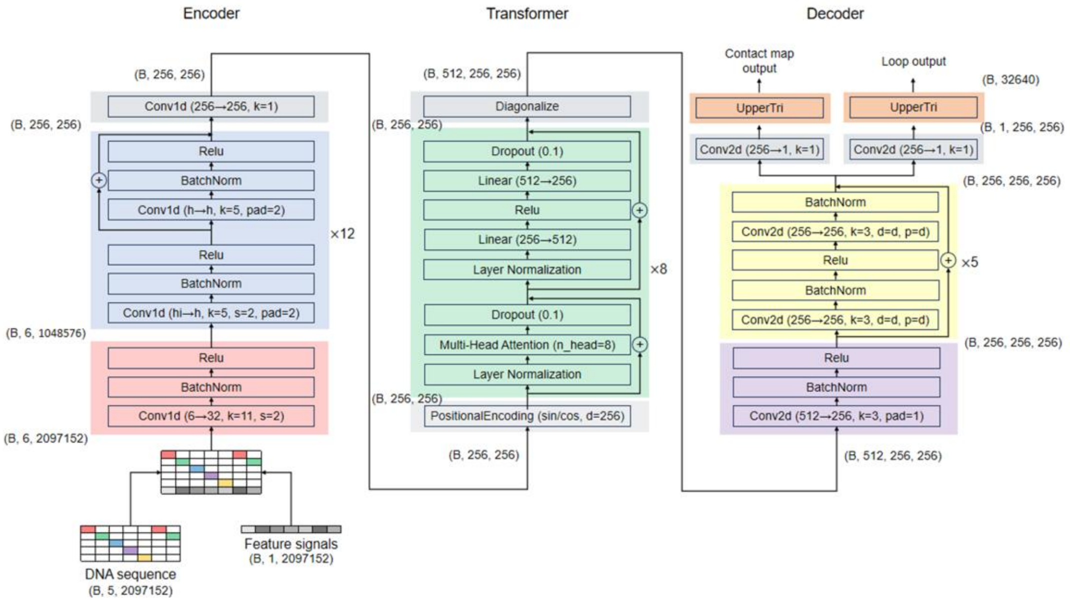


**Supplementary Figure 1. ChIANet model structure and module components.** A schematic of the ChIANet architecture illustrating the encoder, Transformer and decoder modules. The model takes a 2.1-Mb genomic window as input, combining a one-hot–encoded DNA sequence and protein-specific ChIP-seq signals. The encoder consists of twelve 1D convolutional residual blocks that progressively extract local sequence–feature representations and downscale inputs by strided convolutions (stride = 2). Each block contains Batch Normalization and ReLU activations to stabilize deep-layer training. The Transformer module includes eight self-attention blocks, each composed of multi-head attention, feed-forward layers, dropout and layer normalization, with positional encoding to capture long-range dependencies across 256 bins. The decoder contains five stacked 2D convolutional residual blocks that transform the latent representation into two outputs: a contact map and a loop matrix, both at 8,192-bp resolution. Dilated convolutions are used to enlarge the receptive field and ensure that each pixel in the output corresponds to the full genomic context within the 2.1-Mb window.

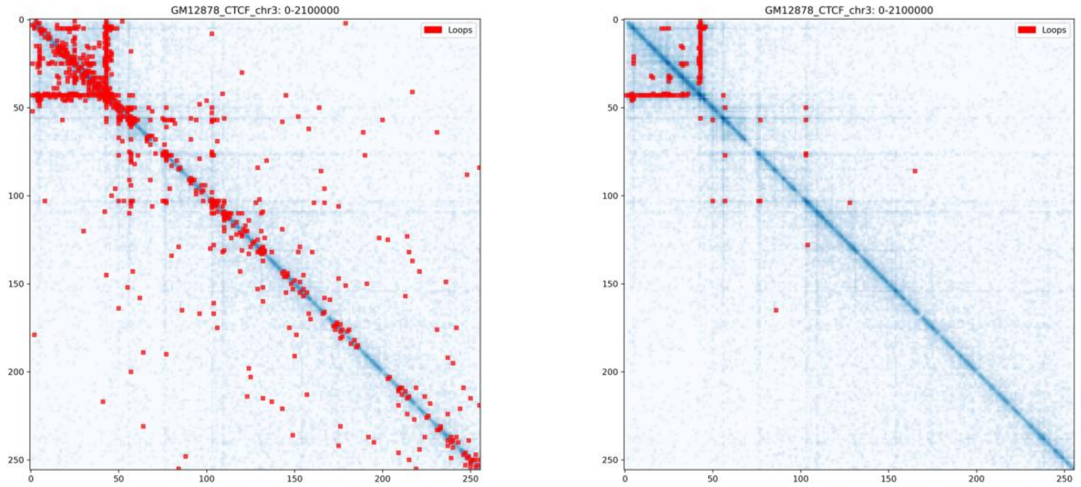


**Supplementary Figure. 2. Example of ChIA-PET loop quality control before and after filtering.** Shown are 10-kb CTCF ChIA-PET contact maps from GM12878 with loop calls overlaid (red squares) for a representative 2.1-Mb region on chromosome 3; left, raw loops output by ChIA-PIPE; right, high-confidence intra-chromosomal loops retained.

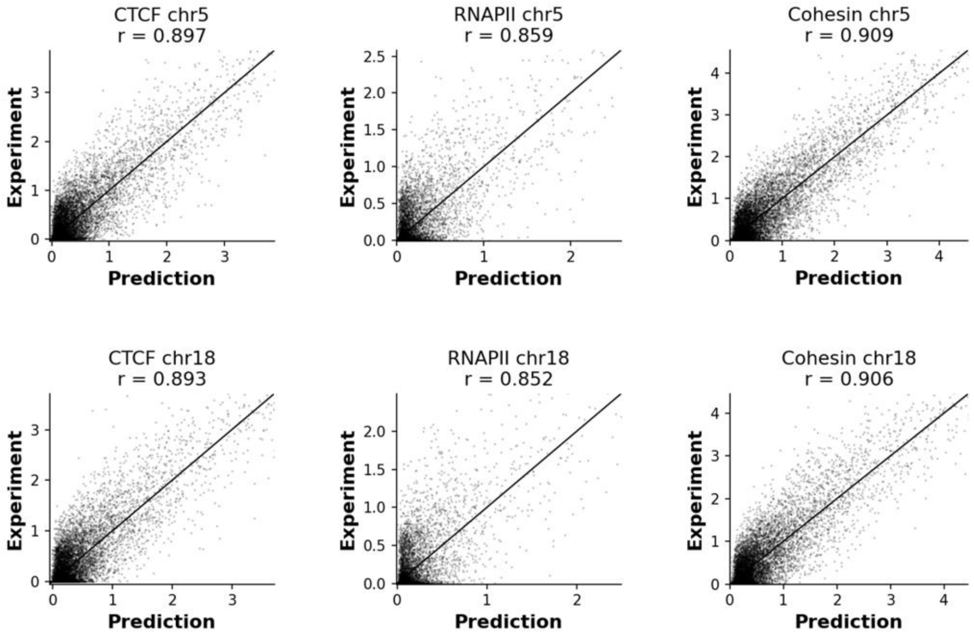


**Supplementary Figure. 3. Chromosome-scale prediction accuracy of ChIANet across proteins.** Scatter plots showing the correlation between predicted and experimental contact map values for CTCF-, RNAPII-, and Cohesin-mediated interactions on chromosomes 5 and 18 (test set). For each protein, 10,000 subsampled contact values were randomly selected from the contact map. Pearson correlation coefficients (r) are indicated in each panel. ChIANet exhibits strong concordance with experimental ChIA-PET maps across proteins and chromosomes, demonstrating consistent prediction fidelity and accurate recovery of quantitative contact intensities at the chromosome scale.

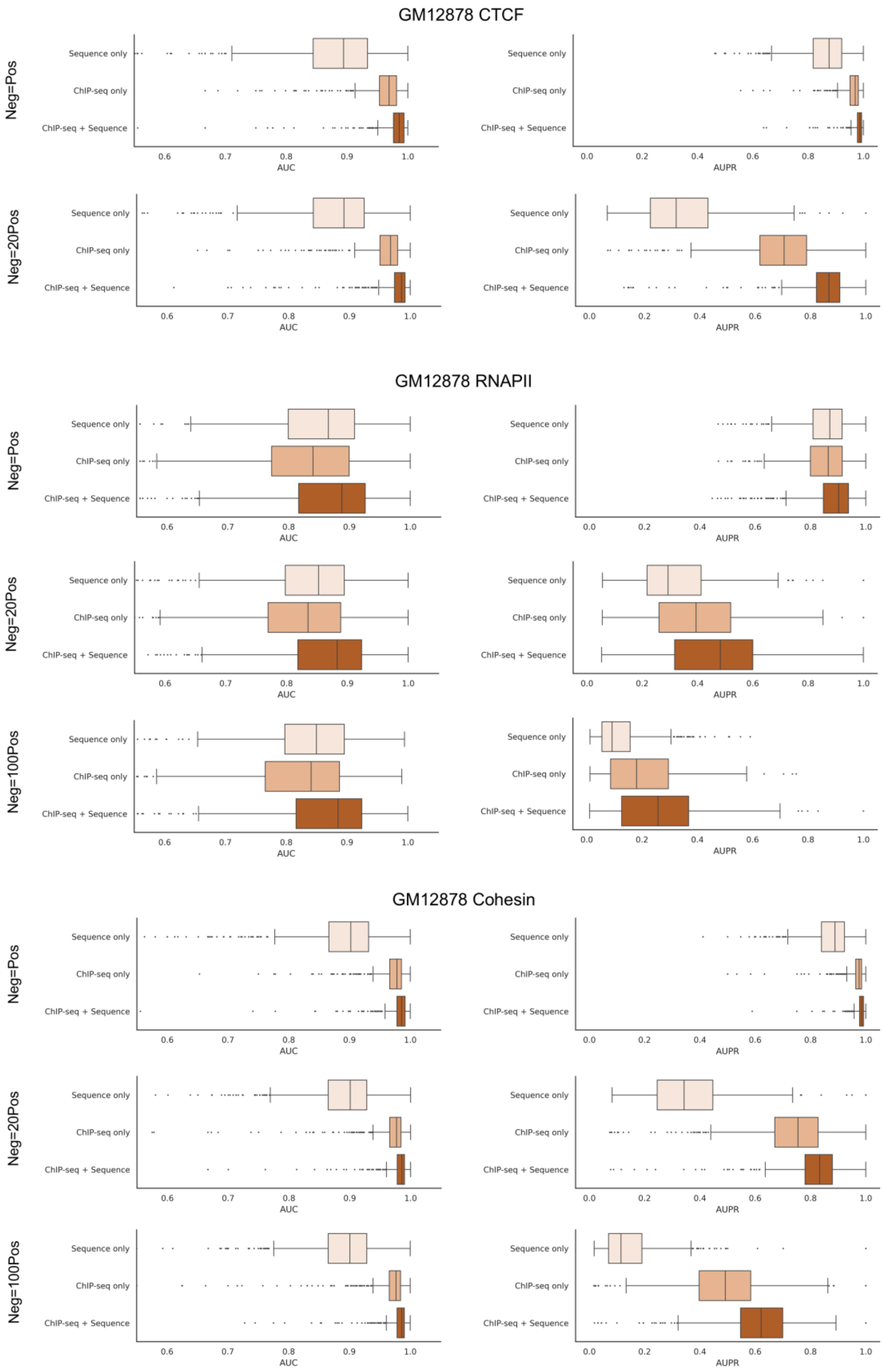


**Supplementary Figure. 4. Comparison of ChIANet ablation models under different class imbalance settings.** Comparison of the sequence only, ChIP-seq only, and sequence + ChIP-seq (full) ChIANet models for CTCF, RNAPII, and Cohesin on GM12878. Each panel shows window-level area under the ROC curve (AUC) (left) and area under the precision–recall curve (AUPR) (right) for loop prediction under three positive-to-negative sampling ratios: 1:1, 1:20, and 1:100. Across all imbalance settings, the full model integrating both modalities consistently outperforms single-modality variants, maintaining higher classification accuracy (AUC) and superior precision-recall balance (AUPR).

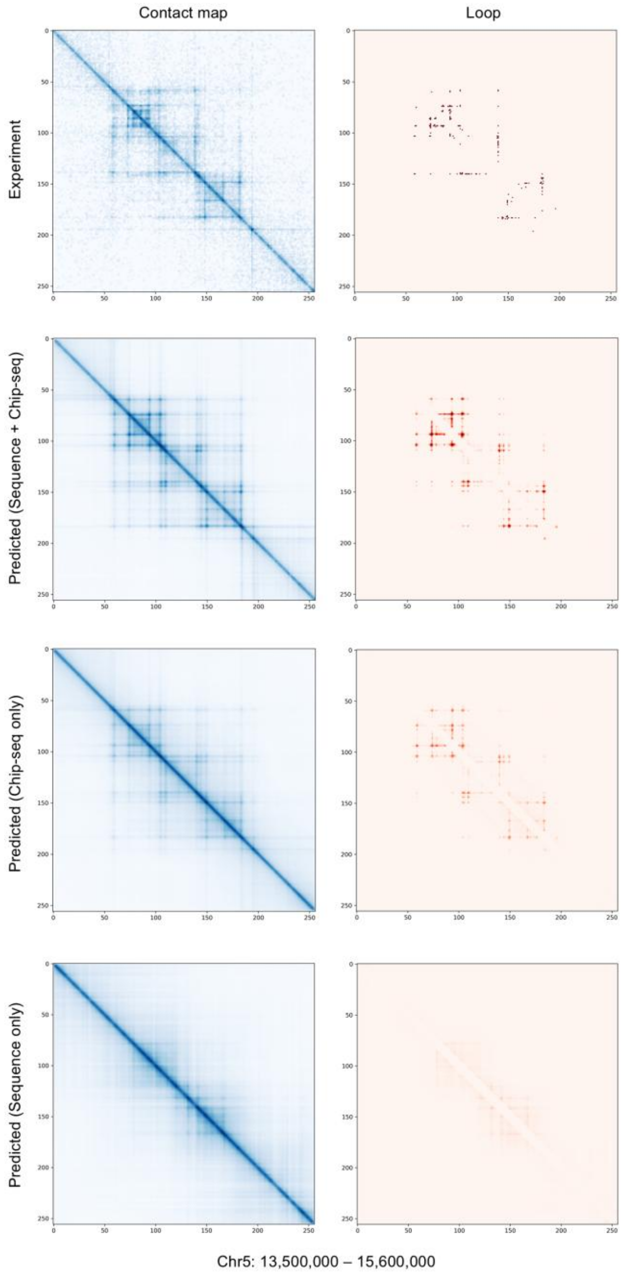


**Supplementary Figure. 5. Examples of ablation analysis.** The full model achieves the highest precision-recall performance among all input configurations. Representative examples of predicted contact maps and loop matrices for the full, ChIP-seq only, and sequence only models compared with experimental ChIA-PET data on chromosome 5, illustrating that both modalities are required to recapitulate fine-scale contact structures and loop patterns.

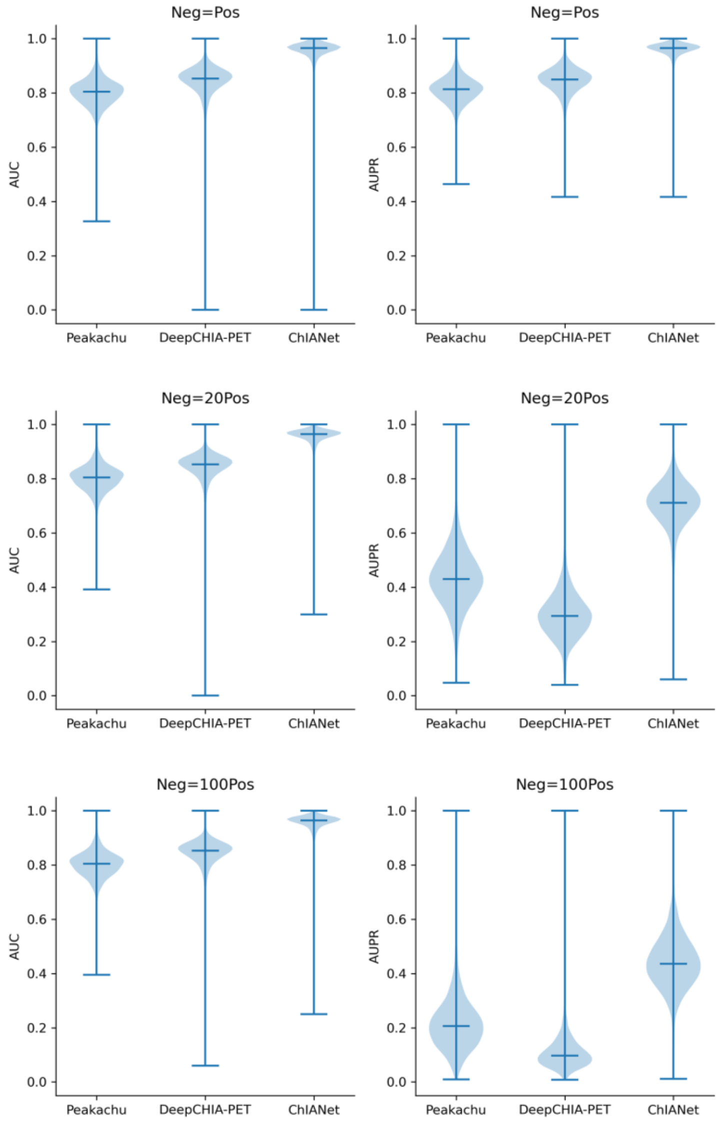


**Supplementary Figure. 6. Evaluation of de novo loop prediction under varying positive-to-negative ratios.** Comparison of ChIANet with baseline models (Peakachu and DeepChIA-PET) across different class-imbalance settings (1:1, 1:20, and 1:100). Each violin plot summarizes model performance in terms of AUC (left) and AUPR (right) for all genomic windows. ChIANet consistently achieves the highest precision–recall performance across increasingly imbalanced conditions, indicating its robustness to sparse positive loop annotations. Unlike the Hi-C-dependent baselines, ChIANet requires only DNA sequence and protein-specific ChIP-seq input, demonstrating strong generalization in *de novo* loop prediction.

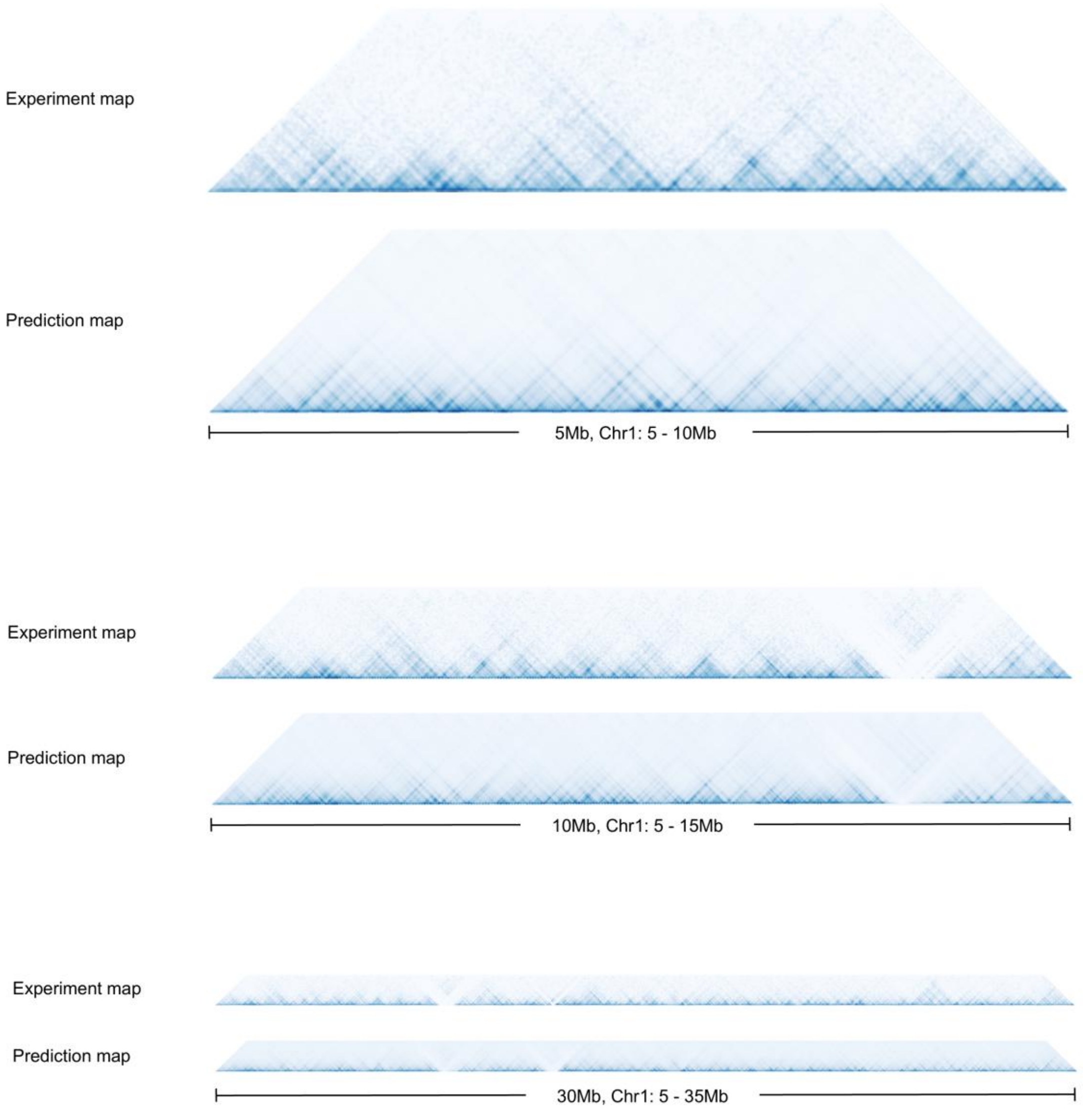


**Supplementary Figure. 7. Reconstruction of large-scale contact maps from tiled ChIANet predictions.** Representative examples showing that local 2.1-Mb contact map predictions generated by ChIANet can be seamlessly merged into larger continuous chromosomal contact maps. Each panel compares the experimentally measure ChIA-PET contact maps (top) with the corresponding ChIANet-predicted maps (bottom) across genomic regions of increasing length—5 Mb (Chr1: 5-10 Mb), 10 Mb (Chr1: 5-15 Mb), and 30 Mb (Chr1: 5-35 Mb). The predicted maps preserve multi-scale topological domain structures and long-range interaction continuity, indicating that ChIANet maintains structural coherence when extended from window-level predictions to chromosome-scale reconstructions.

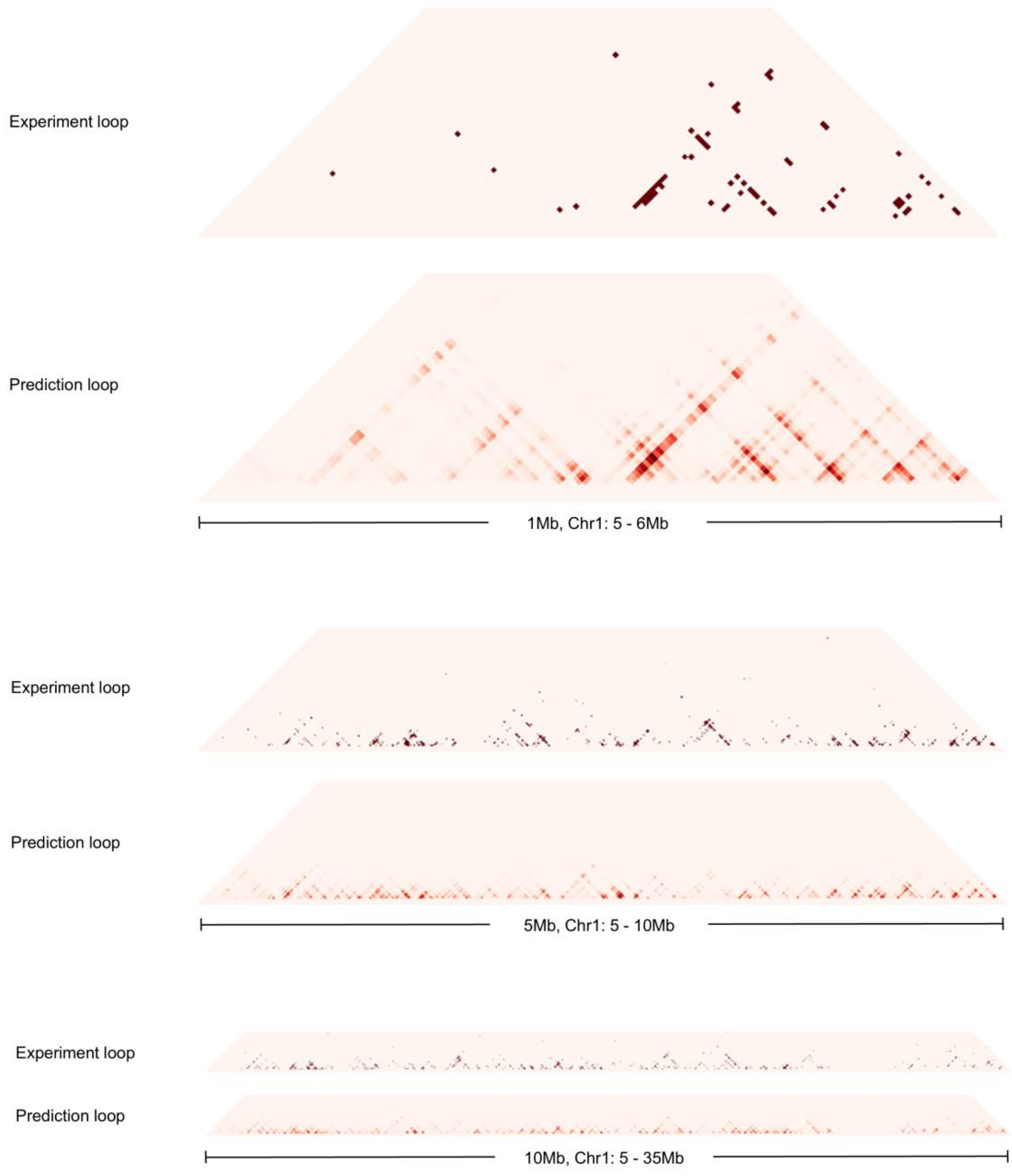


**Supplementary Figure. 8. Reconstruction of large-scale loop maps from tiled ChIANet predictions.** Representative examples showing that local 2.1-Mb loop predictions generated by ChIANet can be seamlessly merged to produce larger chromosomal loop maps. Each panel compares experimentally measured ChIA-PET loops (top) with ChIANet-predicted loops (bottom) across increasing genomic scales—1 Mb (Chr1: 5-6 Mb), 5 Mb (Chr1: 5-10 Mb), and 10 Mb (Chr1: 5-15 Mb). The merged predictions preserve high-confidence focal loops and domain-level interaction patterns across extended genomic regions, demonstrating that ChIANet maintains both local precision and long-range structural continuity when scaled from window-level to chromosome-scale loop reconstruction.

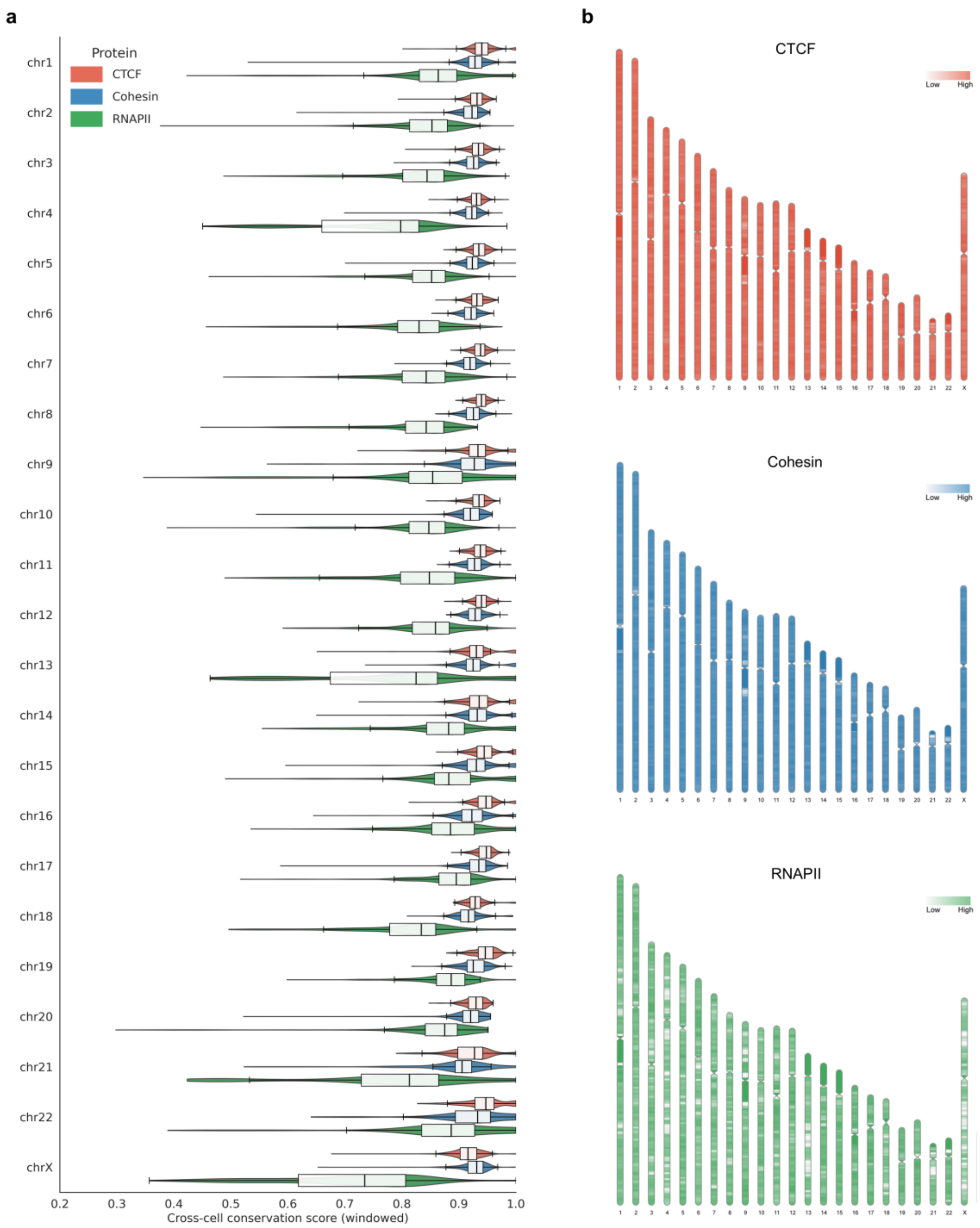


**Supplementary Figure. 9. Chromosome-wise conservation of protein-mediated contact maps across cell types. (a)** Cross-cell-type mean Pearson correlation coefficients (PCCs) of predicted contact maps for CTCF (red), Cohesin (blue), and RNAPII (green) across 23 human chromosomes. Each violin-and-box plot represents the distribution of conservation scores across all 2.1-Mb genomic windows within a chromosome. **(b)** Genome-wide visualization of contact map conservation scores for each protein across the 23 chromosomes. Each vertical bar represents a chromosome divided into 2.1-Mb segments, colored according to the mean cross-cell PCC within each segment (low to high conservation).

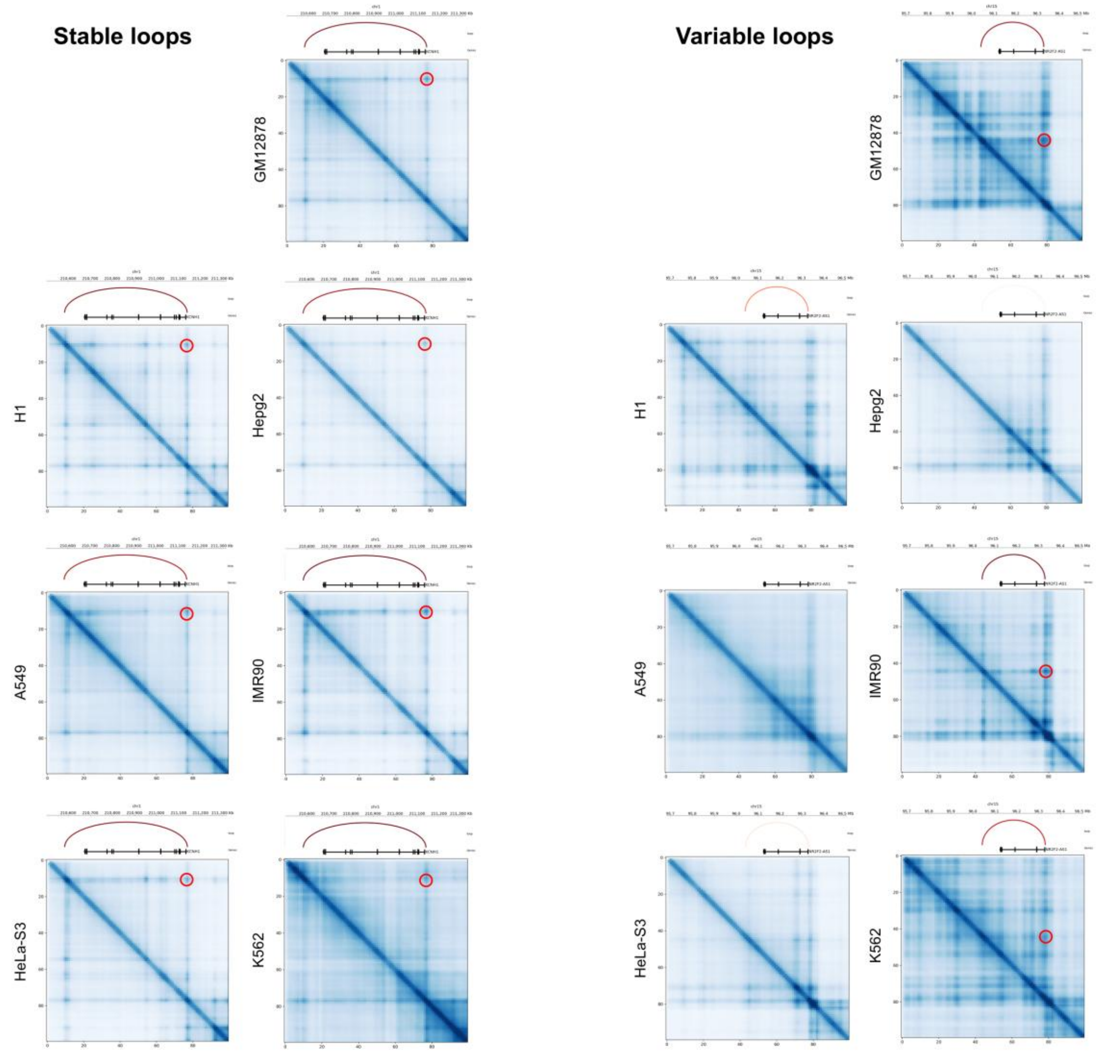


**Supplementary Figure. 10. Examples of stable and variable loops across cell types.** Representative contact maps showing CTCF-mediated loops classified as stable (left) or variable (right) across seven human cell types (GM12878, H1, HepG2, A549, IMR90, HeLa-S3, and K562). Stable loops are consistently observed across all cell types, whereas variable loops appear in only a subset of cell types. Red arcs mark loop positions, and corresponding anchors are indicated above each contact map.

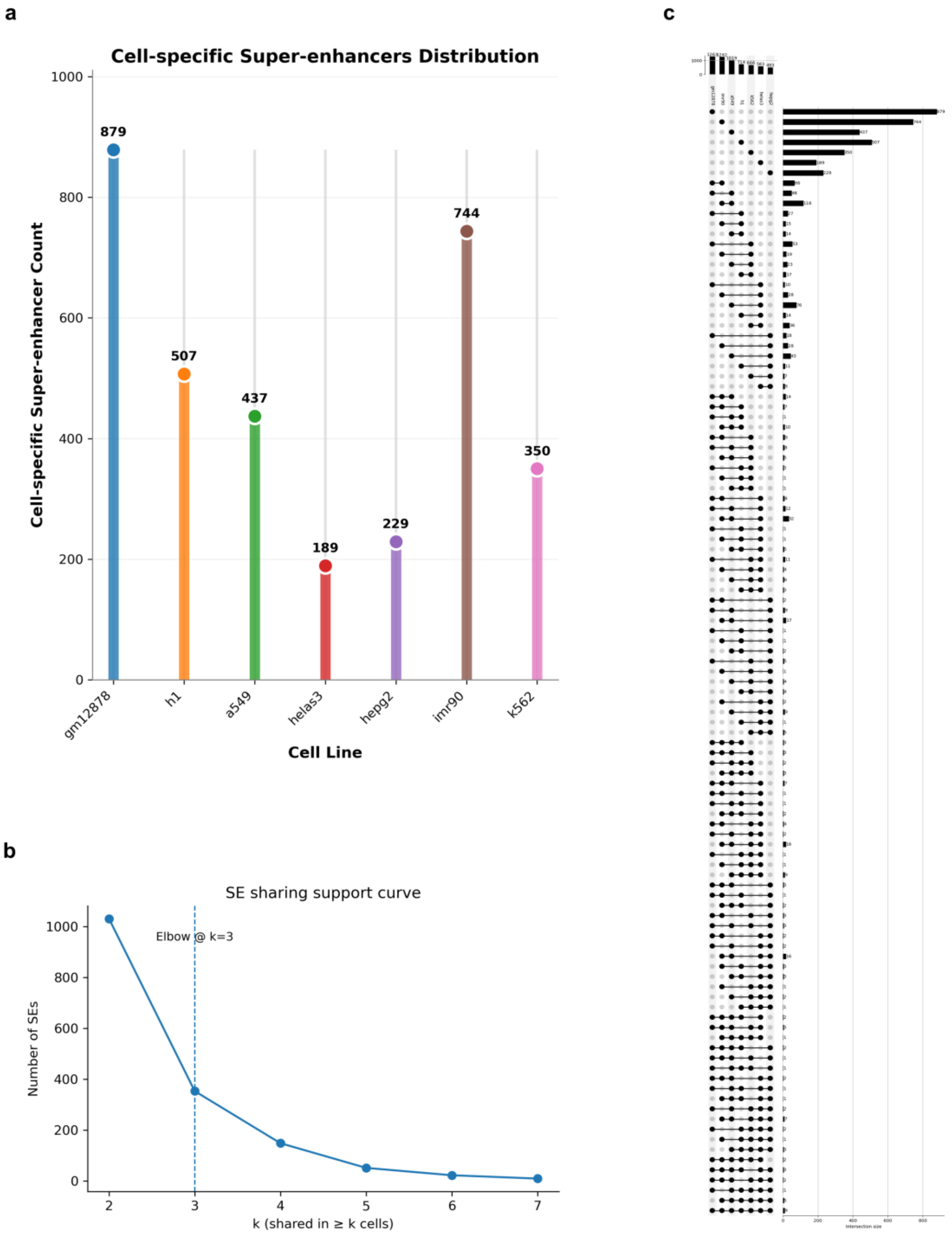


**Supplementary Figure. 11. Characterization of super-enhancer (SE) distribution and sharing across cell types. (a)** Number of cell-type-specific super-enhancers identified in each of the seven human cell lines (GM12878, H1, A549, HeLa-S3, HepG2, IMR90, and K562). GM12878 and HepG2 exhibit the largest numbers of unique SEs, consistent with strong lineage-specific enhancer activity. **(b)** SE sharing support curve showing the relationship between the number of shared SEs and the number of supporting cell types (k). The elbow point at k = 3 defines the threshold used to distinguish cell-type-specific SEs (k < 3) from shared SEs (k ≥ 3). **(c)** UpSet plot displaying the combinatorial overlap patterns of SEs across the seven cell types. The horizontal bars represent the total SE count per cell type, while the vertical bars indicate the number of SEs shared by the corresponding cell-type combinations.

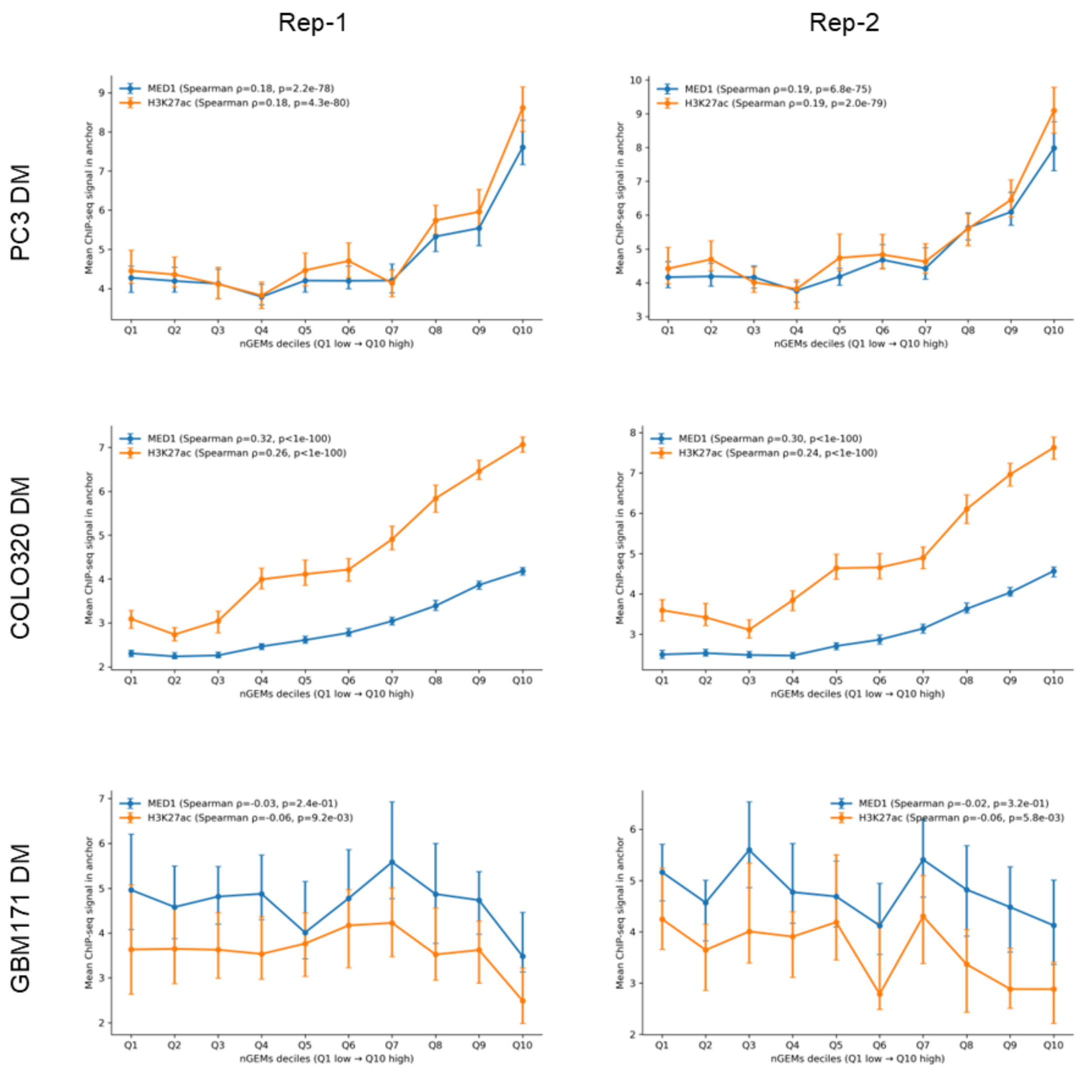


**Supplementary Figure. 12. Association between ChIA-Drop nGEMs and transcription-associated ChIP-seq signals across cancer cell types.** Mean ChIP-seq signal intensities of MED1 (blue) and H3K27ac (orange) at ChIA-Drop nGEM anchors across deciles of nGEM interaction strength in PC3DM (top), COLO320DM (middle) and GBM171DM (bottom) cells. NGEMs were ranked by interaction strength and grouped into deciles from lowest (Q1) to highest (Q10). Results are shown for two biological replicates (Rep-1, left; Rep-2, right). Points represent mean ChIP-seq signal values within anchor regions, and error bars indicate s.e.m. Spearman correlation coefficients (ρ) and corresponding P values between nGEM deciles and ChIP-seq signal intensities are indicated in each panel.
